# Structural basis and mechanism of action of NrdR, a bacterial master regulator of ribonucleotide reduction

**DOI:** 10.1101/2025.03.28.645925

**Authors:** Lucas Pedraz, Arkadiusz Szura, Claus Schmitz, Alba Rubio-Canalejas, Anthony Santella, Gabriel Gomila, Annalisa Calo, Maria Solà, Eduard Torrents

## Abstract

Ribonucleotide reductases (RNRs) are the essential enzymes responsible for synthesizing dNTPs, the building blocks of DNA. In bacteria, the entire RNR network is controlled by the master regulator NrdR. As a regulator of an essential pathway with no eukaryotic equivalent, NrdR is a promising antimicrobial target. Recent studies have outlined a mechanism of action for NrdR, in which ATP and dATP induce changes in the protein quaternary structure, regulating RNR repression. However, due to a lack of functional studies linking the known structures to their biological roles, the role of NrdR is not yet fully understood.

Here, we conducted a comprehensive study of NrdR in *Escherichia coli* and *Pseudomonas aeruginosa*. We delimited the NrdR regulon, combining transcriptomics and motif-based sequence analysis. We crystallized NrdR and obtained structural data on nucleotide binding and its relation to the stability of protein-protein interfaces involved in NrdR oligomerization. We examined the quaternary structures of NrdR using SEC-MALS and atomic force microscopy and correlated structure to function using point mutations, EMSAs, and *in vitro* transcription assays.

Overall, our results demonstrate the mechanism used by NrdR to modulate its quaternary structure and activity and provide structural data that pave the way for targeted antimicrobial therapies.

## 1. Introduction

Ribonucleotide reductases (RNRs) are the enzymes responsible for the reduction of ribonucleotides (NDPs or NTPs) to deoxyribonucleotides (dNDPs or dNTPs), thereby providing the building blocks for DNA synthesis and repair [1,2]. Three different RNR classes have been described: class I (subclassified into Ia, Ib, Ic, Id, and Ie), class II, and class III. RNR classes differ in their radical generation systems, electron donors, and quaternary structures [1–3]. However, all RNRs share a common free-radical-based mechanism and can reduce all four ribonucleotides [1,2,4,5]. Every class can be found in the domain Bacteria, and many species encode multiple RNRs [2,3,5]. Since all living cells require dNTPs, and a balanced supply of all four is critical to prevent mutations during DNA replication [6,7], RNR activity must be finely regulated at both the transcriptional and allosteric levels [2,3,8,9].

The allosteric regulation of RNRs ensures nucleotide homeostasis in two ways. First, it balances dNTPs through the “specificity” allosteric site present in all RNRs, which shifts enzymatic activity to the synthesis of purines or pyrimidines: if dATP is bound, purine synthesis is favoured, whereas dGTP or TTP binding promotes pyrimidine synthesis. [2,9–12]. Second, it balances ribonucleotide and deoxyribonucleotide levels. To achieve this, some RNRs include another allosteric site, the “activity” site. This second regulatory element is structured as a four-helix bundle covered by a three-stranded beta-sheet [13], forming a cone-shaped structure termed the “ATP-cone” [2,13]. The binding of ATP or dATP to the ATP-cone alters the quaternary structure of RNRs, respectively activating or inactivating the overall enzymatic activity [2,13] and consequently increasing or decreasing intracellular ATP or dATP levels.

The transcriptional regulation of RNRs governs the overall expression of RNR genes, coupling it to the cell cycle or increasing it in response to DNA damage [2,8]. In species encoding more than one RNR class, transcriptional regulation also controls differential RNR expression in response to environmental and metabolic conditions [2,8]. Notably, transcription factor NrdR, which is exclusive to bacteria, acts as a dedicated regulator of the entire RNR network [2,14,15].

NrdR was first identified in *Streptomyces coelicolor* [16]. It contains a Zn-finger DNA-binding domain and an ATP-cone domain resembling the “activity” allosteric site of RNRs [15–17]. NrdR was immediately hypothesized to regulate RNR transcription in response to nucleotide binding [15]. Bioinformatic studies identified NrdR orthologs across the domain Bacteria and found a conserved 16-bp sequence motif (NrdR-box) overlapping predicted RNR promoters exclusively in species encoding NrdR [14]. Two repeats of these NrdR-boxes, separated by an integer number of DNA helix turns, often overlap the core -10 and -35 promoter elements [14]. These findings suggested that NrdR functions as a transcriptional repressor of RNRs [14,15,17,18], forming oligomers that recognize both boxes simultaneously [17]. The distribution of *nrdR* genes and NrdR-boxes suggests that NrdR is present in most bacterial species, absent in *Eukarya* and *Archaea,* and regulates all RNR classes [14]. This makes NrdR a compelling potential target for antibacterial therapy.

NrdR has been studied in different bacterial species, including *S. coelicolor* [15,17,19], *Escherichia coli* [18] and *Pseudomonas aeruginosa* [20], consistently acting as a repressor of all RNR classes. Biochemical analyses have confirmed that NrdR contains zinc [15,17], binds nucleotides [17], and forms oligomers [15,17,18,21]. Further studies showed that, as suggested by its ATP-cone and Zn-finger domains, NrdR functions as a nucleotide-controlled transcription factor [22]. A negative cooperative mechanism has been proposed to explain how NrdR binds both ATP and dATP, despite the different orders of magnitude of their *in vivo* concentrations [23,24].

Recent structural studies confirmed the hypothesis that NrdR functions as a sensor of the dATP/ATP balance [19,25]. Crystal and cryo-EM structures of *S. coelicolor* and *E. coli* NrdR in complex with ATP, dATP, and DNA revealed that NrdR activity and oligomerization are indeed regulated by nucleotide concentrations. Grinberg *et al.* described two nucleotide-binding sites in the ATP-cone, similar to those observed in some RNRs [19,26]; the “inner” site binds ATP, while the “outer” site binds ATP or dATP. NrdR [19] bound to any nucleoside triphosphates forms tetramers via ATP-cone-to-ATP-cone and Zn-finger-to-Zn-finger interactions. At low dATP concentrations, two ATP molecules bind NrdR, resulting in a tetramer that further multimerizes into an inactive 12-mer (a trimer of tetramers). At higher dATP concentrations, binding of one ATP and one dATP (at the inner and outer sites, respectively) induces a conformational change, reorienting the Zn-finger domains, disrupting the 12-mer, and favouring the formation of an active octamer (dimer of tetramers). In the presence of target DNA, the active NrdR octamer disassembles into tetramers, with each tetramer binding two consecutive NrdR-boxes separated by 3 DNA helix turns [19]. *E. coli* NrdR displayed similar *in vitro* oligomeric rearrangements depending on the combinations of cofactors bound to its ATP-cone: ATP or ADP with dATP or dADP [25].

All previous studies on the oligomerization-based mechanism of NrdR have fallen short of describing individual NrdR arrangements in terms of absolute molecular weight or, crucially, demonstrating their biological role. The invaluable structural information provided by Grinberg *et al.* now requires functional data to fully establish the mechanism of action of NrdR. One major challenge in studying NrdR is the difficulty of expressing and purifying sufficient recombinant NrdR for molecular, biophysical, and structural studies. NrdR is unstable and tends to precipitate during and after purification [21,22,27]. Although the functional effect of NrdR has been studied using *in vitro* transcription (in *Chlamydia trachomatis*), protein instability made it extremely difficult to differentiate the effects of nucleotide cofactors [27].

In this study, we expand the existing structural information and link it to functional data to better understand the role, structure, and mechanism of NrdR across different bacterial species. We explored the NrdR regulon using transcriptomics and motif-based sequence analysis. To enable analytical experiments, we developed recombinant NrdR fusion proteins with enhanced stability, activity, and purity. We investigated the effects of different nucleotide cofactors on the quaternary structure of NrdR using atomic force microscopy (AFM), size exclusion chromatography (SEC), and SEC coupled with multi-angle light scattering (SEC-MALS). We crystallized *E. coli* NrdR purified in the presence of Zn^2+^, which provided a specific molecular arrangement with contact interfaces involving the Zn-finger and ATP-cone domains; we assessed the significance of these interfaces in oligomerization using targeted mutations. We studied the biological relevance of the observed NrdR quaternary structures using DNA binding assays and *in vitro* transcription experiments. Overall, our results provide extensive insights into NrdR oligomerization, offer functional evidence of the role of NrdR as a nucleotide sensor, extend our understanding of NrdR to new species, and provide detailed structural data that pave the way for the design of new antimicrobial therapies.

## 2. Results

### 2.1. Characterization of the NrdR regulon

Although NrdR is a well-known regulator of ribonucleotide reductase (RNR) transcription, multiple studies have suggested that its regulon may extend beyond that pathway. Non-RNR genes associated with NrdR-boxes, such as *dnaA* in *Shewanella* and *topA* in *Pseudomonas* [14], feature a single NrdR-box in their promoter regions, rather than the usual two boxes separated by an integer number of DNA helix turns, which are typically found in RNR promoters. Deactivating *nrdR* leads to numerous dysregulated genes beyond RNRs, as has been studied using DNA microarrays in *P. aeruginosa* PAO1 [20] and *E. coli* LF82 [28], as well by proteomics in *Bacillus subtillis* YB955 [29]. However, among the differentially-expressed genes, only RNRs were consistently associated with NrdR-boxes.

Unpublished lab DNA microarray data corroborate these findings for *E. coli* K-12; the results are included here for reference (Supplementary Table S1). We compared an *nrdR*ΔATPcone strain (encoding a non-functional NrdR without an ATP-cone) with its isogenic K-12 wild-type strain during both mid-exponential and stationary growth phases. 220 differentially expressed genes (DEGs) were identified, including all three RNR operons in *E. coli*.

To determine whether the observed transcriptomic effects were due to direct NrdR regulation, we conducted a global search for NrdR-boxes. Using the Gammaproteobacteria NrdR-box consensus [14] in FIMO [30,31], we searched upstream of every translation start codon in the *E. coli* K-12 genome (see Materials and Methods). This search revealed 48 putative NrdR-boxes (Supplementary Table S2), including the three previously identified box pairs [14,18] associated with operons *nrdAB-yfaE* (class Ia RNR), *nrdHIEF* (class Ib RNR), and *nrdDG* (class III RNR). In all these cases, NrdR-box pairs were separated by three whole turns of the DNA double helix (31-32 bp). Only one additional gene (transcriptional regulator *slyA*) exhibited two NrdR-boxes, although separated by 6.5 DNA helix turns.

Correlating the DEGs from the *nrdR*ΔATPcone strain to NrdR-box presence revealed that, among the DEGs with a log10 fold-change larger than 1.5 (|log(FC)| ≥ 1.5), only *nrdA* (class Ia RNR) and *nrdH* (class Ib RNR) displayed boxes in their promoter regions (Figure 1, left). Expanding the search to all DEGs, regardless of fold-change, revealed gene *nrdD* (class III RNR, active anaerobically), which is only weakly derepressed in a Δ*nrdR* background under aerobic conditions, as well as two more hits with NrdR-boxes (*dnaK* and *pdhR*), both featuring a single putative box (Supplementary Table S2). The remaining 211 DEGs were not associated with any NrdR-boxes, and all other 40 putative NrdR-boxes were not associated with DEGs.

**Figure 1.**
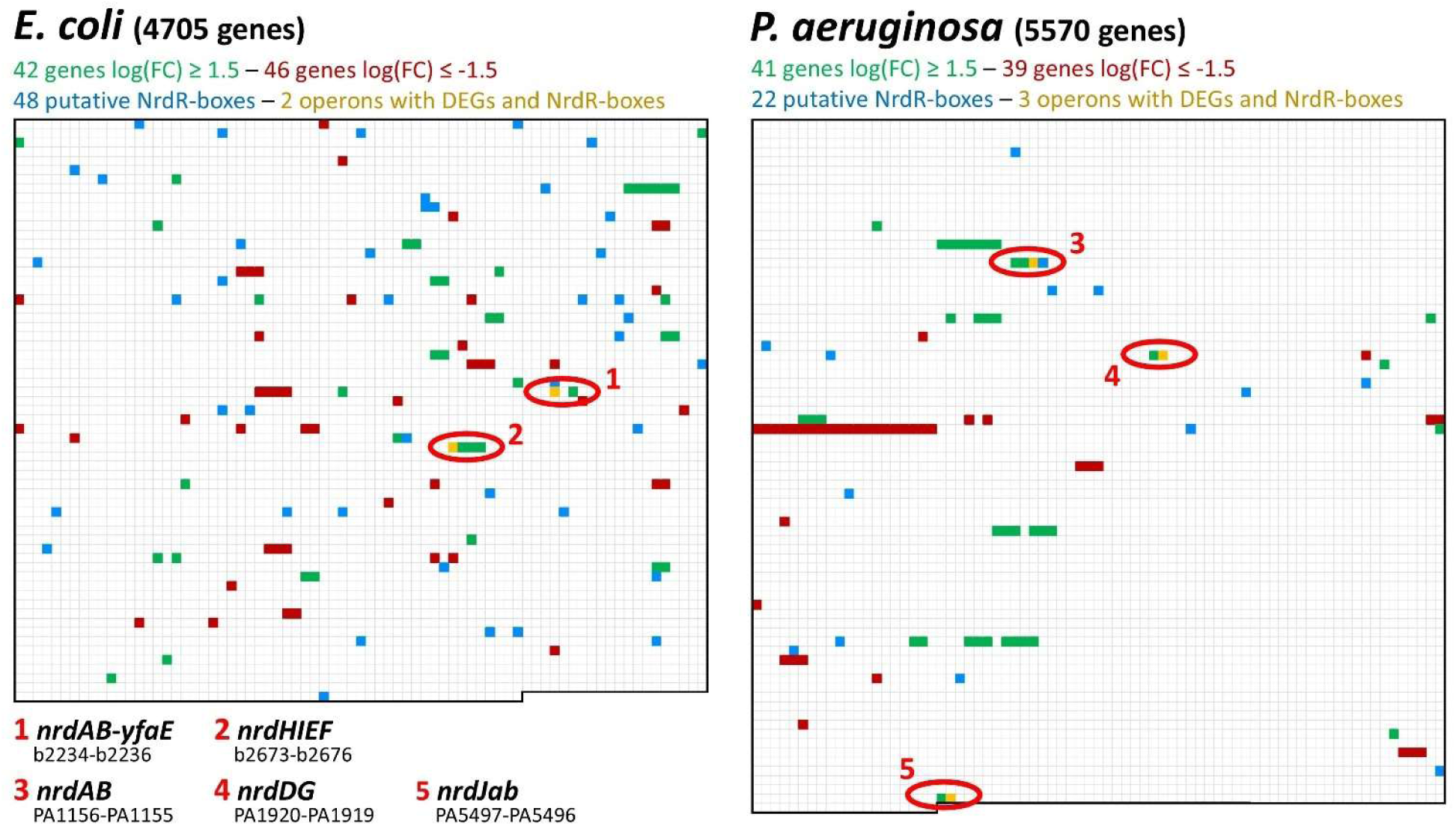
Correlation between DEGs in *nrdR* mutant strains and presence of NrdR-boxes. Data from *E. coli* K-12 *substr*. MG1655 (left) or *P. aeruginosa* PAO1 (right). Each square represents a gene in the genome, from top left (PA0001 *dnaA* in *P. aeruginosa*, b0001 *thrL* in *E. coli*) to bottom right (PA5570 *rpmH* in *P. aeruginosa*, b4705 *mntS* in *E. coli*). Green squares represent genes upregulated in an *nrdR* mutant strain compared to its isogenic WT strain (DNA microarray data for *E. coli*, Supplementary Table S1; RNA-seq data for *P. aeruginosa*, Supplementary Table S3), while red squares indicate downregulated genes. DEGs were only included if |log(FC)] ≥ 1.5. Blue squares represent genes for which at least one NrdR-box was found within 450 bp of its translation start codon; see Supplementary Table S2. Yellow squares represent the colocalization of DEGs and NrdR-boxes; operons including these genes are circled in red. Note that colocalization only occurs in RNR operons.

A similar search in the *P. aeruginosa* PAO1 genome revealed 22 putative NrdR-boxes (Supplementary Table S2). These included all eight previously identified [14,20], namely three associated with operon *nrdAB* (class Ia RNR[14]), two with *nrdJab* (class II RNR), two with *nrdDG* (class III RNR), and one with DNA topoisomerase I gene *topA*. Pairs of NrdR-boxes were all separated by three turns of the DNA double helix (31 bp). The third known box associated with *nrdAB* [14], situated over 200 bp upstream of the transcription start site, is of unknown function. Additionally, a previously undescribed fourth putative NrdR-box was identified further upstream of *nrdAB* (NrdR-box 4). No non-RNR gene presented more than one putative NrdR-box.

Attempting to correlate the NrdR-boxes found in *P. aeruginosa* to DEGs previously identified in a microarray study [20] revealed that, out of 143 DEGs identified under aerobic and anaerobic conditions, only *nrdA* and *nrdJa* presented NrdR-boxes in their promoter regions. However, as that study failed to detect differential expression in class III RNR operon *nrdDG* (even under anaerobic conditions) or *topA*, potentially due to limitations of the technique or biases of the platform, we conducted a thorough exploration of the effects of *nrdR* inactivation in *P. aeruginosa* PAO1 using RNA-Seq.

Compared to *P. aeruginosa* PAO1 wild-type, a Δ*nrdR* mutant strain (with its *nrdR* gene fully interrupted by a transposon insertion) showed 164 DEGs (104 upregulated and 62 downregulated) (Supplementary Table S3). All RNR operons were significantly upregulated: *nrdA* showed a fold-change of 2.96, *nrdJa* of 13.25, and *nrdD* of 13.65. Other upregulated genes included the entire PQS operon (*pqsABCDE*), responsible for one of the quorum-sensing systems in *P. aeruginosa*, several stress-related genes, such as the heat-shock genes *grpE* and *hslU*, and 14 genes related to the type III secretion system. A Gene Ontology Enrichment Analysis revealed hits for type III secretion system, interaction with host cellular response to heat, and protein folding pathways (Supplementary Figure S9; see Discussion). Downregulated genes included the arginine deiminase and pyoverdine biosynthetic operons. Interestingly, the topoisomerase gene *topA*, previously associated with a single NrdR-box, did not show differential expression.

When correlating these results with the presence of putative NrdR-boxes, out of 41 upregulated and 39 downregulated genes with a |log(FC)| ≥ 1.5, only the three RNR operons presented NrdR-boxes in their promoter regions (Figure 1, right). Expanding the search to all DEGs, regardless of log(FC), revealed no further correlations.

We cannot entirely rule out the existence of a mechanism of NrdR regulation based on a single NrdR-box or on binding sites different from the known NrdR-box consensus. Similarly, the indirect transcriptomic effects of ribonucleotide reductase dysregulation are worth exploring. However, only RNR operons simultaneously present pairs of NrdR-boxes and display differential expression upon *nrdR* inactivation, both in *E. coli* and *P. aeruginosa*. Therefore, we propose that the mechanism hypothesized for NrdR since its discovery, based on transcriptional repression via binding to pairs of NrdR-boxes overlapping with the core promoter, is limited to ribonucleotide reductases.

### 2.2. Design, expression, and purification of stable recombinant NrdR

To gain further insight into the mechanism NrdR uses to repress RNR expression, we aimed to obtain recombinant NrdR from both *E. coli* and *P. aeruginosa* for *in vitro* studies. However, NrdR is known to be highly unstable when overexpressed [15,18,22,27]. As part of this study, we developed different fusion proteins with varying compromises in yield, purity, and activity.

Our first attempt was purifying NrdR with a C-terminal 6His-tag (NrdR-H6). However, this protein was primarily found in inclusion bodies, and soluble fractions were highly unstable, precipitating throughout the purification process. This issue was particularly severe for *E. coli* NrdR-H6, which could never be recovered in an active form. For *P. aeruginosa*, despite significant losses due to precipitation, it was possible to obtain sufficient protein for analytical experiments (NrdR^PAO^-H6; details and SDS-PAGE in Supplementary Figure S1). PCA precipitation and ion-pair reverse-phase HPLC proved that NrdR^PAO^-H6 was nearly nucleotide-free, with only traces of ADP detected, corresponding to 1.8 nucleotides per 100 molecules (Supplementary Figure S2A, D). Furthermore, we demonstrated that NrdR^PAO^-H6 could bind ATP and dATP when incubated with a 20:1 nucleotide:protein ratio, although full occupancy was not achieved (Supplementary Figure S2A, E, F).

To enhance stability during purification and preserve functionality [32], we coupled *E. coli* and *P. aeruginosa* NrdR to an N-terminal 6His-Small Ubiquitin-like Modifier (SUMO). This tag helped solubility during overexpression and could later be removed using SUMO protease, leaving an untagged NrdR moiety with a two amino acid linker (NrdR1; see details and SDS-PAGE in Supplementary Figure S1). While this construct improved stability, NrdR1 from both microorganisms was produced at very low yields, limiting the number of experiments.

As the efficiency of SUMO digestion was identified as a major bottleneck in producing NrdR1, we developed a second generation of SUMOylated fusion proteins. This design included an additional TEV protease cleavage site (to use the SUMO tag only for its stabilization/solubilization properties) as well as an extended peptide linker (five amino acids) to improve digestion efficiency (NrdR2; details and SDS-PAGE in Supplementary Figure S1). NrdR2^PAO^ could not be obtained at a high yield and required AMP to be supplied throughout the purification process. In contrast, NrdR2^ECO^ was produced at high yields. PCA precipitation and HPLC analysis revealed that as-prepared NrdR ^ECO^ was recovered bound to ADP and AMP, with a total nucleotide occupancy over 80%, and efficiently bound ATP and dATP when incubated with a 20:1 nucleotide:protein ratio, reaching an occupancy of more than one nucleotide per protein (Supplementary Figure S2A, G, H, I).

Given its superior yield and stability, *E. coli* NrdR2^ECO^ was selected as the standard for subsequent experiments. For *P. aeruginosa*, NrdR^PAO^-H6 was regularly used, with NrdR1^PAO^ and NrdR2^PAO^ used for comparison when necessary.

### 2.3. *In vitro* NrdR dynamic oligomerization

#### 2.3.1. General overview

In multiple classes of ribonucleotide reductases, the ATP-cone domain regulates overall enzyme activity by inducing alterations in quaternary structure depending on the nucleotide bound [2,13]. A similar mechanism was hypothesized for NrdR since its discovery [15,21,22] as it contains a highly conserved ATP-cone domain. Recent studies explored the oligomerization of *E. coli* and *S. coelicolor* NrdR in detail [19,25]; the comparisons with the present study will be further assessed in the Discussion section.

Initial size-exclusion chromatography (SEC) experiments using NrdR^PAO^-H6, NrdR1^PAO^ (*P. aeruginosa*) or NrdR1^ECO^ (*E. coli*) demonstrated that NrdR undergoes dramatic, reproducible changes in quaternary structure depending on the bound nucleotide (see Supplementary Figure S3). When pre-incubated with a 20:1 nucleotide:protein ratio of AMP, all NrdR preparations primarily formed low-molecular-weight species consistent with a dimer or a trimer. In contrast, higher-order complexes were detected in the presence of dATP or ATP, with ATP inducing the highest apparent molecular weight. The limited yield and purity of the protein preparations complicated the interpretation of these results, although NrdR1^PAO^, being nearly nucleotide-free, provided the most reproducible data (see Supplementary Figure S3, B). In the presence of AMP, NrdR1^PAO^ (theoretical monomer Mw = 18.07 kDa) formed a broad peak centred at approx. 44 kDa (consistent with a mixture of dimers and trimers). Upon incubation with dATP, two peaks were observed centred at approx. 30 kDa (dimer) and 130 kDa (hexamer). ATP incubation yielded peaks at approx. 30 kDa (dimer) and 180 kDa (10-mer). Notably, peaks were broad, indicating they encompassed multiple conformations. This suggests that NrdR forms labile complexes with dynamic protein-protein associations.

#### 2.3.2. Diversity of NrdR quaternary structures

To confirm the proposed diversity of oligomeric states, we imaged the changes induced by nucleotide binding in the quaternary structure of NrdR2^ECO^ (*E. coli*) using atomic force microscopy (AFM). NrdR2^ECO^ was pre-incubated with nucleotides (20:1 nucleotide:protein ratio), deposited on mica, dried, and imaged in conventional dynamic mode (Figure 2A).

**Figure 2.**
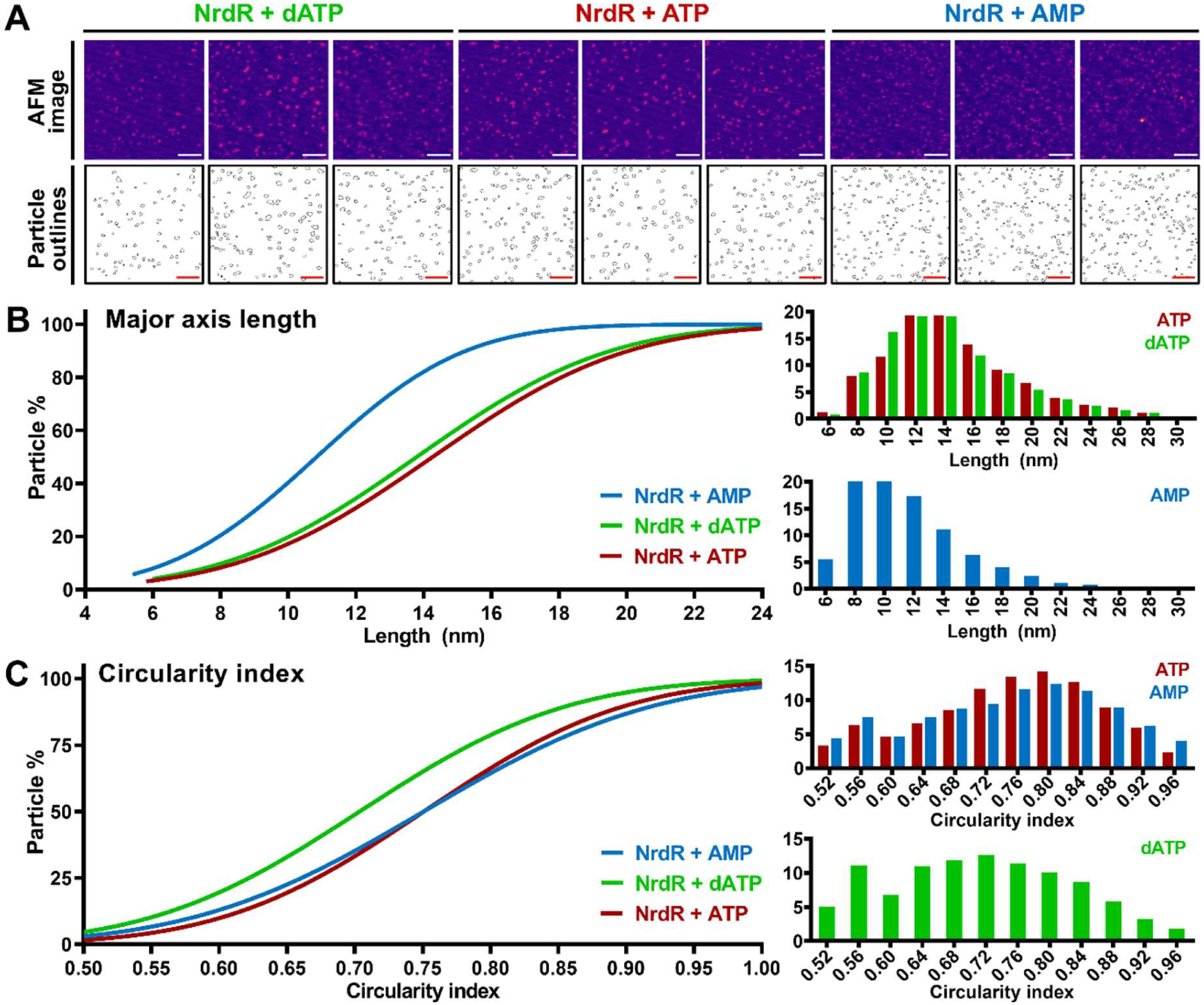
Influence of nucleotide cofactors on the NrdR quaternary structure determined by Atomic Force Microscopy. **A**: AFM images (top) and particle outlines (bottom) of representative NrdR-nucleotide complexes. Before imaging, NrdR ^ECO^ was incubated with cofactors at a 20:1 nucleotide:protein ratio. **B, C**: distribution of complexes based on size (B, major axis length of an ellipse containing the complex) and shape (C, circularity index of the complex, where 1 represents a perfect circle) depending on the nucleotide cofactor used. Data are represented as cumulative frequencies (left) and histograms (right). All distributions were determined as statistically distinct (nonlinear regression, assuming a cumulative Gaussian distribution; p-value ≤ 0 .05, extra sum-of-squares F test).

Compared to the dimensions of NrdR, the sizes of the structures observed were proven to be in the range of discrete NrdR oligomers rather than large aggregates (see Discussion). The size and shape distributions of the complexes (Figure 2B, C) revealed a broad range of structures. Protein oligomers were determined to be reproducibly smaller when coupled to AMP (Figure 2B), while nucleoside triphosphates induced the formation of larger structures. ATP produced a slightly larger complex than dATP, in what was proven to be a statistically significant difference. Analysis of the circularity index (which reflects how close the particle shape is to a circle) revealed that NrdR ^ECO^-dATP complexes were significantly more elongated than those bound to ribonucleotides (Figure 2C).

2.3.3. Absolute molecular weight of NrdR quaternary structures.

To resolve the absolute molecular weight of the complexes identified by SEC and AFM, we used SEC coupled with multiangle light scattering (SEC-MALS). We observed that incubating protein and cofactor resulted in different complex sizes depending on incubation time. To obtain reproducible results, we first exposed NrdR2^ECO^ to 0.025 mM nucleotide that was only present in the chromatography running buffer. Figure 3A shows SEC-MALS elution profiles and the weight-average molar mass (Mw) of the eluting samples (see Table 1, without pre-incubation).

**Figure 3.**
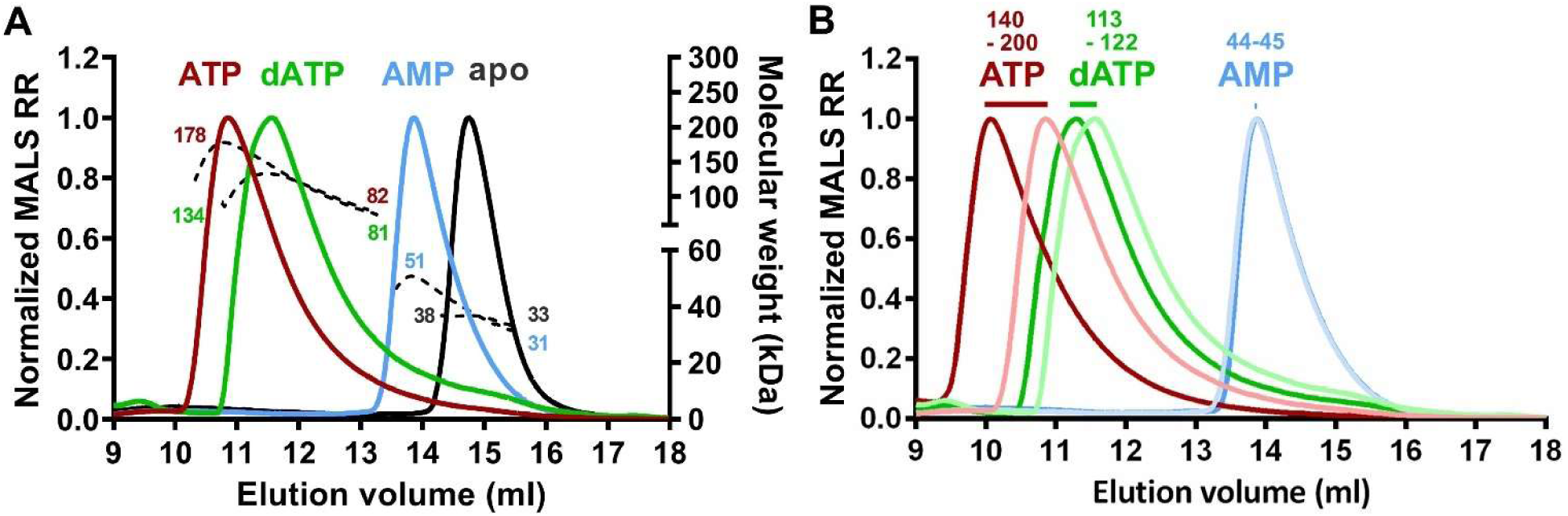
SEC-MALS study of NrdR-nucleotide complexes. **A**: SEC-MALS results of NrdR ^ECO^ exposed to 0.025 mM nucleotide in the running buffer. Peaks in solid lines represent MALS Rayleigh ratio normalized to a maximum of 1.0 for each sample (left Y-axis). Dashed lines represent weight-average molar mass (kDa; right Y-axis). Numbers indicate the minimum and maximum weight-average molar mass corresponding to each peak. Results are representative of two independent experiments. **B**: comparison of SEC-MALS results of NrdR_2_^ECO^ only exposed to nucleotides in the running buffer (light colours) or incubated with a 20:1 nucleotide:protein ratio for 3 hours before the chromatography, which also included nucleotide in the running buffer (dark colours). The difference in average Mw (kDa) is indicated above each peak.

**Table 1.** Molecular weight and quaternary structure of NrdR-nucleotide complexes determined by SEC-MALS. All data were obtained using NrdR ^ECO^. Top: results with nucleotide present only in the running buffer. Bottom: results with the protein pre-incubated with cofactors at a 20:1 nucleotide:protein ratio before the chromatography, which also included nucleotides in the running buffer. Mn, number-average molar mass; Mw, weight-average molar mass; Min/max, estimated minimum and maximum molar masses. The polydispersity index is defined as Mw/Mn. Protein conjugate analysis (ASTRA 7, Wyatt Technology) was applied to Nrd ^ECO^-nucleotide complexes to deconvolute the fractional mass contributed by protein and cofactor. All molar mass values are listed in kDa. Error is listed as ± standard deviation.

| Without pre-incubation |  |  |  |  |  |  |  |  |  |
| --- | --- | --- | --- | --- | --- | --- | --- | --- | --- |
| Nucl. | NrdR<sub>2</sub><sup>ECO</sup>- nucleotide complex |  |  |  |  |  | Protein conjugate analysis |  |  |
|  | Mn | Mw | Min | Max | Polydispersity | Interpretation | Mw (prot.) | Mw (co-factor) | Interpretation |
| + ATP | 131.52 ± 0.32 | 140.17 ± 0.30 | 81.96 | 178.32 | 1.07 ± 0.00 | R<sub>4</sub> ↔ R<sub>10</sub> | 135.59 ± 0.29 | 4.98 ± 0.30 | R<sub>8</sub> + ATP<sub>9-10</sub> |
| + dATP | 110.83 ± 0.21 | 113.36 ± 0.21 | 81.36 | 133.57 | 1.02 ± 0.00 | R<sub>4</sub> ↔ R<sub>8</sub> | 109.36 ± 0.21 | 4.36 ± 0.22 | R<sub>6</sub> + dATP<sub>8-9</sub> |
| + AMP | 43.81 ± 0.09 | 44.56 ± 0.08 | 30.93 | 50.84 | 1.02 ± 0.00 | R<sub>2</sub> ↔ R<sub>3</sub> |  |  |  |
| Apo | 35.95 ± 0.08 | 35.98 ± 0.08 | 32.78 | 38.23 | 1.00 ± 0.00 | R<sub>2</sub> |  |  |  |

**Table 1. Molecular weight and quaternary structure of NrdR-nucleotide complexes determined by SEC-MALS.**
| With pre-incubation |  |  |  |  |  |  |  |  |  |
| --- | --- | --- | --- | --- | --- | --- | --- | --- | --- |
| Nucl. | NrdR<sub>2</sub><sup>ECO</sup>- nucleotide complex |  |  |  |  |  | Protein conjugate analysis |  |  |
|  | Mn | Mw | Min | Max | Polydispersity | Interpretation | Mw (protein) | Mw (co-factor) | Interpretation |
| + ATP | 179.75 ± 0.76 | 200.31 ± 0.53 | 75.11 | 266.47 | 1.11 ± 0.01 | R<sub>4</sub> ↔ R<sub>16</sub> | 193.80 ± 0.51 | 7.09 ± 0.53 | R<sub>10</sub>+ATP<sub>13-15</sub> |
| + dATP | 118.52 ± 0.22 | 121.83 ± 0.21 | 80.06 | 139.16 | 1.02 ± 0.00 | R<sub>4</sub> ↔ R<sub>8</sub> | 117.49 ± 0.22 | 4.73 ± 0.22 | R<sub>6</sub> + dATP<sub>9-10</sub> |
| + AMP | 44.02 ± 0.10 | 44.75 ± 0.09 | 31.07 | 54.53 | 1.02 ± 0.00 | R<sub>2</sub> ↔ R<sub>3</sub> |  |  |  |

As-prepared NrdR ran as a dimer, with an average Mw of 35.98 kDa (theoretical monomer Mw = 17.66 kDa). In the presence of AMP, NrdR2^ECO^ eluted with an average Mw of 44.56 kDa, ranging from 50.84 (trimer) to 30.93 (dimer). ATP and dATP induced higher-order oligomerization of NrdR2^ECO^: NrdR-ATP presented an average Mw of 140.17 kDa, ranging from 81.96 (tetramer) to 178.32 (10-mer), while NrdR-dATP averaged 113.36 kDa, ranging from 81.36 (tetramer) to 133.57 (octamer). The broad molecular weight ranges and high polydispersity again suggest dynamic complex interactions. Interestingly, deconvoluting the fractional mass contributed by protein and ligand using protein-conjugate analysis (Table 1) identified that cofactors were bound in stoichiometries higher than 1:1 at the corresponding peak maxima (8 NrdR + [9–10] ATP; 6 NrdR + [8–9] dATP). SEC-MALS experiments with NrdR2^PAO^ in complex with nucleotides were challenging to perform and only interpretable on one occasion, as the protein was highly unstable and had to be co-purified with AMP. NrdR2^PAO^-AMP showed a narrower peak with a molecular weight consistent with a dimer, whereas in the presence of dATP and ATP, a mixture of oligomers ranging from dimers to higher molecular weight species was identified (Supplementary Figure S4).

Finally, we evaluated the molecular weight shift when pre-incubating the protein for 3 hours with cofactors at a 20:1 nucleotide:protein ratio (keeping the cofactor in the SEC running buffer as well). This pre-incubation accentuated the differences between the effects of the cofactors (Figure 3B; Table 1, with pre-incubation). NrdR-AMP did not vary, maintaining a weight-average molar mass of 44.75 kDa. NrdR-dATP shifted slightly to 121.83 kDa, ranging from tetramers to octamers. NrdR-ATP, however, exhibited a major shift, reaching an average Mw of 200.31 kDa and ranging from 75.11 kDa (tetramer) to 266.47 kDa (15-mer to 16-mer).

### 2.4. NrdR2^ECO^ crystal structure

#### 2.4.1. Crystal structure and contact interfaces

To learn more about the mechanism of action of NrdR and how it relates to its quaternary structure, we attempted the crystallization of NrdR2^ECO^. Efforts to crystalize this protein saturated with ATP or dATP were unsuccessful (see Discussion). Instead, crystallization without adding nucleotides yielded a full-length structure of NrdR2^ECO^ at 2.6 to 2.8 Å resolution (PDB ID: 9GEX, Figure 4; see monomer details in Supplementary Figure 5A and data collection parameters and statistics in Supplementary Table S4).

**Figure 4.**
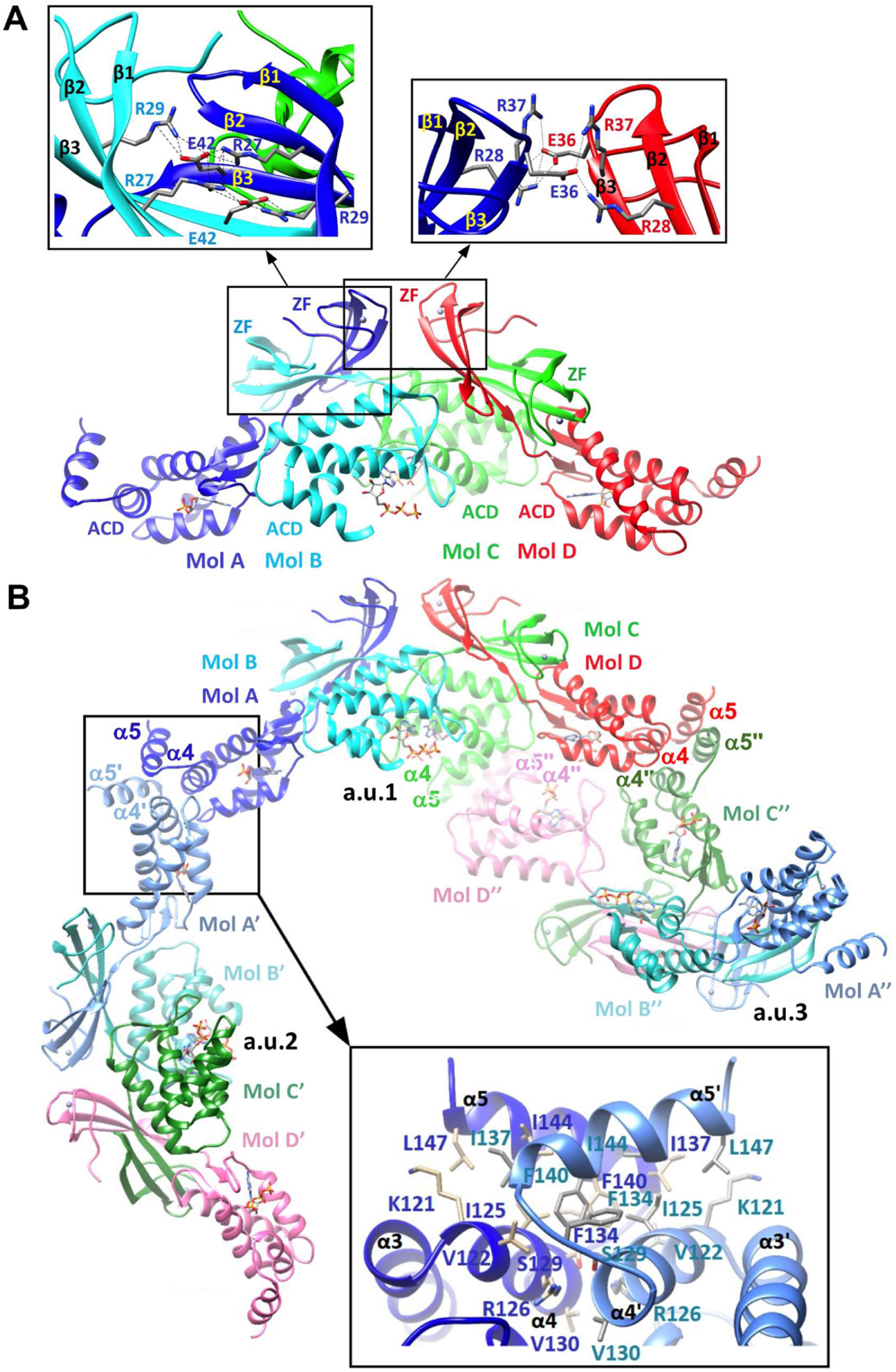
Structure of NrdR_2_^ECO^ crystallized without adding nucleotides. **A**: overall arrangement of the asymmetric unit. A crystallographic pair of dimers consists of dimer AB (dark blue, turquoise) and dimer CD (green, red). Each molecule has an N-terminal zinc-finger domain (ZF) and an ATP-cone domain (ACD) (See Supplementary Figure S5A). Top left inset: close-up view of the head-to-head intra-dimer interaction between molA and molB zinc-fingers involving Arg27, Arg29, and Glu42 from both molecules. Top right inset: close-up view of the inter-dimer interactions between zinc-fingers from molA and molD, involving Arg28, Glu36, and Arg37. **B**: symmetry contacts with AB/CD groups from neighbouring asymmetric units. Contacts are shown between a central a.u.1 complex, (AB/CD) a.u.2 (A’B’/C’D’) and a.u.3 (A’’B’’/C’’D’’). Bottom right inset: close-up view of the interaction between helices α4 and α5 from symmetry partners A and A’. Protein molecules are represented as ribbons: α-helices are depicted as helices, β-strands as arrows, highlighted amino acids and nucleotides as sticks.

The structure shows four protein molecules (molA, molB, molC, and molD) in the asymmetric unit (a.u.) (Figure 4A). The polypeptide chain folds in an elongated N-terminal zinc-finger domain (aa 1-44) composed of three β-strands (β1-β3); four cysteines (Cys3, Cys6, Cys31, Cys34) coordinate a Zn^2+^ ion at the tip of the domain (Supplementary Figure S5B). The zinc-finger is connected via a short loop to an ATP-cone domain (aa 48-131) consisting of a small three-stranded β-sheet (β4-β6) that acts as a lid covering the aperture of a cone formed by four parallel α-helices (α1-α4). Following the ATP-cone, helix α5 exhibits variable orientations and electron density quality across the four molecules: in molA, it is clearly defined, as it is stabilized by hydrophilic and hydrophobic interactions with symmetry mates, whereas in molB, it is exposed to the solvent and remains untraceable, suggesting high flexibility. ATP, ADP, and AMP molecules bound to NrdR were identified (see below).

The four protein molecules in the a.u. are organized as two pairs: AB (1357.1 Å^2^ of solvent accessible area buried by contact, complex formation ΔG = -7.5 Kcal/mol) and CD (1518.2 Å^2^, ΔG = -9.7 Kcal/mol) (Figure 4A). The extensive contact interfaces within the AB and CD pairs (which exceed the critical threshold of 1000 Å^2^) and the negative complex formation ΔG determined by PDBePISA [33] suggest AB and CD represent dimers that would be stable in solution. The AB/CD difference in ΔG can be attributed to small structural rearrangements.

AB and CD dimers present equivalent internal contacts between monomers; AB will be the reference. The AB zinc-fingers intertwine and contact head-to-head, with Arg27 and Arg29 from molA contacting Glu42 from molB and vice versa (intra-dimer contacts; Figure 4A, top left inset). The ATP-cones contact head-to-tail: the loop between helices α1 and α2 at molB contacts the β-sheet lid of molA’s cone. These contacts involve Van der Waals forces, two hydrogen bonds, and a water molecule. The shift from zinc-fingers head-to-head to ATP-cones head-to-tail is mediated by a conformational change at Leu46 in the inter-domain loop (the Ψ angle changes from 46° in molA to 165° in molB), suggesting a hinge function.

In the a.u., pair CD is rotated 180° relative to pair AB, both dimers packing back-to-back. The zinc-fingers of the two dimer pairs contact via an A:D interface, where Arg28 and Arg37 from one molecule form a salt bridge with Glu36 from the other (inter-dimer contacts; Figure 4A, top right inset). This interaction results in a BA:DC zinc-finger arrangement that creates a small, hydrophilic, and unfavourable transient interface (176.3 Å^2^, ΔG = +3.4 Kcal/mol). The Leu46 hinge change leads to an AB:CD orientation for the ATP-cones, which form inter-dimer contacts involving molB and molD, resulting in a small but slightly favourable interface (369.6 Å^2^, ΔG = -1.9 Kcal/mol). Additionally, the molB ATP-cone forms a second contact with the molD zinc-finger (174.7 Å^2^, ΔG = -2.0 Kcal/mol) and molC with molA (205.8 Å^2^, ΔG = +0.1 Kcal/mol).

Compared to the strong and favourable intra-dimer contacts, the weak inter-dimer interactions suggest that the a.u. represents two stable NrdR dimers forming labile interactions with one another rather than a biologically relevant tetramer. These AB and CD dimers expand throughout the crystal, stabilized by symmetry contacts between helices α4 and α5 (Figure 4B). Specifically, molA α4 and α5 interact with symmetric molA’ α5’ and α4’, respectively, whereas molC α4 and α5 interact with symmetric molD **’** α5’’ and α4’’, and vice versa. However, note that the variable electron density quality of helices α5 in molecules A, C, and D makes comparisons between these interactions inconclusive.

#### 2.4.2. Nucleotide co-factor binding

Within the ATP-cones, extra density in molB and molD allowed us to trace ATP molecules (Figure 5B, D), while in molC, we identified an ADP (Figure 5C). These nucleotides were oriented similarly to the activity site of RNRs (e.g., see *E. coli* class Ia RNR R1 subunit bound to AMP-PNP; PDB ID 3R1R), which corresponds to the “inner” nucleotide binding site in *Streptomyces coelicolor* NrdR as determined by Grinberg *et al.* [19]. In molA, density observed after averaging diffraction data from three datasets allowed us to trace an AMP molecule in this site (Figure 5A, Supplementary Figure 5C; see Materials and Methods).

**Figure 5.**
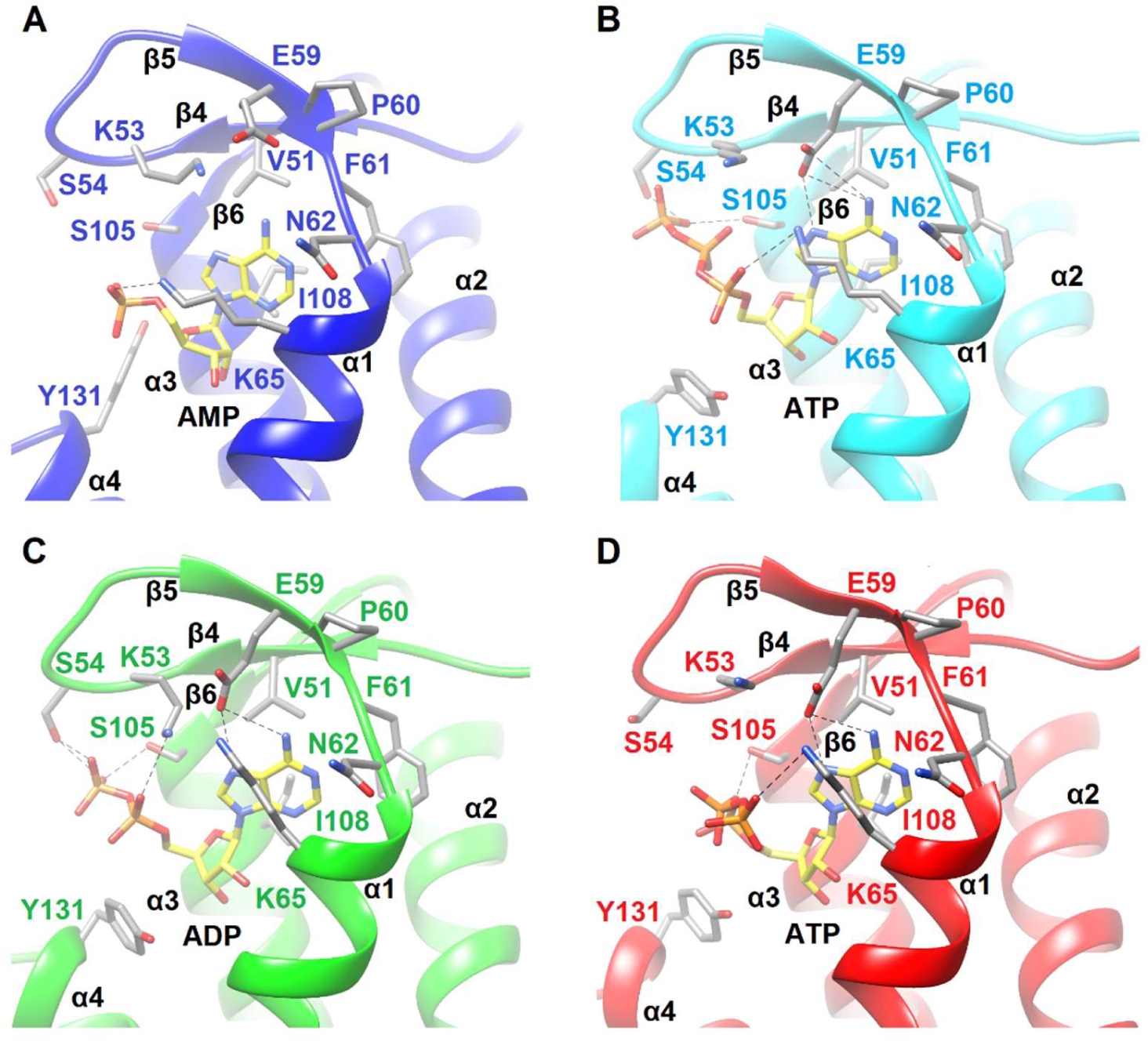
Nucleotide binding to NrdR ^ECO^. AMP was found bound to molA (**A**), ATP to molB and D, in two different orientations (**B**, **D**), and ADP to molC (**C**). MolA’s cofactor was resolved only after averaging three individual datasets. Residues relevant for nucleotide binding are highlighted. ADP or ATP require additional contacts to stabilize the extra phosphate groups. Protein molecules are represented as ribbons: α-helices are depicted as helices, β-strands as arrows, and highlighted amino acids and nucleotides as sticks.

Nucleotides within the cone cavity are stabilized by the β-sheet lid and helices α1, α3, and α4. The cone features a polar entry, where Lys53 and Glu59 (at the lid) and Lys65 (helix α1) cover the adenine base. The adenine rings, in turn, lie over a hydrophobic pocket formed by Val51, Phe61, Leu66, and Ile108.

Importantly, the main chain carbonyl of Pro60, located at the end of the lid and preceding the cone, interacts with the adenine N6 atom, suggesting a role in the specific recognition of adenine nucleotides. Pro60 and the next loop (Phe61-Asn62) are at the B:C ATP-cone interface, establishing a small Van der Waals contact that positions the respective molB and molC nucleotide pockets in an antiparallel face-to-face arrangement at the core of the AB:CD complex. At the opposite end of the rim, the cofactor phosphate groups, which are more exposed to the solvent, present different orientations, consistent with a more labile conformation of this region. For example, in molB, the ATP γ-phosphate group contacts Ser54 and Ser105 -OH groups (cone rim, near helix α3), whereas, in molD, the ATP phosphate groups fold over, positioning the γ-phosphate in close proximity to Lys65 -NH2 group (helix α1), which entails the remodelling of all remaining contacts (Figure 5). Within the Van der Waals distance of the phosphate groups, Tyr131 (helix α4), described as essential for ATP/dATP discrimination at the “outer” nucleotide binding site in *S. coelicolor* NrdR [19], can be seen in molecules B, C, and D orienting the side chain towards α1 and the empty outer site, contacting Arg72 (helix α1) in molB. In molA, Tyr131 is instead oriented towards helix α3, with its -OH group contacting the Ser105 main chain carbonyl. These alternate orientations suggest a gatekeeping role for Tyr131 in the absence of a cofactor at the outer site.

### 2.5. Mutational study of NrdR contact interfaces and oligomerization

To determine whether the interactions observed in the NrdR2^ECO^ structure were biologically relevant, we introduced mutations designed to disrupt the observed interacting surfaces while minimizing additional impact (see the context of mutations in Figures 4A and 6A). Mutation E42A was designed to abolish the intra-dimer salt bridges between zinc-fingers (Glu42 with Arg27 and Arg29); unlike the other amino acids involved, Glu42 is not described as essential for DNA binding [19]. Mutation E36A was designed to disrupt inter-dimer zinc-finger contacts (Glu36 with Arg28 and Arg37); Glu36 is also the only amino acid involved that is not essential for DNA binding. Given the importance of the observed a.u. contacts between symmetry mates (helix α4 with α5), we tested NrdR functionality after a complete deletion of helix α5 and the preceding loop (mutation Δ132-149). Additionally, we mutated Tyr131 (mutation Y131A) due to its recognized functional importance for ATP/dATP discrimination at the outer nucleotide-binding site [19] to assess its impact on oligomerization.

**Figure 6.**
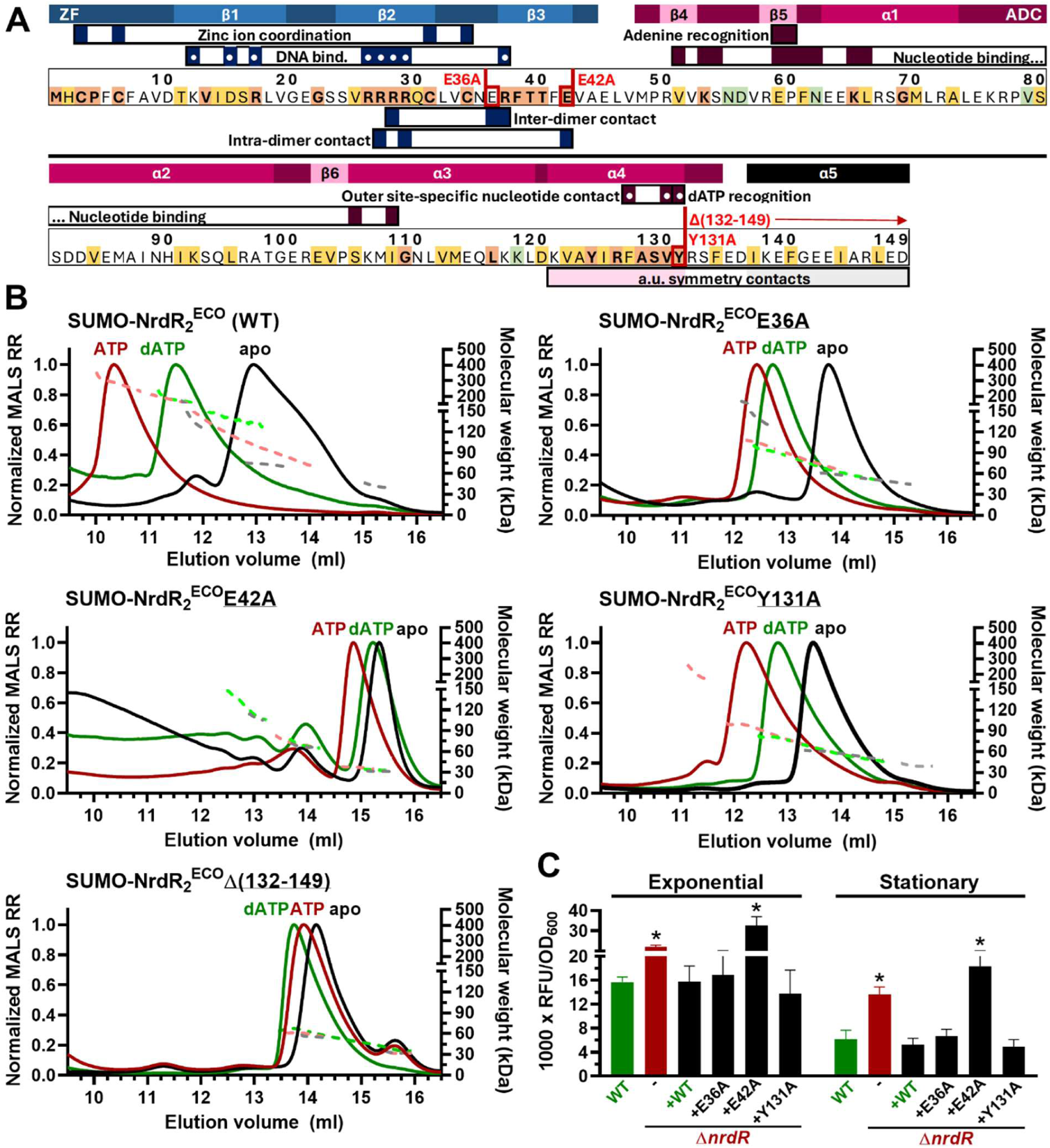
Effects of mutations in crystalline contact interfaces on NrdR_2_^ECO^ mutant derivatives. **A**: Schematic showing mutations E36A, E42A, Y131A, and Δ(132-149) in the context of relevant positional information about NrdR. The amino acid sequence of *E. coli* NrdR is coloured to indicate the degree of conservation: orange (most conserved) → yellow → green → white (least conserved) (see Supplementary Figure S6). The zinc-finger domain (ZF, blue) and ATP-cone domain (ACD, magenta) are indicated along with secondary structure elements. Relevant residues for different functional aspects of NrdR are labelled. A white dot marks information obtained on *S. coelicolor* by Grinberg *et al*. [19]. **B**: SEC-MALS results of NrdR_2_^ECO^ and its mutant derivatives, either as-prepared (apo) or pre-incubated with ATP or dATP. The left Y-axis (solid lines) shows MALS Rayleigh ratio normalized to a maximum of 1.0. The right Y-axis shows weight-average molar mass (kDa); slope lines indicate mixtures of oligomers. **C**: GFP-based reporter gene assays showing the *in vivo* effects of mutations in crystalline contact interfaces. Expression of class Ia RNR promoter P*nrdA* was measured in *E. coli* K-12 (WT), its isogenic *nrdR* deletion mutant (Δ*nrdR*, *-*) and Δ*nrdR* complemented with pLG338 carrying wild-type *nrdR* or its mutant derivatives. Samples were collected at OD_600_ = 0.50 (exponential) and OD_600_ = 2.50 (stationary). The Y-axis represents relative fluorescence units normalized by dividing by OD_600_. Results are the average of three biological replicates. *, p-value ≤ 0.05 (two-tailed t-test), compared to WT.

First, mutant NrdR2^ECO^ derivatives were produced and purified using the same method as the WT protein to conduct SEC-MALS experiments. However, despite the remarkable stability of the NrdR2^ECO^ construct, all mutants except Y131A displayed severe instability after SUMO excision, leading to progressive protein loss during purification. To ensure consistency, we compared the oligomerization of undigested SUMO-NrdR2^ECO^ and its mutant derivatives.

SEC-MALS analysis of wild-type SUMO-NrdR2^ECO^ without additional nucleotides revealed three peaks (Figure 6B, top left; Supplementary Table S5). The largest peak, representing more than 80% of the protein mass, corresponds to dimers, as previously observed for the digested protein (average Mw 72.99 kDa, for a theoretical monomer Mw of 31.23 kDa); smaller peaks represent monomers (Mw from 39.84 kDa) and tetramers and higher oligomers (average Mw 136.31 kDa, range 121.48-172.34). Pre-incubation with dATP increased the oligomeric state, resulting in a larger peak of tetramers (Mw from 127.20 kDa) and one encompassing states from tetramer to octamer (average Mw 176.80 kDa, range 156.90-242.66). In the presence of ATP, SUMO-NrdR formed larger oligomers, with molecular weights ranging from dimer to 12-mer (70.09-352.86 kDa, resolved during MALS data processing as two populations with average Mw 145.84 and 275.41 kDa, respectively).

All mutations targeting crystal contact interfaces caused drastic changes in multimerization. Mutants E36A (inter-dimer contacts disruption; monomer Mw 31.17 kDa) and Y131A (outer site recognition and ATP-cone gatekeeper disruption; monomer Mw 31.14 kDa) displayed similar behaviour. In the absence of nucleotides, both mutants predominantly formed dimers: for E36A (Figure 6B, top right; Supplementary Table S5), we identified two molecular populations leading up to a maximum Mw of 60.94 kDa (dimer) and a small fraction of tetramers and higher oligomers (average Mw 141.21 kDa, range 125.45-172.34). Y131A showed a small monomer peak (average Mw 39.97 kDa) and two close populations leading up to a maximum Mw of 62.03 kDa (dimer). However, in the presence of dATP or ATP, both E36A and Y131A failed to form tetramers or higher-order oligomers. Mutation E36A, in particular, resulted in dimers (average Mw 68.38 kDa) and a few trimers (average Mw 92.02 kDa) after dATP pre-incubation, with minimal change in the presence of ATP: dimers (average Mw 72.61 kDa, range 64.38-90.60) and trimers (average Mw 102.84 kDa, range 95.36-108.33), indicating the relevance of Glu36 for multimerization. Mutation Y131A (Figure 6B, mid right; Supplementary Table S5) similarly produced dimers (average Mw 59.90 kDa) and limited higher oligomerization (average Mw 79.62 kDa, range 75.36-84.13) after dATP pre-incubation; ATP elicited some tetramers and octamers, but these represented only 3% of the protein mass (average Mw 189.95 kDa, range 161.47-263.17), with the remainder consisting of dimers and trimers (average Mw 69.79 and 96.78 kDa, respectively), underpinning the importance of this residue.

The changes in oligomerization caused by these mutations may be due to the disruption of the contact interfaces observed in the crystal structure but may also be explained by altered cofactor binding. Tyr131 is a highly conserved residue across the Bacteria domain (Supplementary Figure S6). In the NrdR2^ECO^ structure, two rotamers are observed: one oriented towards α1 and one towards α3. In the published cryo-EM structure of *S. coelicolor* NrdR, the equivalent Tyr128 orients towards α1 when ATP is bound to the outer site and towards α3 when dATP is bound; hence, mutation Y131A undoubtedly disrupts nucleotide binding. Additionally, Tyr131, at the tip of helix α4, forms Van der Waals contacts with the phosphate groups from the inner site nucleotide; mutation Y131A loses these contacts, presumably destabilizing the α4-α5 interface, thereby cancelling potential protein-protein interactions. Mutation E36A, which caused a similar oligomerization profile, disrupts inter-dimer salt bridges between molA and molD zinc-fingers, resulting in strong electrostatic repulsion between Arg28 and Arg37 due to the lack of the Glu36 compensatory negative charge. However, Glu36 is not a universally conserved residue and is instead restricted to Proteobacteria (Supplementary Figure S6). The potential presence of these protein-protein interfaces is further assessed in the Discussion section.

The other mutations severely impacted oligomerization. The Δ(132-149) mutant (monomer Mw = 29.06 kDa) showed only monomers and dimers (Figure 6, bottom left; Supplementary Table S5) regardless of the cofactor: as-prepared (Mw averages 32.52 and 55.08 kDa; dimer and monomer), with dATP (34.87 and 62.19 kDa; dimer and monomer) or with ATP (31.16 and 59.24 kDa; dimer and monomer). Mutation E42A (intra-dimer zinc-finger contacts disrupted; monomer Mw = 31.17 kDa) had an even more drastic effect, causing most NrdR to remain as a monomer, with only small dimer peaks and no difference between cofactors (Figure 6, mid left; Supplementary Table S5): as-prepared (85% of protein mass concentrated in a peak with average Mw 31.39 kDa), with dATP (63% in a peak with average Mw 33.21 kDa), or with ATP (83% in a peak with average Mw 36.88 kDa). In contrast to Glu36, Glu42 is highly conserved (Supplementary Figure S6), and the major impact of the E42A mutation suggests a pivotal role for this intra-dimer contact surface (see Discussion).

Given the effect that disrupting contact surfaces had on oligomerization, we tested them *in vivo* by evaluating the ability of the mutant NrdR2^ECO^ derivatives to complement an *nrdR* deficient background. We measured the expression of class Ia RNR promoter (P*nrdA*) driving GFP expression in an *E. coli* K12 strain and its isogenic Δ*nrdR* deletion mutant, as well as the Δ*nrdR* strain transformed with low-copy-number plasmid pLG338 carrying either wild-type *nrdR* or its mutant derivatives (E36A, E42A, and Y131A) (Figure 6C). As expected, the Δ*nrdR* mutation derepressed P*nrdA*, increasing its expression by around 40% during exponential growth and more than doubling it during the stationary phase. Wild-type *nrdR* restored *nrdA* expression levels to normal. Similarly, the E36A and Y131A mutants fully rescued NrdR function. However, the E42A mutant failed to restore P*nrdA* expression levels and even significantly increased class Ia RNR transcription. These results underscore the importance of the Glu42 intra-dimer interaction surface, as its disruption profoundly alters NrdR functionality.

### 2.6. Effects of nucleotide cofactors and quaternary structure on *in vitro* NrdR activity

#### 2.6.1. Effects on DNA binding capacity

Once the quaternary structures of NrdR bound to nucleotides were characterized structurally, in solution, and with point mutations, the next step was to explore the functional significance of these oligomerization differences. The DNA-binding capabilities of NrdR have been studied by electrophoretic mobility shift assays (EMSA) in *E. coli* [18,22], *Chlamydia trachomatis* [27], and *Salmonella typhimurium* [34], although challenges related to the activity, purity, and specificity of recombinant NrdR made interpreting the results difficult. Similarly, early experiments in this study using NrdR^PAO^-H6 and NrdR1^ECO^ presented significant issues, including protein aggregation and nonspecific binding to negative controls (DNA lacking NrdR-boxes). Improved specificity and reproducibility for NrdR binding to its target DNA were achieved using NrdR2^ECO^ *(E. coli*), adding fresh DTT, Mg^2+^, and Zn^2+^ to the binding buffer, using 5% triethylene glycol in the EMSA gels to stabilize the complexes, and with high amounts of competition DNA to reduce nonspecific binding (2 µg of salmon sperm DNA, equivalent to an approximate mass ratio of 65:1 competition:probe DNA).

Under these optimized conditions, the EMSA results with an NrdR-sensitive probe (the *E. coli* class Ia RNR promoter, referred to as P*nrdA*) clearly showed the effects of the different nucleotide cofactors (Figure 7A, left). Without nucleotide addition or with AMP, NrdR caused a smeared band shift consistent with a small, unstable complex. NrdR-ATP failed to cause a significant shift, indicating an inability to form a stable complex. NrdR-dATP produced a distinct shift consistent with the binding of a large protein oligomer. NrdR-dATP also showed increased binding to non-specific DNA (Figure 7A, right), though this effect was markedly smaller than with the specific probe. ADP rendered NrdR inactive, while dAMP induced the same unstable band shift as AMP (Supplementary Figure S7, top). NrdR2^PAO^ (*P. aeruginosa*) results mirrored those of its *E. coli* counterpart, with stable shift bands appearing only in the presence of dATP, which also caused a modest increase in nonspecific binding (Supplementary Figure S7, bottom). A substantial portion of the *P. aeruginosa* preparation, however, appeared to be inactive.

**Figure 7.**
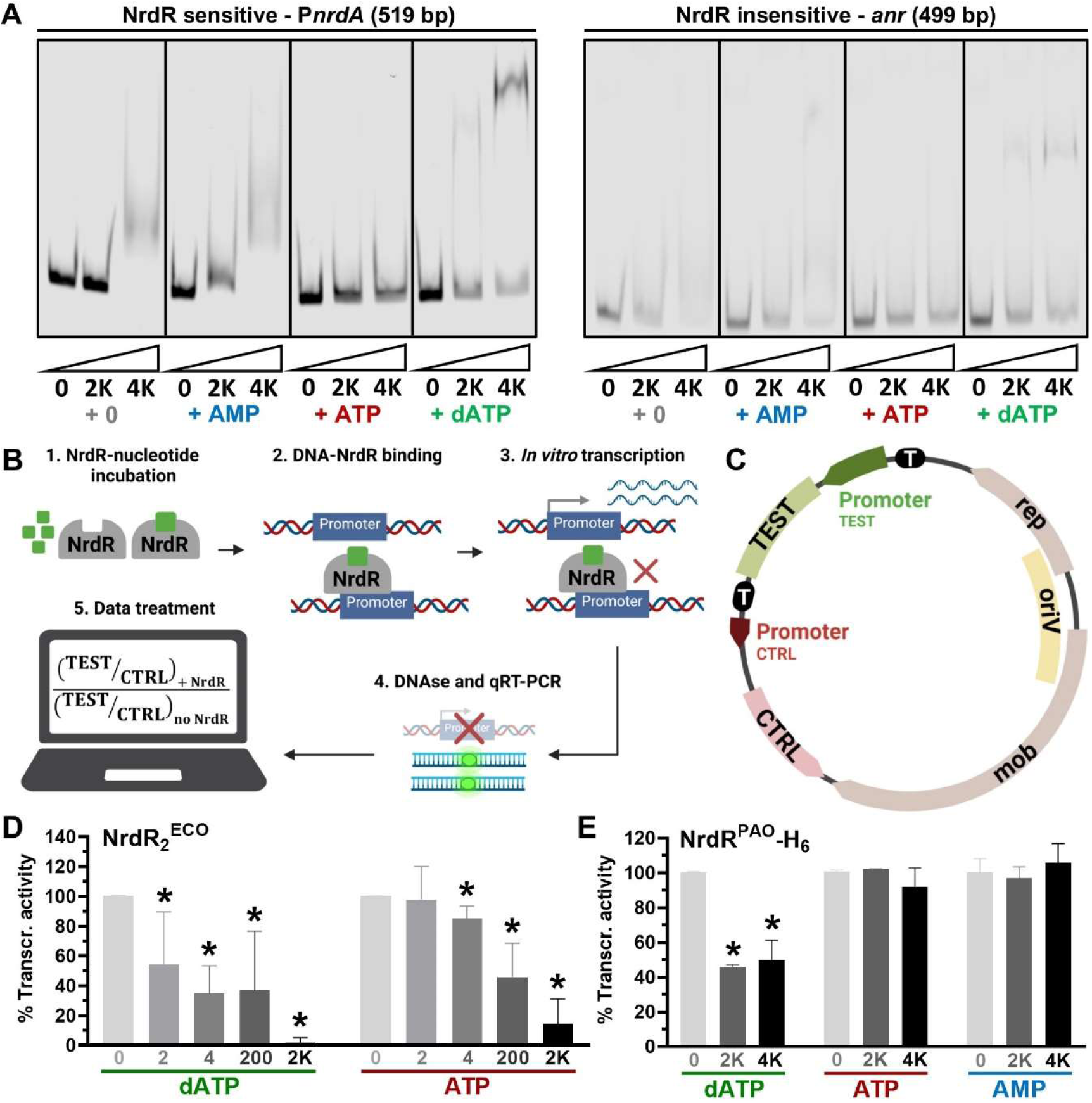
Nucleotide-dependent NrdR activity. **A**: EMSA results for NrdR_2_^ECO^. Two DNA probes were used: P*nrdA*, the NrdR-sensitive promoter of *E. coli* class Ia RNR (left), and a negative control probe *anr* (right). The molar ratio of NrdR to labelled DNA is indicated below the images (0:1, 1000:1, 4000:1). Nucleotides were pre-incubated with NrdR at a 20:1 nucleotide:protein ratio. Representative data from three independent experiments are shown. **B**: schematic of the Reverse *In Vitro* Transcription Assay (ReViTA) used to evaluate transcription factors *in vitro*. Created in BioRender (Pedraz, L., 2025, https://BioRender.com/g34i593). **C**: simplified schematic of the pReViTA plasmids, which include TEST promoters to study the *in vitro* transcription of different operons and a CTRL promoter for normalization. Black segments labelled T represent transcription terminators (see Rubio-Canalejas *et al.* [35]). **D, E**: repression of *E. coli* class III RNR promoter (P*nrdD*) transcription by NrdR coupled to different nucleotide cofactors. NrdR_2_^ECO^ (D) or NrdR^PAO^-H_6_ (E) was incubated with a 20:1 nucleotide:protein ratio and added to 100 fmol of DNA template; the numbers under each bar represent the protein:DNA ratio. The transcription of the TEST sequence was normalized using the unregulated CTRL sequence (see Materials and Methods) and expressed as a percentage of the unrepressed (no NrdR) transcription (% transcriptional activity). Results are the average of three biological replicates. *: p-value ≤ 0.05 (two-tailed t-test), compared to the no NrdR reaction.

#### 2.6.2. Effects on *in vitro* transcription repression activity

After the differences in NrdR DNA-binding capacity had been attested, we assessed their functional implications by measuring whether NrdR bound to different cofactors could repress RNR transcription. To address this, we employed the reverse *in vitro* transcription assay (ReViTA). Briefly, in this technique, two genes are transcribed in a single reaction: one under the control of an NrdR-sensitive promoter (TEST) and another under a constitutive promoter (CTRL). Both genes are encoded on a circular, synthetic plasmid template (pReViTA, Figure 7C) separated by transcriptional terminators. Relative transcription of TEST and CTRL is quantified using qRT-PCR, enabling a precise determination of transcription factor activity (see our previous publication of this methodology [35]). To test the nucleotide-dependent activity of NrdR, the protein was pre-incubated with cofactors (20:1 nucleotide:protein ratio) before being added to a pReViTA DNA template containing the NrdR-sensitive class III RNR promoter from *E. coli* (P*nrdD*). Standard *in vitro* transcription, DNAse treatment, qRT-PCR, and data normalization steps followed (Figure 7B).

Early ReViTA experiments demonstrated that NrdR repression of RNR transcription could be robustly detected and quantified. Nevertheless, at high protein:template ratios (e.g., EMSA conditions), nonspecific inhibition of transcription occurred, so a synthetic, promoter-free DNA analogue (poly-deoxy-inosinic-deoxy-cytidylic acid, poly d[I-C]) was introduced as a promoter-free competitor to mitigate these effects at ratios above 200:1 protein:DNA.

NrdR2^ECO^ (Figure 7D) displayed specific repression when bound to dATP, reducing class III RNR transcription to 54.5% and 34.8% at 2:1 and 4:1 protein:DNA ratios across three independent experiments. ATP-bound NrdR, by contrast, exhibited minimal repression under these conditions. At higher ratios, both dATP and ATP-bound NrdR2^ECO^ repressed transcription, though dATP achieved greater repression. Importantly, at NrdR ratios where repression was dATP-dependent (2:1, 4:1), transcription was never entirely silenced (the same was observed at ratios 6:1, 8:1, 10:1, and 20:1; data now shown), which is consistent with NrdR’s hypothesized role in modulating RNR transcription rather than completely inhibiting it.

ReViTA experiments for NrdR2^PAO^ were not performed due to the large protein quantities required. However, NrdR^PAO^-H6 (Figure 7E) yielded comparable results, although higher protein concentrations were required: dATP-dependent repression was observed at 2000:1 and 4000:1 protein:DNA ratios, reducing transcription to 45.7% and 49.8%, respectively, across three independent experiments. ATP or AMP-bound NrdR^PAO^-H6 caused no significant repression. Similar to its *E. coli* counterpart, no concentration of NrdR^PAO^-H6 completely silenced RNR transcription.

To our knowledge, these experiments provide the first direct observation of NrdR performing its hypothesized biological function: specifically repressing RNR transcription in a dATP-dependent manner, thereby reducing RNR activity under high deoxyribonucleotide levels.

### 2.7. *In vivo* effects of altered NrdR expression

One of the main driving forces behind the study of NrdR is its potential as a target for antimicrobial therapies, given its role as a bacteria-specific master regulator of an essential pathway. However, previous studies in *E. coli* [28], *P. aeruginosa* [20], and *Streptococcus pyogenes* [36] have shown that *nrdR* mutant strains present no significant difference in adherence or infectivity despite significant gene dysregulation. Nevertheless, a more detailed analysis of the impact altering NrdR levels has on the fitness and virulence of *E. coli* [37] demonstrated that, while reduced *nrdR* expression had negligible effects, *nrdR* overexpression let to slower growth rates, reduced proliferation and fitness, and diminished adherence to human epithelial cells. In this study, we extend these findings to *P. aeruginosa*.

First, we compared the growth rate of *P. aeruginosa* PAO1 wild-type, a mutant strain with a fully interrupted *nrdR* gene (Δ*nrdR*) and a complementation strain expressing the pUCP20::*nrdR* plasmid (Δ*nrdR* + Rc), which results in slight constitutive overexpression of *nrdR*. The latter presented an elongated lag phase and a significantly lower growth rate after five hours of culture, ultimately plateauing at a lower optical density (Supplementary Figure S8A).

To evaluate the impact of increased *nrdR* expression on *P. aeruginosa* virulence, we conducted a series of infection tests using *Galleria mellonella*. In two independent experiments, we infected ten larvae per condition, injecting cell suspensions standardized by OD to an average dose of 12 CFU/larva. The results (Supplementary Figure S8B) revealed the striking effect of altered *nrdR* expression levels. In the experiment shown, while the wild-type strain killed its first larva 15 hours post-infection and killed the whole group after 18 hours, all larvae infected with the Δ*nrdR* and Δ*nrdR* + Rc remained alive at that time. These strains required over 24 hours to kill half the larvae, and the overexpression strain was unable to kill all remaining individuals even after 36 hours.

These findings confirm that, in the opportunistic pathogen *P. aeruginosa*, overexpression of *nrdR* significantly reduces bacterial fitness and virulence.

## 3. Discussion

Since its initial description, NrdR has been proposed as a nucleotide-modulated transcriptional regulator of ribonucleotide reductases (RNRs) [14,15]. It is found in almost all bacterial genomes but has no orthologues among eukaryotes or archaea [14]. Several assumptions stem from the location of its cis-elements: NrdR-boxes overlap core promoter elements, indicating transcriptional repression [14,15]. Species encoding NrdR have binding sites upstream of all RNR operons [14,15], indicating a universal role. NrdR-boxes in RNR genes are found in pairs separated by an integer number of DNA helix turns [14], suggesting protein-protein interactions similar to those mediated by ATP-cones in RNRs [2]. The potential of NrdR as a bacteria-exclusive, universal regulator of an essential pathway has made it a candidate target for antimicrobial therapies.

While multiple studies have explored the transcriptomic and proteomic effects of *nrdR* inactivation [20,28,29], the only published search for NrdR-boxes remains its original description [14]. Our genome-wide searches identified all known NrdR-boxes (Supplementary Table 4), along with 42 putative boxes in *E. coli* and 18 in *P. aeruginosa*. Only RNR operons presented pairs of NrdR-boxes separated by an integer number of DNA helix turns, suggesting that the proposed NrdR oligomerization mechanism is specific to RNRs. Other putative NrdR-boxes may indicate an expanded NrdR regulon requiring a different binding mechanism; however, we suspect most are false positives. An equivalent search on randomly generated DNA queries with the same GC-content and number of sequences yielded more hits than the real sequences (Supplementary Figure S10). However, reducing the p-value threshold for whole genome queries resulted in the search missing known boxes. Thus, we concluded that motif-based sequence identification of NrdR-boxes requires correlation with transcriptomics data.

This study presents transcriptomics experiments in *E. coli* and *P. aeruginosa* that confirmed the upregulation of all RNR operons (Supplementary Tables 3 and 5). Other differentially expressed genes were identified, but their correlation with the NrdR-box search revealed that only RNRs were associated with pairs of boxes (Figure 1). However, given the overlap with previous observations [20,28,29], we believe that the absence of NrdR-boxes in other differentially expressed genes does not indicate false positives but rather genes dysregulated by global effects of *nrdR* inactivation, such as reduced fitness and imbalanced NTP/dNTP levels. A comprehensive exploration of the transcriptomic effects of altered NrdR activity was beyond the scope of this study, as it would require comparative RNA-Seq experiments in multiple species under *nrdR* deletion and overexpression conditions. Nevertheless, a limited analysis of our results reveals intriguing trends. Enriched GO biological processes among upregulated genes (Supplementary Figure S9A) included stress-related processes, such as heat response in both species (e.g., chaperone genes *dnaK*, *dnaJ*, *hptG*, *clpB*), proteolysis in *P. aeruginosa* (*lon*, *grepE*, *hslU*) and nucleotide salvage in *E. coli* (*gsk*, *tdk*, *udk*, *apt*, *gpt*). Iron transport and siderophore biosynthesis genes were upregulated in both species, though pyoverdine synthesis specifically was downregulated in *P. aeruginosa*. Other downregulated processes included multiple general anabolic pathways and anaerobic respiration in *E. coli*. A comparison of differentially expressed orthologs between the species (Supplementary Figure S9B) revealed upregulated ribonucleotide reductases, chaperones, heat-shock proteins, and siderophore synthesis proteins for enterobactin (*E. coli*) and pyochelin (*P. aeruginosa*).

The effects of *nrdR* inactivation align with the observations of Wozniak *et al.,* who found that *Bacillus subtilis* Δ*nrdR* shares nearly 60% of differentially expressed genes with RNR enzymatic repression using hydroxyurea [38]. This comprehensive dysregulation contrasts with reports of unchanged infectivity in Δ*nrdR* strains in different infection models [20,28,36]. However, more recently, overexpression of NrdR in *E. coli* was shown to reduce fitness, growth rate, and adherence to human epithelial cells [37]. Our findings extend these observations to *P. aeruginosa*: *nrdR* deletion and overexpression reduced growth rate and infectivity in *Galleria mellonella*, with the latter rendering the strain unable to kill all larvae even after 36 hours (Supplementary Figure S8). We agree with Naveen and Hsiao [37] that therapeutic strategies targeting NrdR should focus on overactivation rather than inhibition. These findings underscore the importance of understanding NrdR’s mechanism of action and structure to design targeted therapeutic strategies.

NrdR oligomerization has been gradually elucidated. Initially, it was determined that NrdR forms dimers without bound nucleotides and higher-order oligomers with nucleoside triphosphates [39]. McKethan and Spiro studied *E. coli* NrdR nucleotide-binding and oligomerization [22], detecting associations up to 20-mers and differences between ATP and dATP complexes. However, they did not establish a fixed stoichiometry for NrdR oligomers, and only detected DNA binding with NrdR-AMP at very high protein:DNA ratios, up to 28000:1. More recently, Grinberg *et al*. offered the first structural information [19,25], revealing that *S. coelicolor* and *E. coli* NrdR ATP-cones bind two nucleotides (as seen in *P. aeruginosa* NrdA [40]) and display tetramer formation when bound to nucleoside triphosphates. With ATP NrdR forms inactive tetramers that further multimerize; in *S. coelicolor*, up to 12-mers have been reported, while *E. coli* NrdR multimerizes into a filament form. When binding ATP and dATP together, it forms active tetramers, which, in *S. coelicolor*, additionally multimerize up to octamers. Our findings are interpreted in the context of these discoveries.

No studies have confirmed the activity differences of NrdR bound to different cofactors, and the protein has repeatedly been reported as highly unstable during purification [15–18,22,27]. Here, we aimed to bridge the gap between structure and function and establish a mechanism of action using stabilized *E. coli* and *P. aeruginosa* fusion proteins (Supplementary Figure 1). NrdR exists as a dynamic population of oligomers, but it was possible to associate cofactors with specific quaternary structure changes and identify their functional differences both *in vivo* and *in vitro*.

When bound to ATP, *E. coli* NrdR formed large multimers visible by atomic force microscopy (Figure 2) and size-exclusion chromatography (Supplementary Figure 3). SEC-MALS confirmed molecular weights starting at tetramers (∼75-80 kDa) and averaging at octamers (Figure 3A, Table 1); the high end of the range shifted towards heavier complexes when NrdR was pre-incubated with ATP (∼180 kDa to 260 kDa, for 10 to 16-mer) (Figure 3B, Table 1). Interestingly, the lower end remained constant regardless of ATP incubation time, suggesting a stable form (tetramer) and higher possible multimerization. While Grinberg *et al.* [19] only reported up to 12-mers for *S. coelicolor* NrdR-ATP, our results align *E. coli* NrdR data showing 16-mers and higher multimers [22,25], which suggests that the reported differences in multimerization are not merely due to variations in the experimental setup but actually reflect distinct particularities of NrdR in different species. NrdR-ATP displayed no DNA binding activity in the presence of competition DNA (Figure 7A, Supplementary Figure S7) and, crucially, did not repress RNR transcription *in vitro* at low concentrations (Figure 7D).

When bound to dATP, *E. coli* NrdR formed complexes determined by size-exclusion chromatography as larger than dimers but smaller than NrdR-ATP (Supplementary Figure 3). Using atomic force microscopy, complexes were observed as slightly smaller than ATP-bound complexes and with a distinctive lower circularity index (Figure 2). These observations align with *Grinberg et al*. cryo-EM and crystal structures, which show tetramers containing dATP as more elongated than their ATP-bound counterparts. SEC-MALS confirmed molecular weights ranging from tetramer to octamer, which did not change over time upon dATP pre-incubation (Figure 3, Table 1). These elongated complexes displayed specific DNA-binding activity (Figure 7A) and repressed RNR transcription at ratios at which NrdR-ATP was inactive (Figure 7D).

We propose that our data represents the fundamental active and inactive states of *E. coli* NrdR, similar to the described structurally in *S. coelicolor* and *E. coli* [19,25]. Despite the limited availability of recombinant *P. aeruginosa* NrdR, the same observations concerning ATP/dATP-bound NrdR can be extended to this species: the protein presented equivalent oligomer sizes on SEC experiments (Supplementary Figure S3B) and was demonstrated to be only active when bound to dATP at both DNA binding (Supplementary Figure 7) and *in vitro* transcription levels (Figure 7E). Given the recent discovery that NrdR simultaneously needs dATP and ATP/ADP to bind DNA [19,25], it now becomes apparent that the activity we detected may have relied on traces of ATP or ADP in as-prepared proteins, which have been found using PCA precipitation and HPLC (Supplementary Figure S2) and on the crystal structure (Figure 5). Incidentally, the inactivity of dATP-bound NrdR in *E. coli* observed by McKethan and Spiro [22] is also explainable this way: as part of that study, NrdR was deprived of all nucleotides using controlled denaturation. Future studies should explore the *in vitro* activity of NrdR in the presence of different combinations of dATP and ribonucleotides, physiological nucleotide mixtures, or cell extracts to replicate biological conditions.

These discoveries expand upon the known structural information and, through functional experiments, link it to a complete mechanism of action (Figure 8). In this model, NrdR exists as a dimer only when bound to monophosphates or without cofactors, representing a transient state. *In vivo,* NrdR is found in equilibrium between an “inactive” ATP-only tetramer and a more elongated “active” tetramer containing dATP (and presumably ribonucleotides). The active tetramer can form octamers, while the inactive one can multimerize into octamers, 12-mers, 16-mers, or higher-order structures. The balance between active and inactive NrdR is modulated by the cellular dNTP:NTP ratio. Despite the substantial difference in the physiological concentrations of ribonucleotides and deoxyribonucleotides [23,24], dATP binding is facilitated by the described negative cooperative nucleotide-binding mechanism [22]. Exploiting this system for antimicrobial therapies would likely require inducing NrdR tetramers to remain in their active state. However, given the universality of nucleotides, dNTP analogs targeting the ATP-cone would likely affect other systems in both bacteria and host cells, rendering these compounds unsuitable for therapy. Detailed structural insights into NrdR with different cofactors could provide invaluable strategies for targeting this system. For example, exploiting both ATP-cone binding sites simultaneously or acting directly on residues involved in protein-protein contacts to alter NrdR oligomerization could be promising approaches.

**Figure 8.**
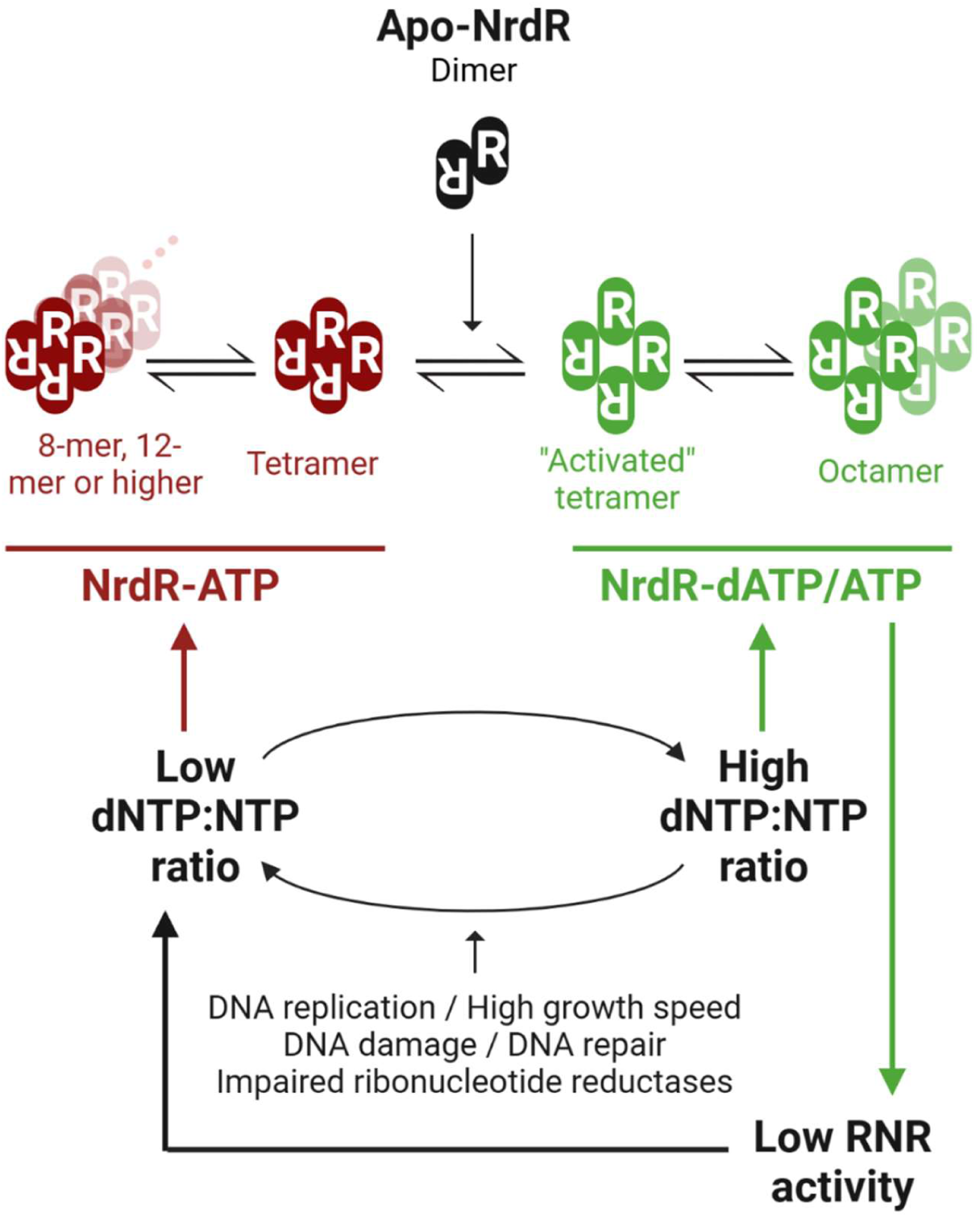
Model of the NrdR mechanism of action. NrdR without a cofactor is dimeric. When bound only to ATP in the presence of a low dNTP:NTP ratio, it forms “inactive” tetramers that multimerize into octamers, 12-mer or higher-order oligomers as an inactive storage state. When bound to dATP along with ATP/ADP in the presence of higher dNTP concentrations, the tetramers adopt an “activated” form, which can assemble into octamers. Activated tetramers repress RNR transcription, thereby reducing the dNTP:NTP ratio. The arrangement of NrdR proteins depicted is for illustration purposes and does not represent actual quaternary structures. Created in BioRender (Pedraz, L., 2025 https://BioRender.com/l48c214).

Here, we provide a crystal structure of *E. coli* NrdR obtained without adding nucleotides to the as-prepared protein (Figure 4, Supplementary Figure S5, PDB ID: 9GEX), and which presents ATP, ADP, and AMP bound to the “inner” ATP-cone site of the different monomers in the asymmetric unit. Efforts to crystalize *E. coli* NrdR-ATP NrdR-dATP tetramers were unsuccessful, potentially due to the high variability in multimerization. A potential strategy to obtain these structures could involve co-crystallizing NrdR with its target DNA in the presence of dATP/ATP mixtures.

Our structure provides detailed insights into the interactions required for cofactor binding at the inner site of the ATP-cone (Figure 5). Without providing additional nucleotides, the outer site was predictably empty in all monomers. In *S. coelicolor*, Tyr128 was described as essential for nucleotide binding and ATP/dATP discrimination at the outer site [19]. This residue has two rotamers: one oriented towards helix α1 when ATP is bound at the outer site and one towards α3 with dATP bound. In our structure, the equivalent Tyr131 displays similar behaviour, interacting with Ser105 in the α3 conformation and with Arg72 in the α1 conformation. The positive charge of Arg72 (or Lys69 in *S. coelicolor*) may stabilize Tyr131 in the absence of nucleotides, such as during ATP/dATP exchange. Mutation Y131A drastically reduced NrdR oligomerization beyond dimers (Figure 6B) but did not render NrdR inactive (Figure 6C), indicating that disrupting Tyr131 alone is insufficient to inactivate NrdR *in vivo*.

We describe the asymmetric unit of the *E. coli* NrdR crystal structure as a pair of interacting dimers, as the interactions observed between molecules A-B and C-D were determined to be stable (based on the estimated complex formation ΔG and contact area) while the inter-dimer AB-CD interactions were weak (Figure 4A). This dimer pair’s geometry is incompatible with DNA binding, as it does not fit NrdR-box arrangements described so far. The dimensions of the complex (approx. maximum length of the dimer 7.4 nm; 11.6 nm for the pair of dimers) demonstrate that the structures previously observed by atomic force microscopy (80% of particles between 8 and 20 nm in length; Figure 2B) are discrete NrdR oligomers, validating the technique for the study of NrdR quaternary structure.

The inter-dimer contacts observed involve Arg28 and Arg37 interacting with Glu36, a residue not conserved beyond Proteobacteria (Supplementary Figure S6). In *S. coelicolor*, an arginine occupies the equivalent position [19], preventing these interactions unless arginine stacking occurred, but none of the available NrdR cryo-EM structures show such a contact. Mutation E36A altered NrdR oligomerization (Figure 6B) but did not prevent NrdR repression (Figure 6C). Similarly, disrupting α4-α5 interactions between symmetry partners in the crystal (Figure 4B) by deleting helix α5 and the preceding loop resulted in drastic oligomerization changes (Figure 6B). Superimposition of α4 from our *E. coli* and Grinberg’s *S. coelicolor* structures (PDB ID:7P37, 7P3Q, 7P3F [19]), however, reveals that α4-α5 contacts are conserved in *S. coelicolor*, and key residues are conserved, suggesting this interaction is biologically relevant. The surprising contrast between the drastically altered oligomer populations of some mutant NrdR proteins and the lack of changes *in vivo* highlights the challenges still to overcome when replicating NrdR functionality *in vitro*: despite not showing it under SEC-MALS conditions, mutants E36A and Y131A must be able to form active tetramers in the cell, preserving NrdR activity. As suggested above, future studies using physiological nucleotide concentrations or whole cell extracts are needed to replicate the *in vivo* environment of the protein.

Encouragingly, disrupting intra-dimer contacts showed significant *in vivo* effects. This contact is mediated by Arg27, Arg29, and Glu42 (Figure 4A). Mutation E42A nearly disabled oligomerization, rending *E. coli* NrdR predominantly monomeric (Figure 6B) and abolishing RNR transcriptional repression (Figure 6C). These residues are fully conserved (Supplementary Figure S6), and equivalent interactions in *S. coelicolor* are evident in a structural superposition with the cryo-EM data [19]. This intra-dimer interface appears essential for NrdR functionality, with Glu42 also potentially critical for the proper folding of NrdR.

Overall, our findings expand on existing structural data, linking different quaternary structures to changes in DNA binding and the capacity to repress RNR transcription *in vitro*. We demonstrated that NrdR’s oligomerization-based repression mechanism is specific to RNR operons using transcriptomics and motif-based NrdR-box identification. Additionally, used the crystal structure of as-prepared NrdR to identify critical elements for nucleotide-binding and protein-protein interaction. We believe future studies should investigate NrdR activity in the presence of *in vivo*-like combinations of ribonucleotides and deoxyribonucleotides, along with pursuing additional NrdR crystal structures with different cofactors and DNA, to develop targeted antimicrobial strategies.

## 4. Materials and Methods

### 4.1. Bacterial strains, plasmids, and growth conditions

Bacterial strains and plasmids are listed in Supplementary Table S6. *Escherichia coli* and *Pseudomonas aeruginosa* cells were routinely grown in Luria-Bertani broth (LB) at 37 °C. When necessary, antibiotics were added at the following concentrations: for *E. coli*, 10 µg/ml gentamicin (Gm^R^), 50 µg/ml ampicillin (Amp^R^), 30 µg/ml kanamycin (Kn^R^); for *P. aeruginosa*, 150 µg/ml gentamicin (Gm^R^), 300 µg/ml carbenicillin (Amp^R^). Liquid cultures were shaken on an orbital shaker at 200 rpm.

For growth curves, the desired strains were used to inoculate overnight cultures (in LB medium, with the required antibiotics). Cells were pelleted by centrifugation (5000 g, 10 minutes) and resuspended in sterile PBS, calculated for an OD_550_ of 1.0. The cultures for the growth curves were prepared with LB medium, the required antibiotics, and cell suspension to a final OD_550_ of 0.05 in 96-well plates. Plates were incubated at 37 °C with orbital shaking and constant humidity in the SPARK Multimode Microplate Reader (TECAN) using the SPARK Small Humidity Cassette (TECAN). OD_550_ was monitored every 20 minutes.

### 4.2. DNA manipulation and plasmid construction

All PCR primers used in this study are listed in Supplementary Table S7 and will be referred to by their numbers. Plasmid DNA was prepared using the GeneJET Plasmid MiniPrep Kit (ThermoFisher), according to the manufacturer’s instructions. Plasmid transformation into *E. coli* was performed using calcium-competent cells, as previously described [41]. Plasmid transformation into *P. aeruginosa* cells was done by electroporation, using a Gene Pulser Xcell electroporator (Bio-Rad), as previously described [42]. Digestion reactions with restriction enzymes followed the manufacturer’s instructions (ThermoFisher). Ligations were performed with the T4 DNA ligase (ThermoFisher). DNA fragments for cloning were obtained by PCR using Phusion High-Fidelity DNA Polymerase (ThermoFisher) unless otherwise stated. During all plasmid construction procedures, fragments synthesized by PCR and digested with restriction enzymes were first cloned via blunt-end cloning to pJET1.2b (CloneJET PCR Cloning Kit, ThermoFisher), according to the manufacturer’s instructions and then digested from the resulting carrier plasmid. Colony PCR reactions to test plasmid incorporation were carried out using DreamTaq Green PCR Master Mix (ThermoFisher). Plasmid cloning was performed in DH5α unless otherwise stated.

Plasmid pET-NrdR(PAO) for the production of NrdR^PAO^-H6 was constructed by amplifying *nrdR* between restriction sites *Nde*I and *Xho*I from genomic *P. aeruginosa* PAO1 DNA using primers 1 and 2 and cloning the purified PCR amplicon into pET22b^+^ as described above. The incorporation of the insert was tested by colony PCR using primers 3 and 4.

Plasmids pSUMO-NrdR(PAO) and pSUMO-NrdR(ECO) for the production of NrdR1^PAO^ and NrdR1^ECO^ were constructed by amplifying *nrdR* from genomic *P. aeruginosa* PAO1 and *E. coli* K-12 MG1655 DNA using primer pairs 5-6 and 7-8, respectively. Primers 5 and 7 add a small two-amino acid linker (GS) to the N-terminus of NrdR to optimize SUMO cleavage. The resulting fragments were inserted into plasmid pAviTag-NN-His SUMO Kan (pSUMO) using a recombineering procedure following the manufacturer’s instructions (Expresso Biotin SUMO Cloning and Expression System, Lucigen). The translational fusion in the final constructs resulted in 6His-AviTag-SUMO-(GS)-NrdR proteins. The incorporation of the inserts was tested by colony PCR using primers 9 and 10.

Plasmids pCri-NrdR(PAO) and pCri-NrdR(ECO) for the production of NrdR2^PAO^ and NrdR2^ECO^ were constructed by amplifying *nrdR* between restriction sites *Nco*I and *Xho*I from genomic *P. aeruginosa* PAO1 and *E. coli* K-12 MG1655 DNA using primer pairs 11-12 and 13-14, respectively. Primers 11 and 13 add a TEV protease cleavage site (TEVcs, ENLYPQG) and a five amino acid linker (SGSGS) to the N-terminus of NrdR, as SUMO cleavage was inefficient. The resulting fragments were cloned into pCri11a [43] as described above. The translation fusion in the final constructs resulted in 6His-SUMO-TEVcs-(SGSGS)-NrdR proteins. The incorporation of the inserts was tested by colony PCR using primers 3 and 4.

pCri-NrdR(ECO) was used as a template to generate NrdR2ECO mutant derivatives by the “round the horn” PCR method [44] using Herculase DNA II Polymerase (Agilent) and primer pairs 29-30 (mutation E36A), 31-32 (mutation E42A), 33-34 (mutation Y131A), resulting in plasmids pCri-NrdR-E36A, pCri-NrdR-E42A, and pCri-NrdR-Y131A, respectively. Plasmid pCri-NrdRΔ132, producing a truncated version of NrdR2^ECO^ with amino acids 132-149 deleted, was obtained by amplifying the NrdR_2_^ECO^ construct from pCri-NrdR(ECO) using primers 35-36 using PfuUltra DNA Polymerase (Agilent) and recloning the resulting amplicon into pCri11a as described above.

Plasmid pReViTA-0 was constructed as an initial version of the more advanced iteration described by Rubio-Canalejas *et al.* [35] (GenBank OP909926). To construct pReViTA-0, the backbone from plasmid pETS130 [45] was amplified by PCR and obtained between restriction sites *Xba*I and *Aat*II using primers 15 and 16. The ReViTA cassette was synthesized *de novo* as a GeneArt Gene Synthesis product (ThermoFisher) containing, among other components, the sequence of a non-functional truncated *cat* gene [45], constitutive promoter J23119 (BBA_J23119, Registry of Standard Biological Parts), and transcription terminator B0015 (BBA_B0015, Registry of Standard Biological Parts). The ReViTA cassette and the pETS130 backbone fragment were digested with *Aat*II and *Xba*I, and ligation was performed to obtain pReViTA-0. A schematic of the pReViTA-0 plasmid is available in Figure 7C; its full sequence is available upon request. To obtain the derivative plasmid pReViTA-PD, containing the NrdR-sensitive P*nrdD* promoter, the promoter of the *nrdDG* operon was amplified from genomic *E. coli* K-12 *substr*. MG1655 DNA between restriction sites *Bam*HI and *Cla*I using primers 17 and 18, and the purified PCR amplicon was cloned into pReViTA-0 as described above. The incorporation of the insert was tested by colony PCR using primers 17 and 23.

Plasmid pLG338-NrdR at its derivatives carrying mutant *nrdR* genes used to complement K-12 Δ*nrdR* [46] were commercially produced as VectorArk System products (GenScript); they carry the backbone of low copy number plasmid pLG338/30 [47] and *nrdR* from *E. coli* K-12 *substr*. MG1655 (wild-type or its mutant derivatives) under the control of its own promoter region (90 bp upstream of the *nrdR* translation start codon, including a known Sigma70 cis-element [18]). The full sequence of these plasmids is available upon request.

### 4.3. Bioinformatic prediction of NrdR-boxes

To perform genome-wide searches for NrdR-boxes, we started by retrieving the sequence and annotation for the *E. coli* K-12 substr. MG1655 genome (NCBI Reference Sequence: NC_000913.3) and the *P. aeruginosa* PAO1 genome (NCBI Reference Sequence: NC_002516.2). We then extracted only the “gene” features from the annotation. Using gene features as a reference, we retrieved the sequences starting 450 bp upstream and ending 20 bp downstream of every translation start codon in the genome using the SAMtools [48] suite and the BEDtools [49] functions flank, slop, and getfasta. A comprehensive shell script for this process is available upon request.

A consensus sequence for the NrdR-box was obtained using the γ-proteobacteria data published by Rodionov and Gelfand [14], representing 135 NrdR-box sequences. First, we used MEME [50] (MEME Suite [31]) to produce a weight matrix. Then, FIMO [30] (MEME Suite [31]) searches were performed, applying a 10^-4^ p-value threshold. Results were deduplicated and assigned to the gene with the closest translation start. Hits inside non-coding RNAs and low-complexity regions were ignored.

### 4.4. Transcriptomics and transcriptomics data treatment

To obtain RNA samples for RNA-seq, we used three independent cultures of *P. aeruginosa* PAO1 in 25 ml LB medium and three of its isogenic mutant strain PAO1 Δ*nrdR* [51] in 25 ml LB medium supplemented with 40 µg/ml tetracycline. Cultures were grown to an OD_550_ of 0.5 and fixed using RNAprotect Bacteria Reagent (QIAGEN) according to the manufacturer’s instructions. Then, RNA was extracted using the RNEasy Mini RNA Isolation Kit (QIAGEN), according to the manufacturer’s instructions. Reverse transcription, library generation, and RNA sequencing were performed by Beckman Coulter Genomics, according to the protocol “Illumina TruSeq Stranded Total RNA with Ribo-Zero rRNA Removal (Bacteria)”. The platform used was Illumina 1.9, with the library TruSeq3-PE-2. 6 samples were analyzed, and 362 million paired-end reads were generated (2 x 100 bp each).

RNA-seq data were received as untreated sequence and quality data (FASTQ). To remove adapter sequences and low-quality bases, a first FASTQ trimming step was introduced using Trimmomatic version 0.36 (LEADING:5, TRAILING:5, SLIDINGWINDOW 4:15, MINLEN:25, LLUMINACLIP:/RNA/REF/TruSeq3-PE-2.fa:2:30:10:2:true) [52]. Data was then mapped using end-to-end alignment with bowtie version 1.5, allowing for multiple binding. Mapping parameters were: -S -t --fr -n 2 -l 28 -e 70 -k 5 --best --strata --allow-contain --no-unal –nomaqround [53]. The *P. aeruginosa* PAO1 genome was used as reference (NCBI Reference Sequence NC_002516.2). The output SAM file was converted to BAM and sorted using SAMtools functions “view” and “sort”, with default parameters [48]. The quality of the mapped data was assessed using Qualimap 2.2.1 [54]. Differentially expressed genes were obtained using DESeq2 version 3.9 (R, Bioconductor) according to the general pipeline [55–57]. Only genes with an absolute fold-change over 2.0 were considered unless otherwise stated.

Enriched GO IDs were determined using PANTHER [58,59], summarized and analyzed using REVIGO [60] and plotted using R Treemap [61]. Orthologous genes for cross-comparisons of *E. coli* and *P. aeruginosa* were obtained from OrtholugeDB [62].

### 4.5. GFP-based reporter gene assays

*E. coli* K-12 Δ*nrdR* was transformed with low-copy number plasmid pLG338 carrying either a wild-type *nrdR* gene or its mutant derivatives E36A, E42A, and Y131A. All those strains and uncomplemented K-12 Δ*nrdR* were subsequently transformed with plasmid pETS130-PA, carrying GFP controlled by the promoter region of class Ia RNR. Cultures were grown overnight on LB with all the required antibiotics. 25 ml cultures on LB without antibiotics were then inoculated to an initial OD_550_ of 0.05 and incubated at 37 °C. Cultures were sampled at OD_550_ of 0.50 (exponential phase) and 2.50 (stationary phase). Three independent 1 ml samples were collected per strain and sampling point. Samples were centrifuged for 10 minutes at 5000 g; supernatants were discarded, and pellets were fixed with 1 ml 1x PBS + 2% formaldehyde. Samples were incubated for 10 minutes at 4 °C, protected from light, centrifuged again for 10 minutes at 5000 g, and pellets were resuspended in 1x PBS. Fluorescence was determined using 96-well plates (Costar 96-Well Black Polystyrene Plate, Corning) in an Infinite 200 Pro Fluorescence Microplate Reader (Tecan).

### 4.6. NrdR overexpression and purification

The NrdR^PAO^-H_6_ protein was overexpressed in BL21(DE3) transformed with plasmid pET-NrdR(PAO). Cells were grown in LB medium with 50 µg/ml ampicillin and 100 µM ZnSO_4_ and incubated at 37 °C with vigorous shaking. When cultures reached an OD_550_ of 0.5, isopropyl β-D-1-thiogalactopyranoside (IPTG) was added to 0.1 mM and cells were cultured at 18 °C overnight. Cells were pelleted by centrifugation (6000 g, 20 min, 4 °C) and resuspended in 25 ml (per liter of original culture) of lysis buffer: 50 mM Tris-HCl (pH 8.5 at 25 °C), 1 M NaCl, 20 mM imidazole, 1 mM phenylmethylsulphonyl fluoride (PMSF), 100 µM ZnSO_4_, and 10% glycerol. The suspension was then sonicated until clear (20 pulses of 20 s, with 50 s cooldown between pulses, using a 1/2’’ tip in a Branson 450 Digital Sonifier, Marshal Scientific) and centrifuged at high speed (15000 g, 20 min, 4 °C), keeping the supernatant, which was frozen at -80 °C for long term storage. NrdR was then purified from the extracts by Immobilized Metal Affinity Chromatography (IMAC) using a 5 ml Mini-Nuvia IMAC Cartridge (Bio-Rad) in an FPLC system (Biologic DuoFlow System, Bio-Rad). Protein samples suffered a DNA precipitation step (30 minutes incubation with streptomycin sulfate 1%, at 4 °C) and were diluted with buffer A to a concentration of less than 1 mg/ml of total protein before being injected into the column. The column was equilibrated with 5 column volumes (CV) of Buffer A (50 mM Tris-HCl pH 8.5, 1 M NaCl, 20 mM imidazole). A washing step was carried out using 5 CV of Buffer A. Buffer B (50 mM Tris-HCl pH 8.5, 1 M NaCl, 500 mM imidazole) was then mixed in at different proportions to start the elution. First, contaminant proteins were removed with a non-specific elution step using 5 CV of buffer with 50 mM imidazole. Then, the protein was recovered in a specific elution step using 5 CV of buffer with 200 mM imidazole. The resulting fractions were analyzed using SDS-PAGE. Fractions containing the protein of interest were concentrated and diafiltrated against buffer 5x NrdR: 100 mM Tris-HCl (pH 9.0 at 25 °C), 400 mM KCl, 5 mM MgCl2, 5 mM DTT, 250 µM ZnSO_4_, and 25% glycerol, using VivaSpin 20 10000 MwCO Ultrafiltration units (Sartorius).

The NrdR_1_^ECO^ and NrdR_1_^PAO^ proteins were overexpressed in XCell F’ Chemically Competent Cells (Expresso Biotin SUMO Cloning and Protein Expression System, Lucigen) transformed with plasmids pSUMO-NrdR(ECO) and pSUMO-NrdR(PAO), respectively. Cells were grown in 1-litre cultures of LB medium supplemented with 30 µg/ml kanamycin and 50 µM ZnSO_4_ and incubated at 37 °C with vigorous shaking. When cultures reached an OD_550_ of 1.0, protein overexpression was induced by adding 0.1% rhamnose. Cells were then cultured at 18 °C overnight. Cells were pelleted, resuspended, sonicated, and centrifuged as for NrdR-H_6_. The crude extracts then suffered a first purification step using the same procedure described above for NrdR-H_6_. The resulting fractions were analyzed using SDS-PAGE. Fractions containing the protein of interest were concentrated and diafiltrated against buffer 4x NrdR-PROT: 100 mM Tris-HCl (pH 8.5 at 25 °C), 600 mM NaCl, 4 mM DTT, 40 µM ZnSO_4_ using VivaSpin 20 10000 MwCO Ultrafiltration units (Sartorius). Before SUMO protease digestion, the protein was diluted with water to a final concentration of 1x NrdR-PROT buffer, and +2 mM fresh DTT was added. SUMO protease (Lucigen) was added (1 unit for each 300 µg of protein), and the reaction mixture was incubated for 3 hours at 30 °C with gentle mixing. Extra DTT was removed from the digested protein mixture (but not completely, to avoid protein precipitation) by dialysis against 50 mM Tris-HCl (pH 9.0 at 25 °C), 1 M NaCl, 0.5 mM DTT, using Slide-a-Lyzer Dialysis Cassettes (ThermoFisher). The digested protein then suffered a second purification step by a negative IMAC in the same column and system mentioned above. First, the column was equilibrated with 5 CV of 50 mM Tris-HCl pH 9.0, 1 M NaCl, 20 mM imidazole. Protein samples were then injected into the column. The protein was recovered from the flow-through, as the SUMO-tag and SUMO-protease were retained. The resulting fractions were analyzed using SDS-PAGE. Fractions containing the protein of interest were concentrated and diafiltrated as described above against buffer 5x NrdR.

The NrdR_2_^ECO^ and NrdR_2_^PAO^ proteins were overexpressed in BL21(DE3) transformed with plasmids pCri-NrdR(ECO) and pCri-NrdR(PAO), respectively. Cells were grown overnight until saturation in 40 ml pre-cultures of LB with 30 μg/ml kanamycin, which were inoculated into 750 ml LB cultures containing 30 μg/ml kanamycin and 200 μM ZnSO_4_. When cultures reached an OD_600_ of 0.6, expression was induced by adding IPTG to 0.2 mM. Cells were then cultured at 18 °C overnight. Cells were spun down by centrifugation for 40 minutes at 4.000 rpm; bacterial pellets were flash-frozen in liquid nitrogen and stored at -80°C. Pellets were resuspended in 30 ml of lysis buffer (50 mM Tris-HCl pH 9.0 at 25 °C, 1 M NaCl, 50 μM ZnSO4, 1 mM DTT, 10 U/ml DNase I, 10% v/v glycerol, 1 mM PMSF) complemented with Complete EDTA-Free antiprotease tablets (Roche). After that, cells were sonicated for 3 minutes (cycling 3 s ON, 2 s OFF at 20% power, using a 1/2’’ tip in a Branson 450 Digital Sonifier, Marshal Scientific). After the addition of streptomycin sulphate to a final concentration of 0.1% to precipitate DNA and nucleotide traces, the lysed mixture was stirred for 45 min at 4 °C and centrifuged for 25 minutes at 48,400 g (Beckman Coulter centrifuge, rotor JA 25-50, 20.000 rpm). The supernatant was filtered with a syringe filter (pore size of 0.22 μm; Merck Millipore) and loaded onto a HisTrap 5 ml nickel column (Cytiva) connected to an FPLC Åkta system and washed with 5 CV of buffer A (1 M NaCl, 50 mM Tris-HCl pH 9.0 at 25 °C, 20 mM imidazole). A 10 ml single step at 200 mM imidazole was performed for protein elution by mixing in buffer B (1 M NaCl, 50 mM Tris-HCl pH 9.0 at 25 °C, 500 mM imidazole). The eluting fractions (2 ml) were analyzed by SDS-PAGE and pooled. Efficient TEV digestion required buffer exchange to 50 mM Tris-HCl pH 9.0, 500 mM NaCl and 1 mM DTT using VivaSpin 20 3000 MwCO Ultrafiltration units (Sartorius). TEV digestion was at a protein:protease molar ratio of 25:1, at 4°C overnight, and analyzed by 15% SDS-PAGE. The nature of the bands was verified using MALDI-TOF Mass Spectrometry (CIB Margarita Salas-CSIC, Madrid). After digestion, a Ni-NTA negative chromatography was run, GSGSGS-NrdR_2_ eluted in the flow-through, and 6His-SUMO-ENLYPQ, traces of TEV protease and traces of undigested fusion were retained in the column. Highly pure GSGSGS-NrdR_2_ was concentrated using the same ultrafiltration units as above to 500 µl at a concentration of 16 mg/ml. A high 260/280 nm absorbance ratio (∼0.9) suggested a nucleotide strongly bound to NrdR, which was >95% pure as assessed by 15% SDS-PAGE, optimal for SEC-MALS and crystallization.

Expression and purification of NrdR_2_^ECO^ mutant derivatives E36A, E42A, Y131A, and Δ132-149 followed the same protocol as for wild-type NrdR_2_^ECO^ using BL21(DE3) transformed with pCri-NrdR-E36A, pCri-NrdR-E42A, pCri-NrdR-Y131A, and pCri-NrdR-Δ132, respectively.

### 4.7. HPLC for nucleotide quantification

Ion-pair reverse-phase HPLC was used to quantify nucleotides bound to NrdR^PAO^-H_6_ and NrdR_2_^ECO^, both to determine the nucleotides bound to as-prepared protein extracts and to establish their capacity to bind other nucleotides in solution. 40 µl reactions were prepared in 1x buffer NrdR (see above) containing 100 µg of protein and, if required, enough nucleotide for a 20:1 nucleotide:protein molar proportion. Samples were incubated for 1 hour at room temperature, protected from light. After incubation, free-nucleotides were removed using Zeba Spin Desalting Columns, 0.5 ml, 7K MwCO (ThermoFisher), according to the manufacturer’s instructions; the desalting process was proven to eliminate around 98% of unbound nucleotide (see Supplementary Figure 2C). After protein elution, protein-cofactor complexes were mixed in with pH indicator bromocresol green (Millipore Sigma) to a final 0.04% w/v and perchloric acid (PCA) to a final 10% w/v for protein precipitation. Reactions were incubated for 10 minutes at room temperature, protected from light, and then centrifuged for 5 min at 16000 g, 4 °C, to pellet the precipitated protein, leaving any cofactor that used to be bound to NrdR free in solution. 40 µl of the supernatant were recovered, and potassium hydroxide 4 M was added to reach a pH of 4.5 (when bromocresol green becomes green). Samples were further centrifuged for 5 min at 16000 g, 4 °C to remove any precipitation, and 30 µl of the supernatant was recovered for HPLC quantification.

Nucleotides were quantified using ion-pair reverse-phase HPLC coupled to UV (260 nm) following an established protocol for nucleotide and nucleoside quantification [63] at the Separative Techniques service, University of Barcelona Scientific and Technological Centers (CCiTUB). Solutions containing 10 nmol AMP, dAMP, ADP, ATP, and dATP were prepared using the same PCA precipitation protocol as the protein samples and used as standards for quantification. Protein precipitation supernatants were diluted to 60 µl with mobile phase, and 50 µl were used for injection. Data analysis was performed using Empower Chromatography Data System (Waters).

### 4.8. SEC and SEC-MALS

For SEC (Size-Exclusion Chromatography) experiments, purified NrdR^PAO^-H_6_, NrdR_1_^ECO^, and NrdR_1_^PAO^ were used. For cofactor pre-incubation, 10 mM nucleotide solutions were added directly to the concentrated protein to a final molar ratio of 20:1 nucleotide:protein, and incubated for 1 hour at room temperature. We used a Superdex 200 10/300 GL column (20 ml bed volume) (GE Healthcare) in a BioLogic DuoFlow FPLC System (Bio-Rad). The chromatography was conducted at a fixed flow rate of 0.5 ml/min with the following elution buffer: 50 mM Tris-HCl (pH 9.0 at 25 °C), 250 mM NaCl, 5 mM DTT. When required, nucleotide was added to the column, calculated for a 1:1 molar ratio of nucleotide:protein (taking the sample concentrations as reference). The column was equilibrated with 2 CV of elution buffer before injecting the samples. When the protein-nucleotide complexes were to be run, the column was re-equilibrated with 0.5 CV of elution buffer containing the corresponding nucleotide. Sample concentrations were normalized at 0.5 mg/ml, and a fixed injection volume of 220 µl was used. Data was analyzed using BioLogic DuoFlow Software (Bio-Rad).

For SEC-MALS (SEC coupled with multi-angle light scattering), NrdR_2_^ECO^ and NrdR_2_^PAO^ were used, as well as NrdR_2_^ECO^ mutant derivatives. Experiments were performed at the Barcelona Science Park Automated Crystallization Platform (PAC). Samples were injected to a Superdex 200/10/300 Increase column (Cytiva), equilibrated with 50 mM Tris-HCl (pH 9.0 at 25 °C), 250 mM NaCl, 1 mM DTT, at a flow rate 0.5 ml/min, attached to a HPLC ‘High Prominence Liquid Chromatography’ (Shimadzu), in turn connected to a DAWN-HELEOS-II detector (Wyatt Technology), equipped with a 664.3 nm laser. A first set of experiments were carried out in the same buffer, with 0.025 mM AMP, ADP, dATP or ATP. For preincubated samples (3 h at RT with the NTPs at a protein:nucleotide molar ratio 1:20), NTPs were added to the elution buffer for maximum nucleotide saturation. SEC and MALS data were analyzed with Unicorn 7 (Cytiva) and ASTRA 7 (Wyatt Technology), respectively. Calculations were performed by the PAC staff.

### 4.9. Electrophoretic mobility shift assay

Two DNA probes were used for EMSA: an NrdR-sensitive probe containing the entire promoter region of *nrdAB* from *E. coli* K-12 MG1655 (P*nrdA* ECO) and a negative-control, NrdR-insensitive probe containing an unrelated internal sequence of the *anr* gene from *P. aeruginosa* PAO1 (*anr*). Probes were generated by a double PCR method. A first PCR is conducted to obtain the corresponding fragments with an arbitrary sequence added at the 3’ end of every fragment (M13 complementarity tail); primers 17 and 25 were used for P*nrdA* ECO, and primers 26 and 27 for *anr*. Then, a second PCR uses a WellRED oligo (Millipore Sigma) coupled to the near-infrared fluorophore D3-phosphoramidite (D3-PA); primers 17 and 28 were used for P*nrdA* ECO, and primers 26 and 28 for *anr*. Probes were purified from agarose gels using the GeneJET Gel Extraction Kit (ThermoFisher) and used in EMSA experiments at a fixed quantity of 100 fmol.

Purified NrdR-H_6_, NrdR_1_, or NrdR_2_ proteins were used in DNA-protein binding reactions in total amounts of 0, 200 or 400 pmol per reaction, corresponding to 0, 2000 or 4000 protein:DNA molar ratios. Binding reactions (20 µl) also contained BSA (0.2 µg/reaction) and salmon sperm DNA (2 µg/reaction), as well as 4x NrdR-binding buffer, added to a final 1x concentration of 20 mM Tris-HCl (pH 9.0 at 25 °C), 80 mM KCl, 1 mM MgCl_2_, 1 mM dithiothreitol, 50 µM ZnSO_4_, and 5% glycerol. Reactions were incubated at room temperature for 60 minutes before gel electrophoresis.

Electrophoresis was performed in 4% acrylamide gels prepared with a 37.5:1 proportion of acrylamide:bis-acrylamide; 5% triethylene glycol was used as an additive to increase DNA-protein complex stability. Gels were cast and run using the PROTEAN II xi system (Bio-Rad), according to the manufacturer’s instructions. Gels were run at 25 mA, constant current, 4 °C, for 4 hours. Final images were obtained by scanning the gels in an Odyssey Imaging System device (LI-COR Biosciences) in the 700 nm channel.

### 4.10. Crystallization and Structure Determination

Nano-crystals of GSGSGS-NrdR_2_^ECO^ (16 mg/ml in 500 mM NaCl, 50 mM Tris-HCl pH 9.0 at 25 °C, 1 mM DTT) appeared in a sitting-drop vapor-diffusion crystallization set-up at room temperature (RT), by using 96-well plate sparse matrix screens at the Automated Crystallography Platform (PAC) at the Barcelona Science Park (PCB). Crystals were optimized in 24-well Hampton research plates and fished from drops containing 0.15 M KSCN, 0.1 M Tris-HCl (pH 9.0 at 25 °C), 7% v/w PEG6000 as crystallization solution; crystals were cryoprotected with 20% glycerol and vitrified in liquid nitrogen. X-ray diffraction data were collected at the German Electron Synchrotron (DESY, Hamburg) at the zinc absorption edge (1.28 Å). Three datasets were collected from different crystal regions with a 0.1° oscillation range (totalling 360°), showing nominal resolutions ranging from 2.35 to 2.55 Å. Datasets were processed with XDS package [64] and merged with XSCALE. Data resolution ranged from 2.6 to 2.8 Å, based on the correlation coefficient CC1/2 of 50, and a signal/noise ratio I/σ >1.0 for the last resolution shell. The merged dataset was formatted to MTZ with XDSCONV. The crystal structure was solved by the SAD experimental phasing method with the CRANK-2 [65] package implemented in the CCP4 [66] suite by simultaneously combining initial estimated phases with density modification and model-building in real space. 5 substructure positions were found, from which CRANK-2 automated model building generated a 284 residues partial model that, together with an I-TASSER [67] shredded model for the ATP-cone, was loaded into AUTOBUILD from the PHENIX suite [68], which traced 497 residues. It followed further positional refinement with PHENIX Refine, alternated with manual model building and real space refinement with COOT [69]. An anomalous density map showed discrete peaks higher than 3.0 σ corresponding to zinc at each zinc-finger, and sulfur atoms from five methionine residues (Met48, 70, 86, 107 and 113) from the ATP-cone domain, enabling sequence assignment. Manual model building was alternated with AUTOBUILD followed by PHENIX Refine, imposing non-crystallography symmetry restraints. In addition, analysis of the electron density in molecules A, B, C and D showed clear positive difference densities at the inner cavity of the cone domain that suggested the presence of nucleotides at the “inner” site (see Introduction), which were tentatively placed, refined and visually inspected. The presence of an adenosine was confirmed, yet the number of phosphates clearly varied among subunits. However, tracing got stacked probably due to inconsistencies between datasets; therefore, the structure was refined only against the best data set, which, surprisingly, did not contain adenine at the cone domain in molecule A. The stereochemistry was analyzed, showing no Ramachandran outliers (see crystal quality parameters in Supplementary Table S4). X-ray diffraction data and the crystal structure were deposited at the Protein Data Bank with code 9GEX.

### 4.11. Other protein techniques

Proteins were routinely examined in pre-cast SDS-PAGE gels (4–20% Mini-PROTEAN TGX Precast Protein Gels, Bio-Rad) and stained with a Coomasie-blue-based stain (PageBlue Protein Staining Solution, Thermo Scientific, ThermoFisher), according to the manufacturer’s instructions.

Anti-NrdR western blotting was carried out. A TransBlot-Turbo device and TransBlot-Turbo Mini PVDF Transfer packs (Bio-Rad) were used to transfer the proteins to the membranes, according to the manufacturer’s instructions. As the primary antibody, we used a rabbit polyclonal anti-NrdR serum (ThermoFisher), applying 2 hours of incubation at 4 °C with a 1:500 serum dilution. The detection of primary antibodies was carried out using a goat anti-rabbit horseradish peroxidase (HRP) conjugate (Bio-Rad), applying 1 hour of incubation at 4 °C with a 1:5000 dilution of the antibody. The antibody-antigen complex was detected using the Amersham ECL Primer Western Blotting Reagent (GE Healthcare) according to the manufacturer’s instructions. Proteins were visualized in an ImageQuant LAS4000 Mini device (GE Healthcare).

### 4.12. Atomic force microscopy

NrdR_2_^ECO^-nucleotide (cofactors AMP, ATP, dATP) complexes were prepared as follows. Concentrated NrdR_2_^ECO^ (13.6 mg/ml, approx. 0.8 mM) in 5x NrdR buffer (100 mM Tris-HCl –pH 9.0 at 25 °C–, 400 mM KCl, 5 mM MgCl2, 5 mM DTT, 250 µM ZnSO_4_, 25% glycerol) was mixed in with 10 mM nucleotides prepared in 5x AFM buffer (50 mM HEPES pH 8.2, 250 mM NaCl, 25 mM MgCl_2_, 5 mM TCEP) to a final molar ratio of 20:1 nucleotide:protein and incubated for 1 hour at 25 °C, protected from light (protein concentration = 5.3 mg/ml, approx.. 0.3 mM). Protein-nucleotide complexes were diluted 1:50 in 1x AFM buffer containing 0.2 mM of fresh cofactor (protein concentration = 0.1 mg/ml, approx. 6 µM) and further incubated for 1 hour at 25 °C, protected from light (protein concentration 0.53 mg/ml, approx. 30 µM). After the second incubation, the protein-cofactor mixtures were further diluted in 1x AFM buffer containing no additional cofactor to a final concentration of approx. 1 µM protein and transferred to ice.

Immediately prior to imaging, protein-cofactor complexes were further diluted 1:10 in 1x AFM buffer (protein concentration 0.053 mg/ml, approx. 0.1 µM). 10 µL of the protein-cofactor complexes were deposited on 1-cm diameter freshly exfoliated mica discs. After 2 minutes of incubation, samples were washed with 1 ml of ultrapure, nuclease-free water and gently dried under N_2_ flow. AFM measurements were performed using a Cypher S AFM (Oxford Instruments). Topographic images were obtained in tapping mode using SSS-FMR cantilevers (Nanoandmore) with nominal k = 2.8 N/m, resonance f = 75 kHz, and tip curvature radius < 5 nm. Imaging was performed under N_2_ atmosphere using low oscillation amplitudes (< 300 mV) at the highest amplitude setpoint compatible with image quality to guarantee gentle tip-sample interaction. Under these conditions, the tip-sample interaction regime was attractive, as confirmed by the value of the phase images, which was always higher than 90°. Over 70 images of size 512x512 and 1024x1024 pixels were collected.

For quantification, 5 images per condition were chosen randomly and analyzed in Fiji – ImageJ [70]; three representative images are shown in Figure 2A with +20% brightness and +40% contrast; all quantification used the original, unedited images. Objects were identified using the built-in function *analyze particles* with fixed colour thresholds, a minimal area of 10 nm^2^ and a minimum circularity index of 0.3. Major ellipsoid axis length, minor ellipsoid axis length, area, and circularity of objects were determined, and cumulative distributions of major axis length and circularity index were computed using PRISM 9 (GraphPad Software).

### 4.13. Reverse In Vitro Transcription Assay (ReViTA)

ReViTA is a technique devised to assess the effect of transcription regulators and their cofactors by coupling *in vitro* transcription and qRT-PCR. The pReViTA plasmid contains an arbitrary TEST sequence placed between strong synthetic transcription terminators (see “DNA manipulation and plasmid construction” above) to be transcribed under the control of any promoter cloned into its multi-cloning site. All ReViTA experiments included in this work were performed using pReViTA-PD, a derivative of pReViTA containing the promoter of the *nrdDG* operon from *E. coli* K-12 *substr*. MG1655. A single *in vitro* transcription buffer (IVT buffer) was used for all reactions, with the following composition at a 1x concentration: 40 mM TrisHCl (pH 8.5 at 25 °C), 60 mM KCl, 4 mM MgCl_2_, 1 mM DTT, 10 µM ZnSO_4_, 50 ng/µl bovine serum albumin (BSA), 5% glycerol. DTT and ZnSO_4_ are necessary to keep NrdR functional; BSA and glycerol were included to preserve the protein and facilitate DNA-protein interaction. A ReViTA experiment is comprised of five steps, as previously described [35]:

*Step 1: protein-nucleotide incubation*. The required amount of NrdR was mixed with the nucleotide cofactors at a 20:1 nucleotide protein ratio. Nucleotides were added directly to the concentrated protein as 10 mM solutions dissolved in 1x IVT buffer, and the mixtures were incubated for 1 hour at room temperature.

*Step 2: DNA-protein binding*. Protein-nucleotide complexes were added to 15 µl reactions containing 25 fmol of pReViTA-PD, IVT buffer (at a final 1x concentration), and 5 mM of the same nucleotide used to form the complexes. Reactions with a protein:DNA ratio of 200 or higher also included 1 µg of poly(d[I-C]) (Roche LifeScience) as competition DNA. Reactions were incubated for 1 hour at room temperature.

*Step 3: in vitro transcription* (IVT). To the previous DNA-protein binding reactions, we added the following components in a total of 4.5 µl: NTPs (ATP, CTP, GTP, UTP) to a final concentration of 5 mM each, spermidine to a final concentration of 1 mM, 20 U (0.5 µl) of Ribolock RNase Inhibitor (ThermoFisher Scientific), 0.05 U of yeast inorganic pyrophosphatase (IPPase) (Millipore Sigma), as well as additional nucleotide cofactor and IVT buffer to keep the concentrations at 5 mM and 1x, respectively. Spermidine and IPPase were added to increase the IVT yield. Before starting the IVT reactions, the previous mixes were incubated for 1 hour at room temperature to allow any DNA-protein complexes that had been broken to be re-formed. Then, 0.5 U (0.5 µl) of *E. coli* RNA polymerase saturated with Sigma-70 factor (New England Biolabs) were added to each tube (bringing the volume to 20 µl). Reactions were incubated for 30 min at 37 °C, after which they were stopped by removing the DNA template using TURBO DNase (Invitrogen), according to the manufacturer’s instructions.

*Step 4: RNA quantification by two-step qRT-PCR*. RNA was reverse transcribed using Maxima Reverse Transcriptase (ThermoFisher Scientific), according to the manufacturer’s instructions, and using two gene-specific primers for the TEST and CTRL sequences (primers 21 and 24, respectively). The resulting cDNAs were diluted 1:50 in water and quantified in a qPCR reaction using PowerUP SYBR Green Master Mix (Applied Biosystems), according to the manufacturer’s instructions for absolute quantitation (standard curve). Two independent qPCR reactions were conducted for each cDNA to quantify TEST (primers 19 and 20) and CTRL sequences (primers 22 and 23). Standard curves for both sequences were made by obtaining the purified TEST and CTRL amplicons as purified PCRs as described above, diluting them to a 1 ng/µl concentration, preparing decimal serial dilutions up to 10^-7^, and using 10^-4^, 10^-6^, 10^-7^, and 10^-9^ as standard curve points.

*Step 5: data treatment*. All qRT-PCR data were converted into absolute copy numbers using the standard curves. Then, TEST counts were normalized for each sample by dividing by its corresponding CTRL values to capture all unspecific inhibition. Normalized copies of problem reactions (with NrdR) were then divided by the normalized copies of control reactions (without NrdR) to obtain the relative percentage of transcription activity.

### 4.14. *Galleria mellonella* model of infection

Virulence of wild-type *P. aeruginosa* PAO1, its isogenic mutant PAO1 Δ*nrdR* with *nrdR* expression fully interrupted by a transposon insertion [51], and the complementation of the previous mutation with pUCP20T::*nrdR* [20] were tested on *Galleria mellonella*. *G. mellonella* larvae were routinely grown at 35 °C and 100% humidity. 3 weeks-old larvae were separated for infection and placed in Petri dishes (5 larvae per dish). To prepare bacterial cultures for infection, overnight cultures (in LB medium, with the required antibiotics) of the desired strains were first prepared. Cells were pelleted by centrifugation (5000 g, 10 minutes) and resuspended in sterile PBS, calculated for an OD_550_ of 1.0. The centrifugation-resuspension step was repeated three additional times. Sterile PBS was added to the final suspension to obtain an OD_550_ of 1.0. Decimal serial dilutions of these suspensions in PBS were prepared from 10^-1^ to 10^-5^, and then a 1:5 dilution to 2·10^-6^. 10 µl of the 2·10^-6^ dilutions per larva were injected using Hamilton syringes (Hamilton).

10 larvae (2 Petri dishes) per condition were infected, and a control group of 10 larvae injected with sterile PBS was included. Larvae were then incubated at 37 °C and monitored from 12 h to 24 h after injection every 30 minutes, and a final time 36 h after infection. Kaplan-Meier survival curves were drawn and analyzed using PRISM 9.0 (GraphPad Software).

## Data availability statement

The NrdR ^ECO^ crystal structure data can be found on Protein Data Bank under PDB ID 9GEX. All other data are available in the main text or the supplementary materials.

## CRediT authorship contribution statement

Conceptualization: LP, AS, MS, ET Methodology: LP, AS, CS, MS, ET Investigation: LP, AS, CS, ARC, AnS, AC Supervision: GG, MS, ET Writing—original draft: LP, AS Writing—review and editing: LP, AS, CS, AC, MS, ET

## Acknowledgements

We thank Neus Gual and Carmen Mesas for their help with protein production and DNA binding experiments. We thank the personnel at synchrotrons DESY (Hamburg, Germany) and ALBA (Cerdanyola del Vallès, Spain), and at the Automated Crystallography Platform (IBMB-CSIC) for their highly valuable support.

