## Supplementary Material (figures and tables) for "Structural basis and mechanism of action of NrdR, a bacterial master regulator of ribonucleotide reduction"

Supplementary Figure S1. Expression and purification of NrdR fusion proteins. 2
Supplementary Figure S2. Ion-paired reverse phase HPLC quantification of nucleotides bound to NrdR. 3
Supplementary Figure S3. Size-exclusion chromatography of NrdR-nucleotide complexes. 4
Supplementary Figure S4. SEC-MALS of $NrdR_2^{PAO}$-nucleotide complexes. 5
Supplementary Figure S5. Fine details of the NrdR monomer structure. 5
Supplementary Figure S6. Alignment of 17 NrdR orthologs across the Bacteria domain. 6
Supplementary Figure S7. Full images of EMSA gels. 7
Supplementary Figure S8. Effects of alterations in *nrdR* expression on bacterial virulence and fitness. 8
Supplementary Figure S9. Comparative analysis of *ΔnrdR* transcriptomics data. 9
Supplementary Figure S10. Bioinformatic prediction of NrdR-boxes, random DNA negative control 10
Supplementary Table S1. Transcriptomic effects of *nrdR* inactivation, DNA microarray assay in *E. coli*. 11
Supplementary Table S2. Bioinformatic prediction of NrdR-boxes in *E. coli* and *P. aeruginosa*. 15
Supplementary Table S3. Transcriptomic effects of *nrdR* inactivation, RNA-seq study in *P. aeruginosa*. 17
Supplementary Table S4. X-ray diffraction data collection parameters (…) 20
Supplementary Table S5. Absolute molecular weight of SUMO-$NrdR_2^{ECO}$ (…) 21
Supplementary Table S6. Strains and plasmids in this study. 22
Supplementary Table S7. Sequence and application of the primers used in this study 24

**Supplementary Figure S1. Expression and purification of NrdR fusion proteins.**

**A**: Schematic of the different NrdR fusion proteins used in this study, from N-terminus (left) to C-terminus (right). The following abbreviations are used: His6/$H_6$ (6x histidine tag), SUMO (Small Ubiquitin-like Modifier), AviTag (Protein Biotinylation Tag, not used in this study), TEVcs (Tobacco Etch Virus Cleavage Site), PAO (*Pseudomonas aeruginosa*), ECO (*Escherichia coli*). A red vertical line indicates the exact cleavage site after digestion with SUMO protease ($NrdR_1$) or TEV protease ($NrdR_2$). The unlabeled grey box to the right of the cleavage site represents a linker peptide. Protein schematics are not to scale. **B**: Detailed production of $NrdR_1^{PAO}$. The different steps are illustrated on the schematic (left); samples of each step were analyzed on Coomasie Blue-stained SDS-PAGE gels and Anti-NrdR western blots (right). **C**: highlights of the production of all other fusion proteins. Numbers in the weight marker (WM) represent kDa.

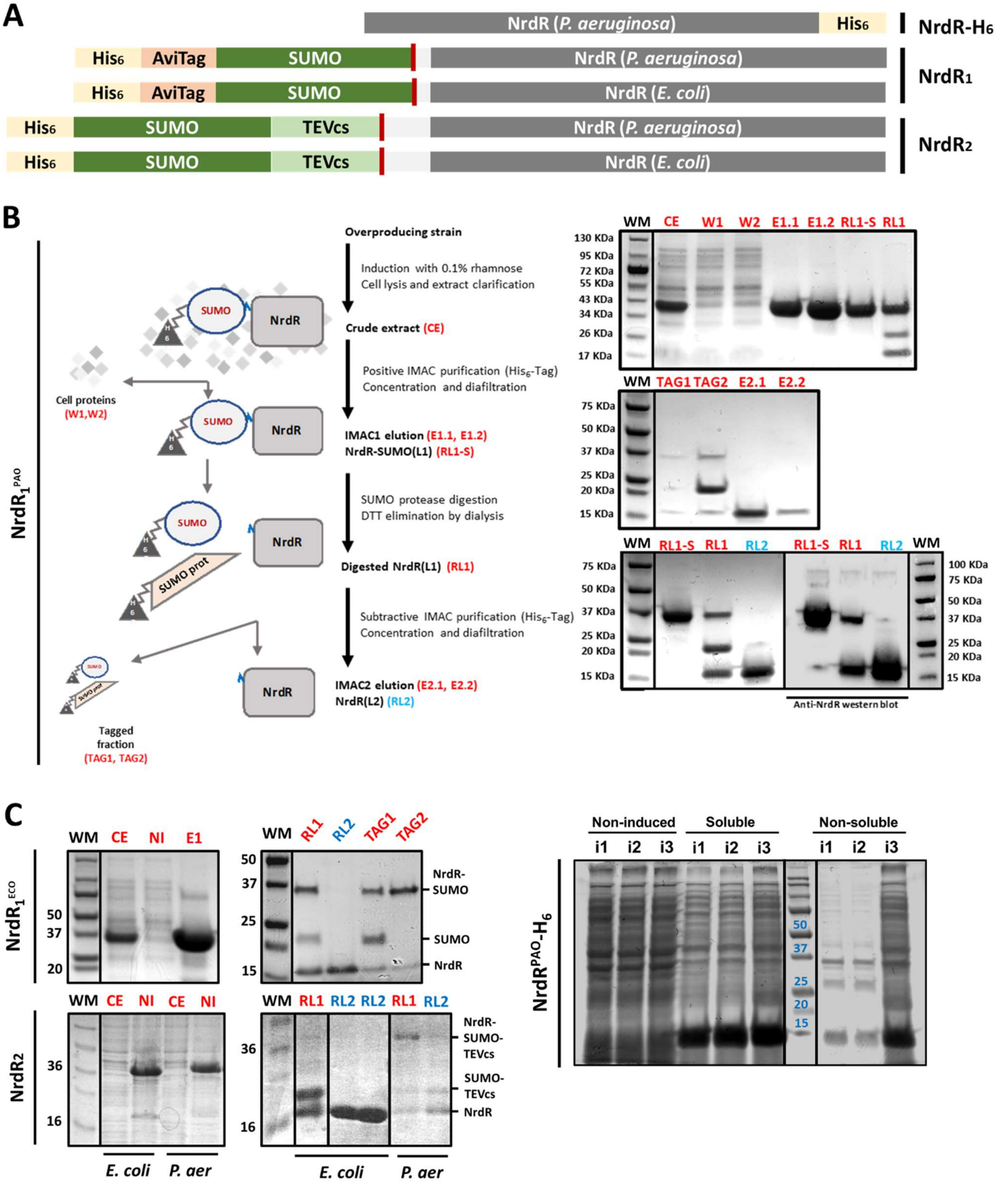

**Supplementary Figure S2. Ion-paired reverse phase HPLC quantification of nucleotides bound to NrdR.**

**A**: Nucleotide quantification in the supernatant of PCA-precipitated protein samples, expressed as percentages relative to moles of protein. As-prepared NrdR$^{PAO}$-$H_6$ (*P. aeruginosa,* top) and $NrdR_2^{ECO}$ (*E. coli*, bottom) are represented in the first column. The next two columns correspond to proteins pre-incubated with ATP and dATP respectively, treated with size-exclusion desalting columns to remove non-bound nucleotides, and precipitated with PCA. dADP was not included in this analysis. **B-I**: Ion-paired reverse phase HPLC chromatograms. (B) is a control experiment without protein in which 20 µmol ATP were eliminated from the sample applying the desalting step from the protocol used for protein samples. The pie chart details the effects of the procedure: 97.57% of nucleotide was successfully eliminated. (C) is a control experiment without protein in which 20 µmol of ATP and 20 µmol of dATP were subject to the PCA precipitation protocol to evaluate the stability of the nucleotides. Pie chart detail the effects for ATP: 87.62% of ATP was recovered as ATP, 8.77% as ADP and 1.07% as AMP; 2.54% was not recovered. (D, E, F) correspond to the experiments using NrdR$^{PAO}$-$H_6$. (G, H, I) correspond to the experiments using $NrdR_2^{ECO}$.

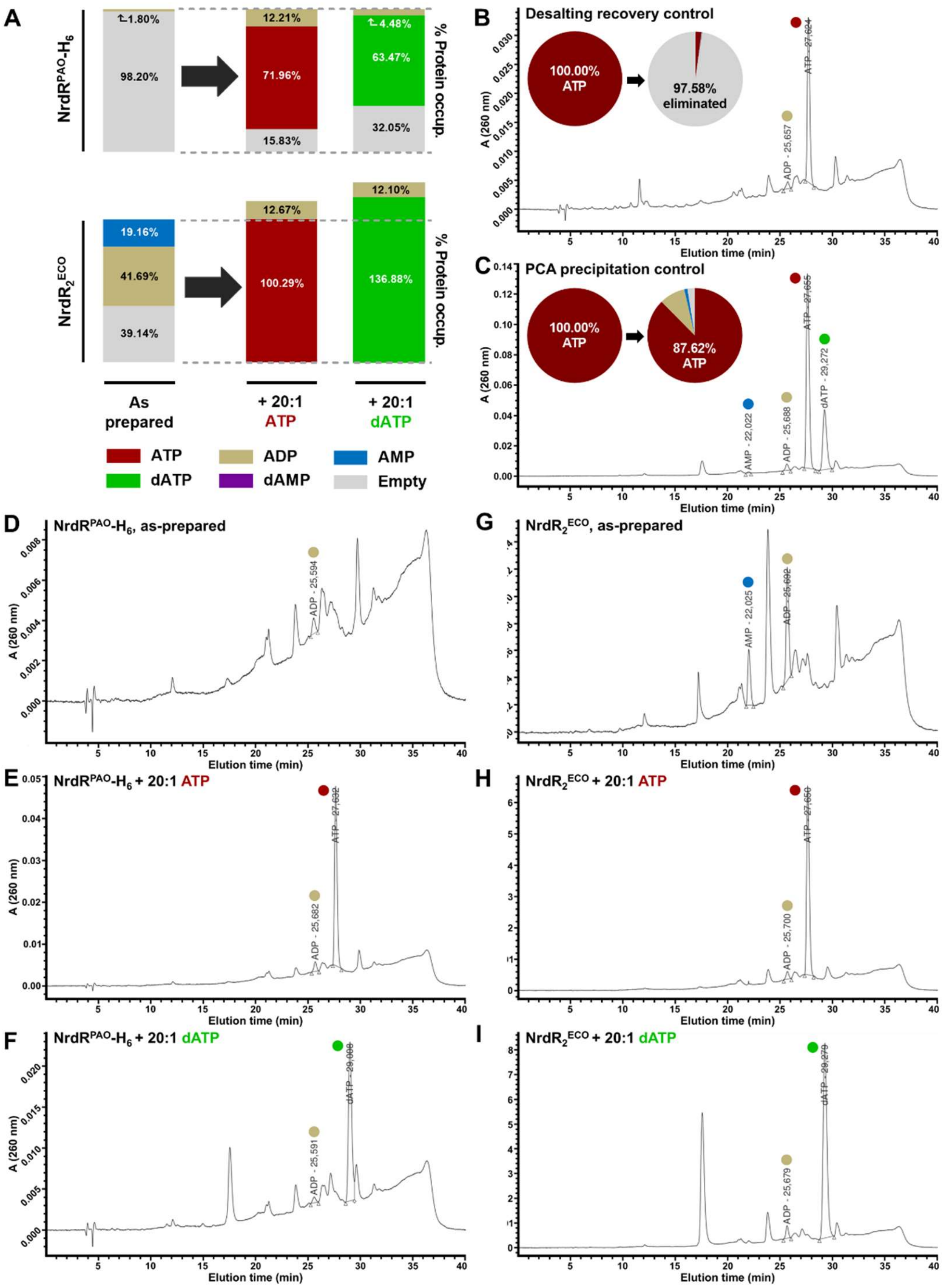

**Supplementary Figure S3. Size-exclusion chromatography of NrdR-nucleotide complexes.**

**A**: SEC chromatogram of NrdR$_1^{ECO}$ protein (from *E. coli*), NrdR$_1^{PAO}$ (from *P. aeruginosa*), and NrdR$^{PAO}$-H$_6$ (from *P. aeruginosa*), pre-incubated with nucleotides at a 20:1 nucleotide:protein ratio and run with the corresponding nucleotide in the elution buffer. Y-axis represents UV absorbance (280 nm) and X-axis represents elution time (min). Equilibration and washing steps between nucleotide series have been removed from the chromatogram for clarity. Protein-containing peaks are labelled indicating their injection and peak numbers (e.g., 1.1 for the first peak of the first injection). Other peaks are caused by the free nucleotide in the samples (labeled as N) or the oxidized DTT in the samples (labeled as D). **B**: Independent chromatograms of NrdR$_1^{PAO}$ bound to ATP, dATP and AMP to compare their elution times. Numbers near the peaks indicate the average estimated molecular weight of the complex based on reference standards (see Materials and Methods). **C**: SDS-PAGE control of representative peaks recovered from the previous SEC experiment. Numbers in the weight marker lane indicate molecular weight in kDa of the band above. **D**: EMSA of the proteins used for SEC experiments before incubation with nucleotides, as an activity control. Numbers below the images indicate protein amounts in pmol. The DNA fragment used is P*nrdA* (*E. coli*). See Materials and Methods for more details on EMSA conditions.

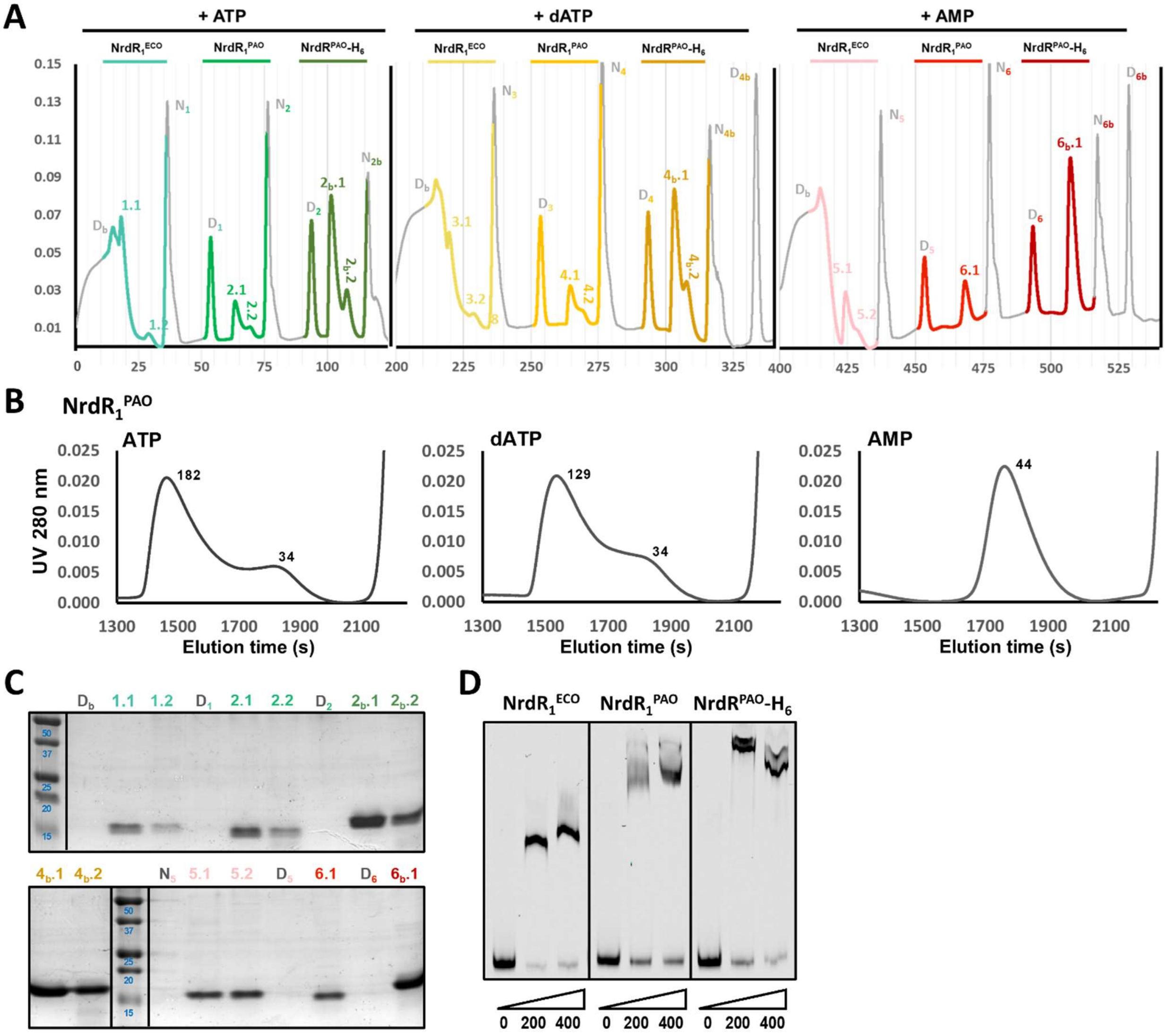

**Supplementary Figure S4. SEC-MALS of $NrdR_2^{PAO}$-nucleotide complexes.**

SEC-MALS results of $NrdR_2^{PAO}$ exposed to 0.025 mM nucleotide in the running buffer. Left Y-axis (solid lines) represents MALS detection data normalized to a maximum signal of 1.0 in each sample. Right Y-axis (dashed lines) represents weight-average molar mass (kDa). Numbers near the peaks indicate the corresponding peak's minimum and maximum molecular weight. Results are representative of two independent experiments.

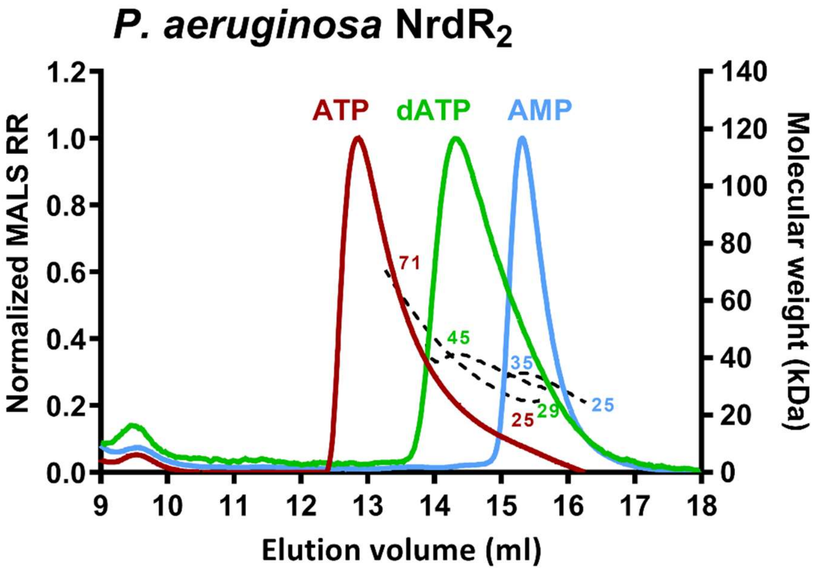

**Supplementary Figure S5. Fine details of the NrdR monomer structure.**

**A**: Overall organization of the NrdR monomer; averaged electron density map. The secondary structure elements are indicated. **B**: Coordination of the $Zn^{2+}$ ion by Cys3 and Cys6 (at the loop preceding strand β1), and Cys31 and Cys34 (Loop Lβ2- β3). **C**: Electron density in molecule A corresponding to an AMP, which could only be solved after averaging three individual datasets.

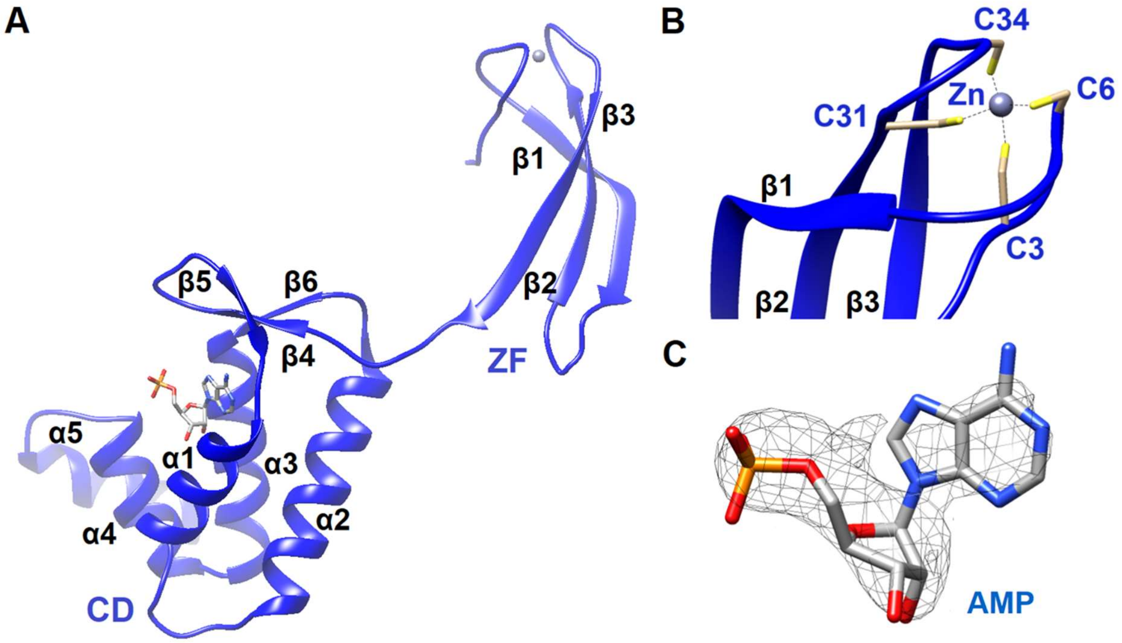

**Supplementary Figure S6. Alignment of 17 NrdR orthologs across the Bacteria domain.**

Multiple sequence alignment performed using Clustal Omega (EMBL-EBI [71]) with default parameters of 17 exemplar NrdR orthologs. To identify key residues responsible for DNA and nucleotide binding, residues have been highlighted depending on the consensus for their position as follows: Orange indicates identity; yellow indicates strong similarity (> 0.5 in the Gonnet PAM 250 matrix); green indicates weak similarity (≤ 0.5 and > 0 in the Gonnet PAM 250 matrix). Numbers above the alignment indicate the position in the *E. coli* sequence, which is used as a reference. The estimated Clustal Omega phylogenetic tree is provided as a guide. The sequences with the following NCBI accession numbers were used for the alignment: *C. acnes (*WP_138224456.1*), C. subtropica (*GAA1996258.1*), S. avermitilis (*WP_403032377.1*), S. spongiae (*WP_152776870.1*), S. coelicolor (*WMT36239.1*), S. clavuligerus (*QPJ95584.1*), P. marinus (*WP_011819663.1*), R. orientalis (*WP_012596313.1*), E. coli (*AAC73516.1*), S. enterica subsp. enterica serovar Typhi (*XGI91390.1*), P. aeruginosa (*AAG07444.1*), P. putida (*WP_392519980.1*), P. fluorescens (*EJZ60269.1*), B. anthracis (*BBK97531.1*), S. pneumoniae (*VDG77748.1*), S. thermophilus (*VUW84032.1*), C. aggregans (*WP_012615593.1*).*

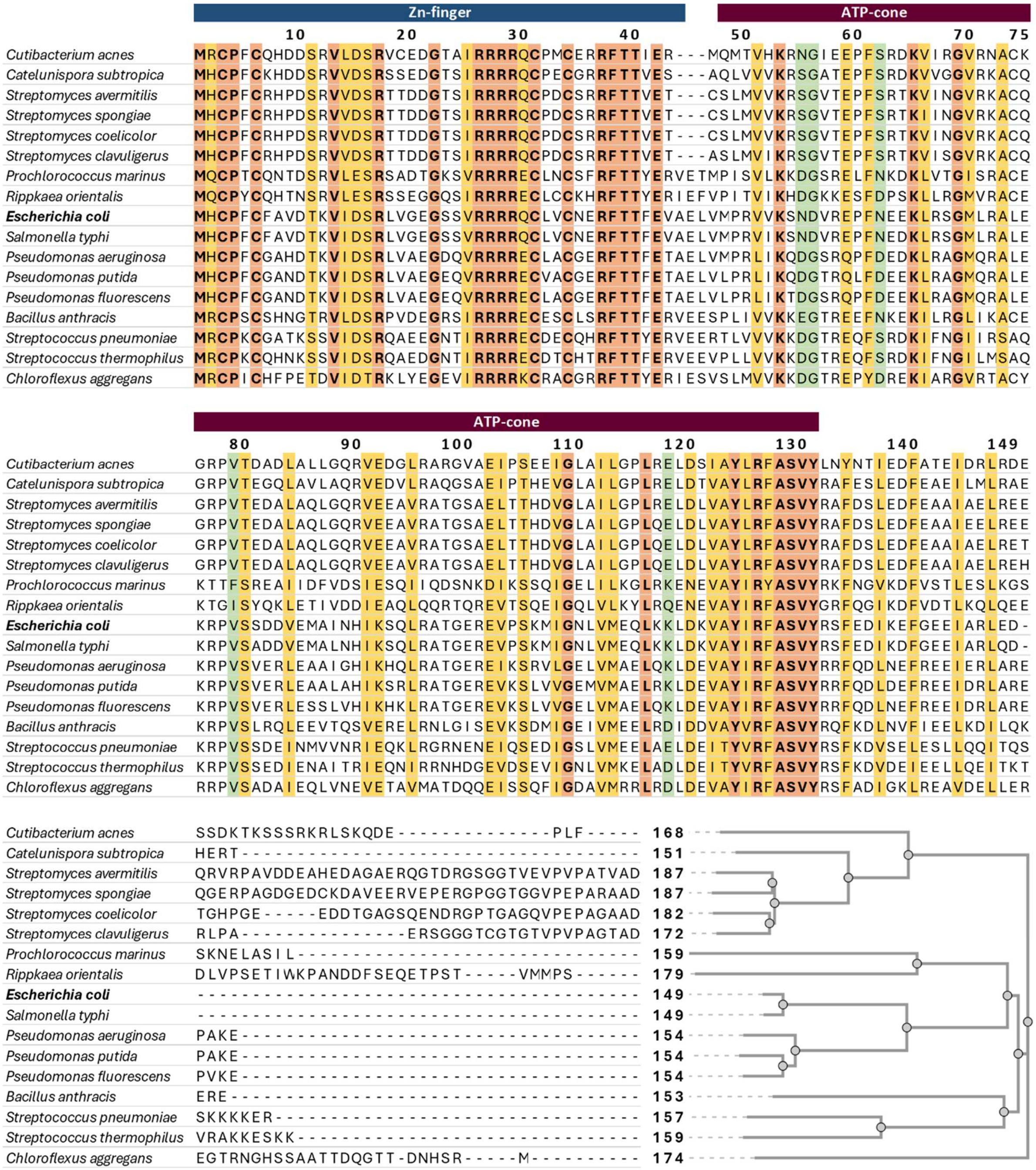

**Supplementary Figure S7. Full images of EMSA gels.**

Full, unedited images of EMSAs conducted using NrdR$_2^{ECO}$ and NrdR$_2^{PAO}$. Two DNA probes were used: P*nrdA*, containing the promoter from the *nrdAB-yfaE* operon in *E. coli* (NrdR sensitive, labeled P*nrdA* (ECO)) and an NrdR-insensitive negative control labeled Ctrl(-). Numbers below the gels indicate the molar ratio of NrdR protein and marked DNA (0:1, 2000:1, and 4000:1 for *E. coli*, 0:1, 2500:1, and 5000:1 for *P. aeruginosa*). Nucleotides indicated below the protein ratios were pre-incubated with NrdR at a fixed 20:1 nucleotide:protein ratio. Two replicate experiments are provided for both protein sources.

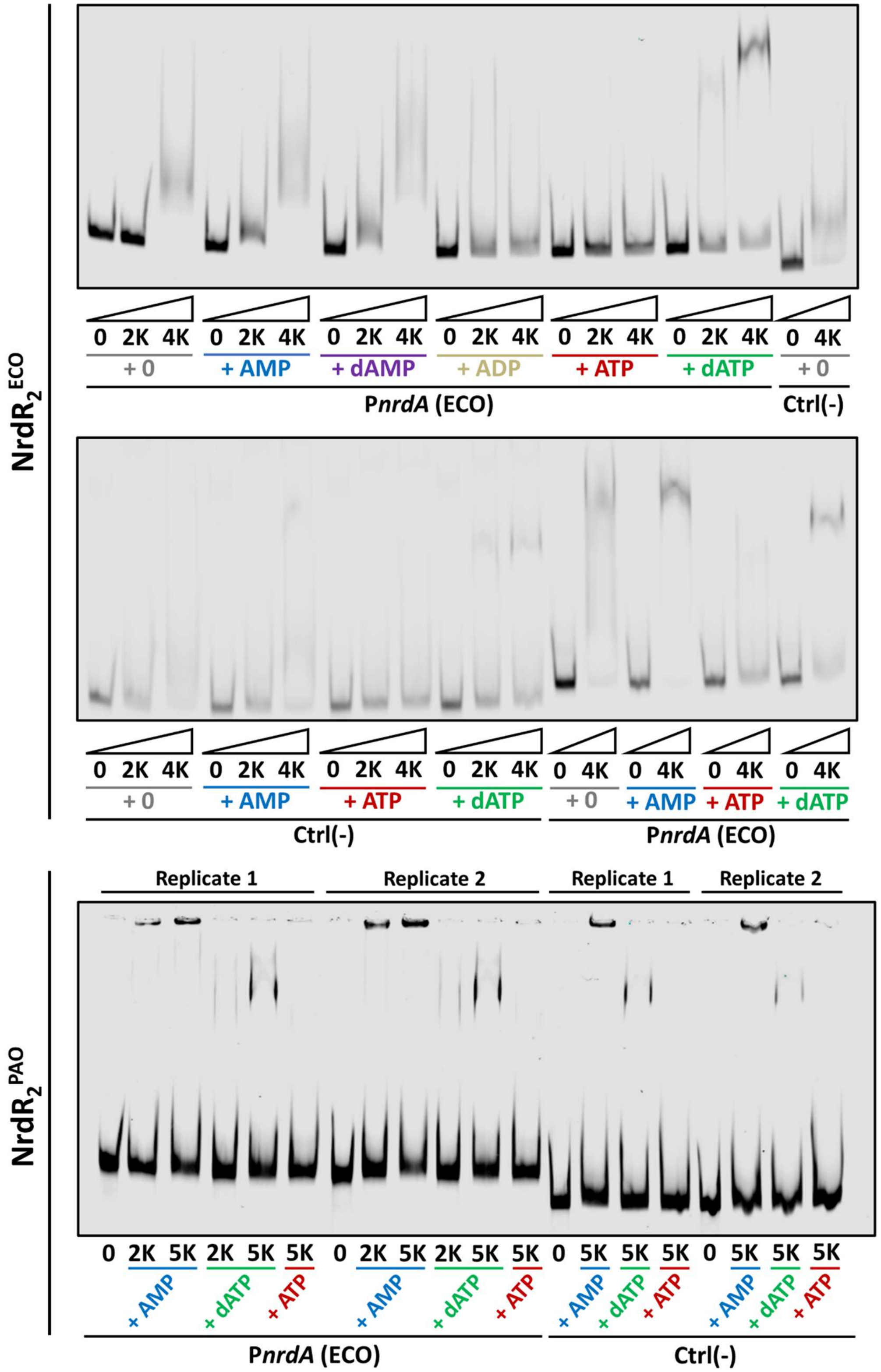

**Supplementary Figure S8. Effects of alterations in *nrdR* expression on bacterial virulence and fitness.**

**A**: growth curve of *P. aeruginosa* PAO1 wild-type (wt), its isogenic *nrdR* mutant strain (Δ*nrdR*), and a complementation strain containing the pUCP20T::*nrdR* plasmid (Δ*nrdR* + $R_c$). $OD_{550}$ values were converted to the equivalent for a 1 cm path length for the reader's convenience. Error bars represent standard deviation. Result is representative of two independent experiments. **B**: Kaplan-Meier survival curve of the previous strains in a *Galleria mellonella* infection assay. Result is representative of two individual experiments.

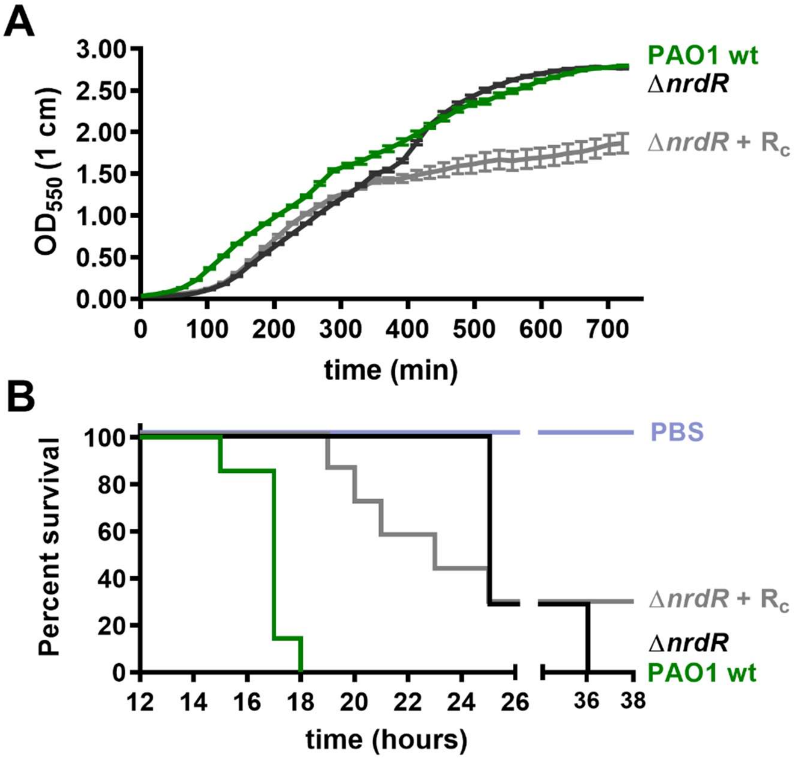

**<u>Supplementary Figure S9. Comparative analysis of *ΔnrdR* transcriptomics data.</u>**

**A**: Comparison of enriched gene ontology (GO) biological processes in the DEGs determined from *nrdR* mutant strains of *E. coli* K-12 *substr*. MG1655 (see Supplementary Table S1) and *P. aeruginosa* PAO1 (see Supplementary Table S3), compared to their corresponding wild-type strains. Lists of enriched GO IDs were produced using PANTHER [58,59], summarized and analyzed with REVIGO [60] and plotted using R Treemap [61]. **B**: Representation of the correlation between DEGs in *P. aeruginosa* and its corresponding orthologs in *E. coli*, as determined by the aforementioned studies. Each square represents a gene in the *P. aeruginosa* genome, from top left (PA0001 *dnaA*) to bottom right (PA5570 *rpmH*). Green-coloured squares represent genes upregulated in PAO Δ*nrdR*, while red-coloured genes were downregulated. Blue-coloured squares represent genes for which at least one ortholog was found to be differentially expressed in *E. coli*. Yellow squares represent genes differentially expressed in both species; operons including these genes are circled in red and detailed to the right. Orthologous genes were determined using OrthologeDB [62].

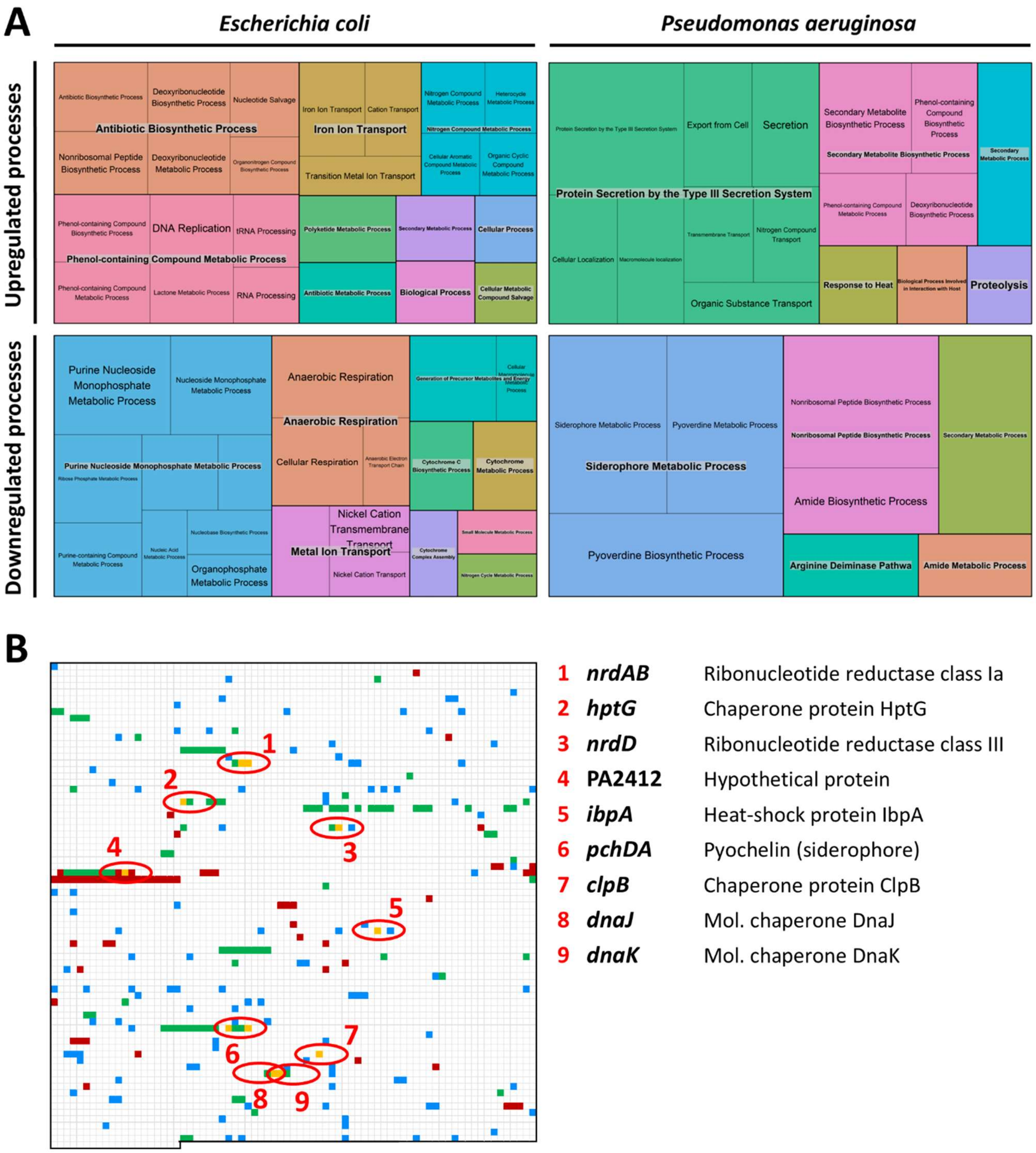

**Supplementary Figure S10. Bioinformatic prediction of NrdR-boxes, random DNA negative control**

Random DNA control to estimate the number of false positives on whole-genome searches for NrdR-boxes in *E. coli* K-12 *substr*. MG1655 (A) and *P. aeruginosa* PAO1 (B). Searches in the real sequences (450 bp upstream of each gene translation start codon; top left) were compared to searches in the same number of sequences, randomly generated with the same length and %GC (Random Sequence Generator, molbiotools.com), repeated three times. Note that the number of hits on random sequences is as high as that of the actual genome or, in the case of *E. coli*, even higher (given the more even distribution of AT/GC pairs), emphasizing the importance of focusing on NrdR-box pairs and correlating their presence with transcriptomics data.

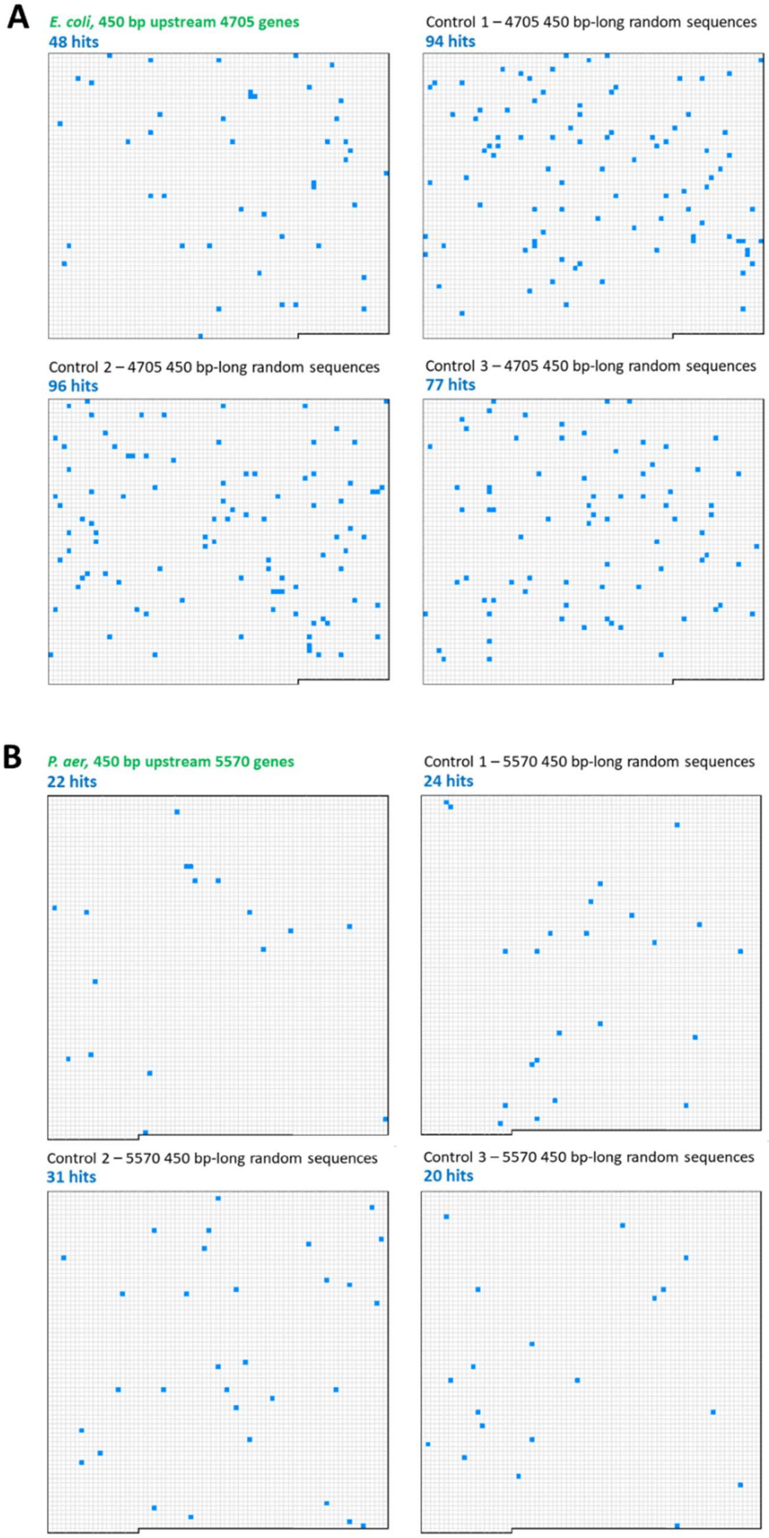

**Supplementary Table S1. Transcriptomic effects of *nrdR* inactivation, DNA microarray assay in *E. coli.***

Differentially expressed genes (DEGs) in a *nrdR*-ATPcone mutant *E. coli* strain compared to its isogenic wild-type K-12 *substr*. MG1655 strain. DEGs were only accepted with a p-value lower than 0.01 and a log(fold-change) higher than 1.0 in at least either exponential or stationary phase. Genes are listed by locus-tag; when available, gene name and a short functional description are also included. Genes in operons displaying putative NrdR-boxes in their upstream region (see Supplementary Table S2) are indicated. Genes in the *nrd* operons are highlighted in bold.

| Upregulated genes | | | | | | | |
|---|---|---|---|---|---|---|---|
| Gene | | | Exponential phase | | Stationary phase | | NrdR-box |
| Locus | Name | Function/Product | LogFC | P-value | LogFC | P-value | Comments |
| b0014 | *dnaK* | Chaperone protein DnaK | 1.31 | 3.63E-06 | | | Single NrdR-box |
| b0015 | *dnaJ* | Chaperone protein DnaJ | 1.39 | 3.10E-06 | | | |
| b0068 | *thiB* | Thiamine ABC transp., periplasmic binding protein | | | 1.07 | 5.22E-03 | |
| b0071 | *leuD* | 3-isopropylmalate dehydratase subunit LeuD | 1.06 | 1.07E-03 | | | |
| b0113 | *pdhR* | DNA-binding transcriptional dual regulator PdhR | | | 1.42 | 1.46E-04 | Single NrdR-box |
| b0150 | *fhuA* | Ferrichrome outer membrane transp. | | | 2.78 | 2.54E-03 | |
| b0151 | *fhuC* | Fe(III) hydroxamate ABC transp., ATP binding subunit | | | 2.09 | 7.93E-03 | |
| b0195 | *trmO* | tRNA m(6)t(6)A37 methyltransferase | | | 1.21 | 7.55E-04 | |
| b0199 | *metN* | L-methionine/D-methionine ABC transp. | | | 1.00 | 8.95E-05 | |
| b0224 | *yafK* | L,D-transpeptidase domain-containing protein | | | 1.08 | 1.07E-04 | |
| b0238 | *gpt* | Xanthine-guanine phsophoribosyltransferase | | | 1.43 | 8.43E-03 | |
| b0314 | *betT* | Choline:H(+) symporter | | | 1.25 | 9.30E-03 | |
| b0315 | *pdeL* | c-di-GMP phosphodiesterase PdeL | | | 1.27 | 2.00E-03 | |
| b0405 | *queA* | TRNA preQ1(34) SAM ribosyltransferase-isomerase | | | 1.37 | 4.24E-03 | |
| b0433 | *ampG* | Muropeptide:H(+) symporter | | | 1.24 | 7.34E-03 | |
| b0468 | *ybaN* | DUF454 domain-containing inner membrane protein | | | 1.75 | 2.48E-03 | |
| b0469 | *apt* | Adenine phosphoribosyltransferase | | | 1.31 | 6.51E-03 | |
| b0473 | *htpG* | Chaperone protein HtpG | 1.11 | 7.52E-05 | | | |
| b0477 | *gsk* | Inosine/guanosine kinase | | | 1.09 | 4.60E-03 | |
| b0524 | *lpxH* | UDP-2,3-diacylglucosamine diphosphatase | | | 1.04 | 8.03E-03 | |
| b0592 | *fepB* | Fe-enterobactin ABC transp., perip. binding protein | | | 2.08 | 2.98E-03 | |
| b0593 | *entC* | Isochorismate synthase EntC | | | 4.26 | 6.92E-03 | |
| b0594 | *entE* | 2,3-dihydroxybenzoate-AMP ligase | | | 3.67 | 8.13E-03 | |
| b0595 | *entB* | Enterobactin synthase component B | | | 2.75 | 8.26E-03 | |
| b0596 | *entA* | 2,3-dihydro-2,3-dihydroxybenzoate dehydrogenase | | | 2.32 | 8.85E-03 | |
| b0597 | *entH* | Proofreading thioesterase, enterobactin biosynthesis | | | 2.34 | 9.91E-03 | |
| b0630 | *lipB* | Lipoyl(octanoyl) transferase | | | 1.26 | 1.87E-03 | |
| b0636 | *rlmH* | 23S rRNA m(3)psi1915 methyltransferase | | | 1.17 | 5.40E-03 | |
| b0737 | *tolQ* | Tol-Pal system protein TolQ | | | 1.40 | 4.11E-03 | |
| b0813 | *rhtA* | L-threonine/L-homoserine exporter | | | 1.01 | 6.81E-03 | |
| b0819 | *ldtB* | L,D-transpeptidase LdtB | | | 1.37 | 3.25E-03 | |
| b0822 | *ybiV* | Sugar phosphatase YbiV | | | 1.25 | 6.59E-04 | |
| b0841 | *ybjG* | Undecaprenyl pyrophosphate phosphatase | | | 1.76 | 3.77E-03 | |
| b0874 | *lysO* | L-lysine exporter | | | 1.39 | 4.72E-03 | |
| b0877 | *ybjX* | DUF535 domain-containing protein YbjX | | | 1.06 | 7.37E-04 | |
| b0931 | *pncB* | Nicotinate phosphoribosyltransferase | | | 1.09 | 2.90E-04 | |
| b0946 | *zapC* | Cell division protein ZapC | 1.05 | 8.72E-03 | | | |
| b1018 | *efeO* | Ferrous iron transport system protein EfeO | | | 2.28 | 1.48E-03 | |
| b1019 | *efeB* | Heme-containing peroxidase/deferrochelatase | | | 1.66 | 4.07E-03 | |
| b1074 | *flgC* | Flagellar basal-body rod protein FlgC | | | 1.12 | 7.84E-03 | |
| b1133 | *mnmA* | TRNA-specific 2-thiouridylase | | | 1.15 | 7.96E-03 | |
| b1203 | *ychF* | Redox-responsive ATPase YchF | | | 1.02 | 4.69E-03 | |
| b1204 | *pth* | Peptidyl-tRNA hydrolase | | | 1.40 | 3.58E-03 | |
| b1216 | *chaA* | Na(+)/K(+):H(+) antiporter ChaA | | | 1.41 | 6.18E-03 | |
| b1238 | *tdk* | Thymidine/deoxyuridine kinase | | | 1.21 | 5.06E-03 | |
| b1253 | *yciA* | Acyl-CoA thioesterase YciA | | | 1.61 | 3.83E-03 | |
| b1295 | *ymjA* | DUF2543 domain-containing protein YmjA | | | 1.30 | 5.47E-03 | |
| b1321 | *ycjX* | DUF463 domain-containing protein YcjX | 1.61 | 2.61E-03 | | | |
| b1322 | *ycjF* | DUF697 domain-containing inner membrane protein | 1.67 | 3.31E-04 | | | |
| b1439 | *ydcR* | Putative DNA-binding transcriptional regulator | | | 1.03 | 7.78E-04 | |
| b1452 | *yncE* | PQQ-like domain-containing protein YncE | | | 1.91 | 8.46E-04 | |
| b1494 | *pqqL* | Periplasmic metalloprotease | | | 1.12 | 4.85E-03 | |
| b1495 | *yddB* | Putative TonB-dependent receptor YddB | | | 1.47 | 5.36E-05 | |
| b1496 | *yddA* | ABC transp. family protein YddA | | | 2.22 | 3.45E-03 | |
| b1524 | *glsB* | Glutaminase 2 | | | 1.01 | 2.64E-03 | |
| b1615 | *uidC* | Outer membrane porin family protein UidC | 1.05 | 2.52E-03 | | | |
| b1626 | *ydgK* | DUF2569 domain-containing inner membrane protein | | | 1.36 | 6.18E-04 | |
| b1627 | *rsxA* | SoxR [2Fe-2S] reducing system protein RsxA | | | 1.70 | 2.48E-03 | |

| | | | | | | | |
|---|---|---|---|---|---|---|---|
| b1628 | *rsxB* | SoxR [2Fe-2S] reducing system protein RsxB | | | 1.55 | 9.96E-03 | |
| b1652 | *rnt* | RNase T | | | 1.21 | 9.02E-03 | |
| b1660 | *ydhC* | Putative transp. YdhC | | | 1.21 | 5.40E-04 | |
| b1704 | *aroH* | 3-deoxy-7-phosphoheptulonate synthase, Trp-sensitive | | | 1.29 | 3.50E-03 | |
| b1759 | *nudG* | 5-hydroxy-CTP diphosphatase | | | 1.02 | 1.93E-03 | |
| b1786 | *dgcJ* | Putative diguanylate cyclase DgcJ | | | 1.44 | 2.21E-03 | |
| b1796 | *yoaG* | DUF1869 domain-containing protein YoaG | 1.30 | 9.53E-03 | 1.59 | 9.21E-04 | |
| b1797 | *yeaR* | DUF1971 domain-containing protein YeaR | | | 1.77 | 7.87E-03 | |
| b1828 | *yebQ* | Putative transp. YebQ | | | 1.38 | 9.76E-03 | |
| b1870 | *cmoA* | Carboxy-S-adenosyl-L-methionine synthase | | | 1.10 | 2.20E-03 | |
| b1888 | *cheA* | Chemotaxis protein CheA | | | 1.08 | 2.49E-03 | |
| b1890 | *motA* | Motility protein A | | | 1.40 | 7.24E-04 | |
| b1905 | *ftnA* | Ferritin iron storage protein | 1.13 | 6.78E-03 | | | |
| b1921 | *fliZ* | DNA-binding transcriptional regulator FliZ | | | 1.57 | 6.28E-03 | |
| b1922 | *fliA* | RNA polymerase, sigma 28 (sigma F) factor | | | 1.60 | 4.91E-03 | |
| b1925 | *fliS* | Flagellar biosynthesis protein FliS | | | 1.03 | 4.17E-03 | |
| b2022 | *hisB* | Imidazoleglycerol-phosphate dehydratase | 1.04 | 1.25E-03 | | | |
| b2066 | *udk* | Uridine/cytidine kinase | | | 1.46 | 5.63E-03 | |
| b2155 | *cirA* | Iron-catecholate outer membrane transp. CirA | | | 3.95 | 7.87E-03 | |
| b2173 | *yeiR* | Zinc-binding GTPase YeiR | | | 1.08 | 4.61E-04 | |
| b2182 | *bcr* | Multidrug efflux pump Bcr | | | 1.40 | 7.28E-03 | |
| b2187 | *yejL* | DUF1414 domain-containing protein YejL | | | 1.00 | 8.81E-03 | |
| b2211 | *yojI* | Microcin J25 efflux protein | | | 3.06 | 7.83E-03 | |
| **b2234** | ***nrdA*** | **RNR class Ia, α subunit** | | | **1.90** | **4.78E-06** | NrdR-box pair |
| **b2235** | ***nrdB*** | **RNR class Ia, β subunit** | | | **1.45** | **4.35E-06** | NrdR-box pair |
| b2236 | *yfaE* | 2Fe-tyrosyl radical cofactor maintenance protein | | | 1.71 | 3.38E-06 | |
| b2273 | *yfbN* | Uncharacterized protein | | | 1.14 | 1.82E-03 | |
| b2350 | *yfdG* | CPS-53 prophage; putative glucose translocase | | | 1.32 | 2.60E-03 | |
| b2392 | *mntH* | Mn(2(+))/Fe(2(+)): H(+) symporter MntH | | | 1.02 | 7.42E-03 | |
| b2393 | *nupC* | Nucleoside:H(+) symporter NupC | | | 1.21 | 2.19E-03 | |
| b2509 | *xseA* | Exodeoxyribonuclease VII subunit XseA | | | 1.37 | 2.06E-04 | |
| b2567 | *rnc* | RNase III | | | 1.23 | 2.37E-03 | |
| b2575 | *yfiC* | TRNA m(6)A37 methyltransferase | | | 1.21 | 1.72E-03 | |
| b2592 | *clpB* | Chaperone protein ClpB | 1.95 | 4.22E-04 | | | |
| b2669 | *stpA* | DNA-binding transcriptional repressor StpA | | | 1.34 | 1.58E-03 | |
| **b2673** | ***nrdH*** | **RNR class Ib, associated glutarredoxin** | **3.19** | **2.36E-08** | **6.52** | **2.12E-09** | NrdR-box pair |
| **b2674** | ***nrdI*** | **RNR class Ib, maintenance flavodoxin** | **3.17** | **1.80E-07** | **6.34** | **6.33E-09** | NrdR-box pair |
| **b2675** | ***nrdE*** | **RNR class Ib, α subunit** | **2.85** | **3.02E-07** | **4.87** | **7.37E-08** | NrdR-box pair |
| **b2676** | ***nrdF*** | **RNR class Ib, β subunit** | **2.67** | **1.94E-05** | **5.98** | **1.95E-07** | NrdR-box pair |
| b2749 | *ygbE* | DUF3561 domain-containing inner membrane protein | | | 1.25 | 6.99E-03 | |
| b2807 | *ygdD* | DUF423 domain-containing inner membrane protein | | | 1.11 | 4.72E-03 | |
| b2817 | *amiC* | N-acetylmuramoyl-L-alanine amidase C | | | 1.01 | 1.08E-04 | |
| b2832 | *ygdQ* | UPF0053 inner membrane protein YgdQ | | | 1.18 | 6.20E-03 | |
| b2838 | *lysA* | Diaminopimelate decarboxylase | | | 1.02 | 3.68E-03 | |
| b2845 | *yqeG* | Putative transp. YqeG | | | 1.10 | 5.33E-03 | |
| b2944 | *yggI* | Protein YggI | | | 1.54 | 1.16E-03 | |
| b3064 | *tsaD* | N(6)-L-threonylcarbamoyladenine synthase subunit | | | 1.00 | 2.70E-03 | |
| b3190 | *ibaG* | Acid stress protein IbaG | | | 1.01 | 4.17E-03 | |
| b3212 | *gltB* | Glutamate synthase subunit GltB | 1.30 | 5.26E-05 | | | |
| b3213 | *gltD* | Glutamate synthase subunit GltD | 1.01 | 2.01E-03 | 1.11 | 8.35E-04 | |
| b3424 | *glpG* | Rhomboid protease GlpG | | | 1.47 | 2.30E-03 | |
| b3425 | *glpE* | Thiosulfate sulfurtransferase GlpE | | | 1.78 | 7.92E-03 | |
| b3541 | *dppD* | Dipeptide ABC transp. ATP binding subunit DppD | 2.05 | 6.69E-03 | | | |
| b3542 | *dppC* | Dipeptide ABC transp. membrane subunit DppC | 1.39 | 3.72E-03 | | | |
| b3543 | *dppB* | Dipeptide ABC transp. membrane subunit DppB | 2.66 | 2.23E-03 | | | |
| b3643 | *rph* | Truncated RNase PH | | | 1.18 | 8.57E-03 | |
| b3661 | *nlpA* | Lipoprotein-28 | 1.24 | 3.44E-03 | | | |
| b3670 | *ilvN* | Acetohydroxy acid synthase I subunit IlvN | 1.79 | 5.89E-05 | | | |
| b3671 | *ilvB* | Acetohydroxy acid synthase I subunit IlvB | 1.78 | 2.90E-04 | | | |
| b3687 | *ibpA* | Small heat shock protein IbpA | | | 1.01 | 9.58E-03 | |
| b3702 | *dnaA* | Chromosomal replication initiator protein DnaA | | | 1.01 | 4.84E-03 | |

| | | | | | | | |
|---|---|---|---|---|---|---|---|
| b3707 | *tnaC* | TnaAB operon leader peptide | 3.65 | 8.96E-04 | | | |
| b3708 | *tnaA* | Tryptophanase | 2.84 | 2.60E-04 | | | |
| b3742 | *mioC* | Flavoprotein MioC | | | 1.09 | 1.06E-03 | |
| b3766 | *ilvL* | IlvXGMEDA operon leader peptide | 1.47 | 3.77E-03 | | | |
| b3778 | *rep* | ATP-dependent DNA helicase Rep | | | 1.17 | 6.44E-03 | |
| b4049 | *dusA* | TRNA-dihydrouridine synthase A | | | 1.11 | 4.63E-03 | |
| **b4238** | ***nrdD*** | **RNR class III, α subunit** | **1.26** | **6.35E-03** | | | NrdR-box pair |
| b4312 | *fimB* | Regulator for *fimA* | 1.08 | 3.28E-03 | | | |
| b4367 | *fhuF* | Hydroxamate siderophore iron reductase | | | 2.67 | 9.19E-03 | |
| b4372 | *holD* | DNA polymerase III subunit | | | 1.06 | 7.48E-03 | |
| b4511 | *ybdZ* | Enterobactin biosynthesis protein YbdZ | | | 3.70 | 5.19E-03 | |

| Downregulated genes | | | | | | | |
|---|---|---|---|---|---|---|---|
| Gene | | | Exponential phase | | Stationary phase | | NrdR-box |
| Locus | Name | Function/Product | LogFC | P-value | LogFC | P-value | Comments |
| b0007 | *yaaJ* | Putative transp. YaaJ | | | -1.22 | 6.31E-03 | |
| b0034 | *caiF* | DNA-binding transcriptional activator CaiF | | | -1.90 | 2.98E-03 | |
| b0119 | *yacL* | UPF0231 family protein YacL | | | -1.02 | 1.45E-03 | |
| b0129 | *yadI* | Putative PTS enzyme IIA component YadI | | | -1.32 | 3.66E-03 | |
| b0336 | *codB* | Cytosine transp. | -1.69 | 1.02E-03 | | | |
| b0337 | *codA* | Cytosine/isoguanine deaminase | -1.23 | 4.68E-06 | | | |
| b0509 | *glxR* | Tartronate semialdehyde reductase 2 | | | -1.18 | 9.60E-03 | |
| b0798 | *ybiA* | N-glycosidase YbiA | | | -1.96 | 2.80E-03 | |
| b0895 | *dmsB* | Dimethyl sulfoxide reductase subunit B | | | -2.04 | 2.31E-03 | |
| b0896 | *dmsC* | Dimethyl sulfoxide reductase subunit C | | | -1.62 | 1.37E-04 | |
| b0953 | *rmf* | Ribosome modulation factor | | | -1.40 | 4.38E-03 | |
| b0994 | *torT* | Periplasmic trimethylamine-N-oxide binding protein | | | -1.20 | 9.57E-03 | |
| b1131 | *purB* | Adenylosuccinate lyase | -1.21 | 6.15E-03 | | | |
| b1182 | *hlyE* | Hemolysin E | | | -1.06 | 8.05E-03 | |
| b1225 | *narH* | Nitrate reductase A subunit beta | | | -3.10 | 7.89E-03 | |
| b1226 | *narJ* | Nitrate reductase 1, Mo-cofactor assembly chaperone | | | -3.35 | 8.73E-03 | |
| b1227 | *narI* | Nitrate reductase A subunit gamma | | | -3.17 | 4.35E-03 | |
| b1420 | *mokB* | Putative regulatory protein MokB | | | -1.55 | 3.05E-03 | |
| b1426 | *ydcH* | Protein YdcH | | | -2.33 | 6.00E-03 | |
| b1462 | *yddH* | Flavin reductase-like protein YddH | | | -1.51 | 7.38E-03 | |
| b1475 | *fdnH* | Formate dehydrogenase N subunit beta | | | -2.53 | 2.43E-03 | |
| b1752 | *ydjZ* | DedA family protein YdjZ | | | -1.82 | 5.07E-03 | |
| b1849 | *purT* | Phosphoribosylglycinamide formyltransferase 2 | -2.50 | 6.17E-03 | | | |
| b2000 | *flu* | CP4-44 prophage; Ag43 autotransp. | -2.20 | 1.15E-05 | -2.69 | 1.58E-06 | |
| b2001 | *yeeR* | CP4-44 prophage; inner membrane protein | -2.69 | 2.86E-07 | -2.45 | 2.66E-07 | |
| b2002 | *yeeS* | CP4-44 prophage; JAB domain-containing protein | -1.94 | 7.38E-05 | -1.39 | 3.87E-05 | |
| b2003 | *yeeT* | CP4-44 prophage; DUF987 domain-containing protein | -1.01 | 1.02E-04 | | | |
| b2009 | *sbmC* | DNA gyrase inhibitor | | | -1.51 | 6.94E-03 | |
| b2197 | *ccmE* | Periplasmic heme chaperone | | | -1.33 | 1.30E-03 | |
| b2198 | *ccmD* | Cytochrome C maturation protein D | | | -1.30 | 1.26E-03 | |
| b2199 | *ccmC* | Cytochrome C maturation protein C | | | -1.23 | 6.38E-03 | |
| b2201 | *ccmA* | Cytochrome C maturation protein A | | | -1.40 | 9.78E-03 | |
| b2202 | *napC* | Periplasmic nitrate reductase cytochrome C protein | | | -1.65 | 7.79E-03 | |
| b2203 | *napB* | Periplasmic nitrate reductase cytochrome C550 protein | | | -1.88 | 2.91E-03 | |
| b2204 | *napH* | Ferredoxin-type protein NapH | | | -1.94 | 2.68E-03 | |
| b2205 | *napG* | Ferredoxin-type protein NapG | | | -2.31 | 4.14E-03 | |
| b2292 | *yfbS* | Putative transp. YfbS | | | -1.88 | 9.29E-03 | |
| b2312 | *purF* | Amidophosphoribosyltransferase | -2.33 | 5.29E-03 | | | |
| b2398 | *yfeC* | Putative DNA-binding transcriptional regulator YfeC | | | -1.67 | 5.24E-03 | |
| b2399 | *yfeD* | Putative DNA-binding transcriptional regulator YfeD | | | -1.45 | 2.06E-03 | |
| b2419 | *yfeK* | DUF5329 domain-containing protein YfeK | -1.87 | 9.32E-04 | | | |
| b2476 | *purC* | Pi-ribosylaminoimidazole-succinocarboxamide synth. | -1.89 | 8.48E-04 | | | |
| b2500 | *purN* | Phosphoribosylglycinamide formyltransferase 1 | -2.14 | 5.71E-03 | | | |

| | | | | | | |
|---|---|---|---|---|---|---|
| b2507 | *guaA* | GMP synthetase | -1.62 | 5.09E-04 | | |
| b2508 | *guaB* | Inosine 5'-monophosphate dehydrogenase | -1.80 | 6.96E-03 | | |
| b2557 | *purL* | Phosphoribosylformylglycinamide synthetase | -2.40 | 1.12E-04 | | |
| b2685 | *emrA* | Multidrug efflux pump membrane fusion protein EmrA | -1.22 | 8.83E-03 | | |
| b2686 | *emrB* | Multidrug efflux pump membrane subunit EmrB | -1.16 | 5.56E-03 | | |
| b2730 | *hypE* | Hydrogenase maturation protein, carbamoyl deH. | | | -1.19 | 1.38E-03 |
| b2768 | *ygcP* | Putative anti-terminator regulatory protein | | | -1.16 | 9.18E-03 |
| b2902 | *ygfF* | Putative oxidoreductase YgfF | -1.26 | 1.39E-04 | | |
| b2971 | *yghG* | Lipoprotein YghG | -1.55 | 5.46E-03 | | |
| b2994 | *hybC* | Hydrogenase 2 large subunit | | | -1.25 | 1.32E-03 |
| b2995 | *hybB* | Hydrogenase 2 membrane subunit | | | -1.70 | 7.53E-04 |
| b2996 | *hybA* | Hydrogenase 2 iron-sulfur protein | | | -2.19 | 8.71E-03 |
| b3055 | *ygiM* | Putative signal transduction protein (SH3 domain) | -1.31 | 3.26E-03 | | |
| b3091 | *uxaA* | D-altronate dehydratase | | | -1.43 | 9.65E-03 |
| b3115 | *tdcD* | Propionate kinase | | | -1.48 | 3.30E-04 |
| b3116 | *tdcC* | Threonine/serine:H(+) symporter | | | -2.04 | 1.02E-03 |
| b3350 | *kefB* | K(+) : H(+) antiporter KefB | | | -1.04 | 2.09E-03 |
| b3478 | *nikC* | Ni(2(+)) ABC transp. membrane subunit NikC | | | -2.26 | 2.44E-04 |
| b3479 | *nikD* | Ni(2(+)) ABC transp. ATP binding subunit NikD | | | -2.28 | 4.51E-04 |
| b3480 | *nikE* | Ni(2(+)) ABC transp. ATP binding subunit NikE | | | -1.65 | 3.42E-05 |
| b3481 | *nikR* | DNA-binding transcriptional repressor NikR | | | -1.07 | 1.60E-04 |
| b3571 | *malS* | Alpha-amylase | | | -2.09 | 1.24E-03 |
| b3573 | *ysaA* | Putative electron transport protein YsaA | | | -2.07 | 8.53E-03 |
| b3612 | *gpmM* | 2,3-BPG-independent phosphoglycerate mutase | | | -1.03 | 9.97E-03 |
| b3645 | *dinD* | DNA damage-inducible protein D | | | -1.37 | 4.50E-05 |
| b3715 | *yieH* | 6-phosphogluconate phosphatase | -1.01 | 7.25E-03 | | |
| b3755 | *yieP* | DNA-binding transcriptional regulator YieP | | | -1.08 | 9.97E-03 |
| b3774 | *ilvC* | Ketol-acid reductoisomerase (NADP(+)) | | | -2.12 | 2.23E-03 |
| b3924 | *fpr* | Flavodoxin/ferredoxin-NADP(+) reductase | | | -1.02 | 9.95E-03 |
| b4005 | *purD* | Phosphoribosylamine--glycine ligase | -2.91 | 1.33E-04 | | |
| b4006 | *purH* | Bifunctional AICAR transformylase/IMP cyclohydrolase | -2.31 | 7.36E-03 | | |
| b4064 | *ghxP* | Guanine/hypoxanthine transp. GhxP | -2.70 | 9.56E-03 | | |
| b4065 | *yjcE* | Putative transp. YjcE | | | -1.05 | 8.91E-03 |
| b4072 | *nrfC* | Putative menaquinol-cytochrome C reductase subunit | | | -2.17 | 4.95E-03 |
| b4073 | *nrfD* | Putative menaquinol-cytochrome C reductase subunit | | | -1.11 | 6.73E-04 |
| b4079 | *fdhF* | Formate dehydrogenase H | | | -1.34 | 6.68E-03 |
| b4217 | *ytfK* | Stringent response modulator YtfK | | | -1.28 | 5.79E-03 |
| b4224 | *chpS* | ChpS antitoxin, ChpB-ChpS toxin-antitoxin system | | | -1.06 | 6.65E-03 |
| b4244 | *pyrI* | Aspartate carbamoyltransferase, PyrI subunit | -1.26 | 2.02E-04 | | |
| b4245 | *pyrB* | Aspartate carbamoyltransferase catalytic subunit | -1.30 | 5.30E-05 | | |
| b4334 | *yjiL* | Activator of (R)-hydroxyglutaryl-CoA dehdratase | | | -1.85 | 4.06E-03 |
| b4428 | *hokB* | Toxin HokB | | | -1.43 | 7.05E-03 |

**Supplementary Table S2. Bioinformatic prediction of NrdR-boxes in *E. coli* and *P. aeruginosa*.**

Putative NrdR-boxes identified in a FIMO (MEME Suite) search on promoter-enriched whole-genome queries containing the sequences 450 bp upstream and 20 bp downstream of the translation start codon for each in gene in the *P. aeruginosa* PAO1 and *E. coli* K-12 *substr*. MG1655 genomes. Only hits with a p-value lower than 5 x $10^{-5}$were included. Results were deduplicated and assigned to the gene with the closest translation start. *nrd* operons are highlighted in bold.

| ***Escherichia coli*** | | | | | | | |
|---|---|---|---|---|---|---|---|
| **Closest gene** | | | **Distance to ATG** | | **Putative NrdR-box** | | |
| **Locus** | **Name** | **Function/ Product** | **Start** | **Stop** | **P-value** | **Sequence** | **Comments** |
| b0014 | *dnaK* | Chaperone protein DnaK | -26 | -11 | 1.98E-05 | ACCGAATATATAGTGG | *dnaKJ* - NrdR box |
| b0055 | *djlA* | Co-chaperone protein DjlA | -55 | -40 | 2.85E-06 | CACCTTTATATTGTGG | |
| b0098 | *secA* | Protein translocation ATPase | -238 | -223 | 4.45E-05 | GCGCAACATCTTGCAT | |
| b0113 | *pdhR* | DNA-binding transcriptional dual regulator PdhR | -312 | -297 | 2.30E-05 | TCTCAATATGTAGAAT | *pdhR-aceEF-lpd* - NrdR box |
| b0213 | *yafS* | Putative SAM-dependent methyltransferase | -286 | -271 | 1.59E-05 | TTCCCTTATCTTGTGT | |
| b0382 | *iraP* | Anti-adaptor protein, sigma stabilization | -173 | -158 | 1.56E-05 | AGCCTATATTTTGTGT | |
| b0460 | *hha* | Hemolysin expression-modulating protein Hha | -155 | -140 | 4.48E-05 | CACCTTTATGTTGTTC | |
| b0583 | *entD* | Phosphopantetheinyl transferase EntD | -88 | -73 | 4.01E-05 | ATTCAATATATTGCAG | |
| b0645 | *ybeR* | Uncharacterized protein YbeR | -257 | -242 | 9.71E-06 | CAACAATATATTGAGC | |
| b0720 | *gltA* | Citrate synthase | -449 | -434 | 2.88E-06 | AACCTACATATAGTTT | |
| b0721 | *sdhC* | Succinate:quinone oxidoreductase | -275 | -260 | 1.57E-05 | AAACTATATGTAGGTT | |
| b0815 | *opgE* | Phosphoethanolamine transferase | -335 | -320 | 2.84E-05 | GCTCTCTATGTTGTGC | |
| b1000 | *cbpA* | Curved DNA-binding protein | -267 | -252 | 9.48E-06 | TACCCATATATAGCGT | |
| b1089 | *rpmF* | 50S ribosomal subunit protein L32 | -320 | -305 | 3.36E-05 | ACACAACGTATTGTTT | |
| b1114 | *mfd* | Transcription-repair coupling factor | -45 | -30 | 1.11E-05 | CCCCCATATGTTGAGG | |
| b1128 | *roxA* | Ribosomal protein-arginine oxygenase | -315 | -300 | 8.90E-06 | CATCTCTATATTGTGG | |
| b1298 | *puuD* | γ-glutamyl-γ-aminobutyrate hydrolase | -335 | -320 | 1.02E-05 | CATCAACATATTGCGT | |
| b1443 | *ydcV* | Putative ABC transporter membrane subunit | -359 | -344 | 3.48E-05 | TCTCTATATCTGGTTG | |
| b1466 | *narW* | Putative private chaperone for NarZ nitrate reductase | -375 | -360 | 3.24E-05 | CGCCAATATGTTGAGT | |
| b1487 | *ddpA* | 2CD-dipep. ABC transporter, periplasmic bind. protein | -340 | -325 | 1.22E-05 | CCGCAATATGTTGTTG | |
| b1491 | *yddW* | Divisome-localized glycosyl hydrolase | -166 | -151 | 2.73E-06 | CCACACTATATTGTGA | |
| b1642 | *slyA* | DNA-binding transcriptional dual regulator SlyA | -249 | -234 | 2.06E-05 | ACACCAGATCTTGTAA | |
| b1642 | *slyA* | DNA-binding transcriptional dual regulator SlyA | -182 | -167 | 3.63E-05 | ACCGAATATATTGCGT | |
| b1791 | *yeaN* | 2-nitroimidazole exporter | -301 | -286 | 3.75E-05 | AAACCGCATATTGTGG | |
| b2025 | *hisF* | Imidazole glycerol phosphate synthase | -398 | -383 | 4.27E-05 | TCACAAGATATGGTGA | |
| b2159 | *nfo* | Endonuclease IV | -72 | -57 | 1.06E-05 | CCACTACATCTTGCTC | |
| **b2234** | ***nrdA*** | **RNR class Ia, α subunit** | **-93** | **-78** | **5.42E-09** | **CCCCTATATATAGTGT** | ***nrdAB*** **- NrdR box 2** |
| **b2234** | ***nrdA*** | **RNR class Ia, α subunit** | **-125** | **-110** | **2.45E-05** | **TCACACTATCTTGCAG** | ***nrdAB*** **- NrdR box 1** |
| b2348 | *argW* | tRNA-Arg | -297 | -282 | 3.45E-05 | ATCCTCTATCTGGTGT | |
| b2351 | *gtrB* | CPS-53 prophage, 3B bactoprenol glucosyl transferase | -158 | -143 | 2.49E-05 | ATGCTATATGTTGGGT | |
| b2543 | *yphA* | Putative inner membrane protein | -27 | -12 | 4.94E-05 | TCACATTATCTTGCAA | |
| b2593 | *yfiH* | Polyphenol oxidase YfiH | -74 | -59 | 1.16E-05 | CCACAAGATATGGTGG | |
| **b2673** | ***nrdH*** | **RNR class Ib, associated glutarredoxin** | **-89** | **-74** | **8.63E-07** | **CAACTACATCTAGTAT** | ***nrdHIEF*** **- NrdR box 2** |
| **b2673** | ***nrdH*** | **RNR class Ib, associated glutarredoxin** | **-120** | **-105** | **2.75E-06** | **TTGCTATATATTGTGT** | ***nrdHIEF*** **- NrdR box 1** |
| b3052 | *hldE* | Heptose 7-P kinase/heptose 1-P adenyltransferase | -314 | -299 | 1.33E-06 | ACCCAATATCTGGTGT | |
| b3155 | *yhbQ* | DNA damage response nuclease YhbQ | -102 | -87 | 3.22E-05 | TGACAACATGTTGTTT | |
| b3180 | *yhbY* | Ribosome assembly factor YhbY | -336 | -321 | 3.56E-05 | TGACCACATATTGTGA | |
| b3186 | *rplU* | 50S ribosomal subunit protein L21 | -304 | -289 | 2.36E-05 | CCGCCATATCTTGCGC | |
| b3210 | *arcB* | Sensor histidine kinase ArcB | -383 | -368 | 4.65E-05 | CCGCTGCATATTGTGA | |
| b3454 | *livF* | Branched chain aa/phenylalanine ABC transporter | -158 | -143 | 3.54E-05 | CACCACTATCTTGTTG | |
| b3647 | *ligB* | DNA ligase B | -285 | -270 | 3.52E-05 | AAACAATATAAAGCGT | |
| b3745 | *viaA* | Putative ATPase cofactor | -242 | -227 | 3.26E-05 | GCCCAACATCTTGTCG | |
| b4177 | *purA* | Adenylosuccinate synthetase | -134 | -119 | 2.28E-05 | CTACTACATGTTGAGG | |
| b4180 | *rlmB* | 23S rRNA 2'-O-ribose G2251 methyltransferase | -76 | -61 | 4.01E-05 | ATTCAATATATTGCAG | |
| **b4238** | ***nrdD*** | **RNR class III** | **-177** | **-162** | **3.18E-07** | **ACCCAATATGTTGTAT** | ***nrdDG*** **- NrdR box 2** |
| **b4238** | ***nrdD*** | **RNR class III** | **-208** | **-193** | **2.99E-05** | **GCACTATATATAGACT** | ***nrdDG*** **- NrdR box 1** |
| b4270 | *leuX* | tRNA-Leu | -292 | -277 | 9.03E-06 | CTTCAACATCTTGTGG | |
| b4684 | *yqfG* | Uncharacterized protein YqfG | -277 | -262 | 3.71E-06 | AACCTCTATATTGTGG | |

| *Pseudomonas aeruginosa* | | | | | | | |
|---|---|---|---|---|---|---|---|
| Closest gene | | | Distance to ATG | | Putative NrdR-box | | |
| Locus | Name | Function/Product | Start | Stop | P-value | Sequence | Comments |
| PA0254 | *hudA* | Virulence attenuating factor HudA | -23 | -8 | 3.11E-05 | CACCTATATGGAGTGG | |
| **PA1156** | ***nrdA*** | **RNR class Ia, α subunit** | **-398** | **-383** | **2.88E-07** | **CCCCTATATCTTGGGT** | ***nrdAB* - NrdR-box 2** |
| **PA1156** | ***nrdA*** | **RNR class Ia, α subunit** | **-194** | **-179** | **1.70E-05** | **CCACTAGGTATTGTGT** | ***nrdDG* - NrdR-box 4** |
| **PA1156** | ***nrdA*** | **RNR class Ia, α subunit** | **-429** | **-414** | **2.70E-05** | **GCGCATTATCTTGTAT** | ***nrdAB* - NrdR-box 1** |
| **PA1157** | | **Probable TCS response regulator** | **-125** | **-110** | **4.86E-08** | **CCACAATATGTAGTGT** | ***nrdAB* - NrdR-box 3** |
| PA1383 | | Hypothetical protein | -181 | -166 | 2.25E-06 | CCACTTCATGTAGTGG | |
| PA1388 | | Hypothetical protein | -32 | -17 | 2.72E-05 | CCACTACGTCTTGTAG | |
| PA1802 | *clpX* | ClpX protease | -42 | -27 | 1.94E-05 | TGCCTTCATCTTGTGT | |
| PA1884 | | Probable transcriptional regulator | -56 | -41 | 4.27E-05 | TTCCAATATATTGGAC | |
| **PA1920** | ***nrdD*** | **RNR class III** | **-57** | **-42** | **1.15E-06** | **CCACAACATATTGTTG** | ***nrdDG* - NrdR-box 1** |
| **PA1920** | ***nrdD*** | **RNR class III** | **-26** | **-11** | **3.37E-06** | **ACACATCATGTTGTGG** | ***nrdDG* - NrdR-box 2** |
| PA2167 | | Hypothetical protein | -250 | -235 | 4.60E-05 | CCCCTCCATCTAGAAG | |
| PA2229 | | Conserved hypothetical protein | -53 | -38 | 3.39E-05 | GGACTCTATATTGTAT | |
| PA2523 | *czcR* | Heavy metal resp., TCS response regulator | -21 | -6 | 2.35E-05 | ATACTTTATATAGGGG | |
| PA3011 | *topA* | DNA topoisomerase I | -76 | -61 | 3.72E-07 | CCACTATATATAGCGG | *topA* - NrdR-box |
| PA4210 | *phzA1* | Phenazine biosynthesis protein PhzA1 | -388 | -373 | 3.86E-05 | CTACCAGATCTTGTAG | |
| PA4280 | *birA* | BirA bifunctional protein | -38 | -23 | 3.54E-05 | CCTCTATATGATGCGT | |
| PA4523 | | Hypothetical protein | -152 | -137 | 3.11E-05 | ACCCTCTATCTAGATT | |
| PA5325 | *sphA* | Sphingosine-dependent virulence factor | -156 | -141 | 1.04E-05 | TCCCATCATATAGCGT | |
| **PA5497** | ***nrdJa*** | **RNR class II, polypeptide A** | **-7** | **8** | **2.25E-05** | **TAACTAGATGTTGCGT** | ***nrdJab* - NrdR-box 2** |
| **PA5497** | ***nrdJa*** | **RNR class II, polypeptide A** | **-38** | **-23** | **2.41E-05** | **ACACAAGATATTGATT** | ***nrdJab* - NrdR-box 1** |

**<u>Supplementary Table S3. Transcriptomic effects of *nrdR* inactivation, RNA-seq study in *P. aeruginosa*.</u>**

DEGs in a Δ*nrdR* mutant strain in *P. aeruginosa* compared to its isogenic wild-type PAO1 strain. DEGs were only accepted with a false discovery rate (FDR) lower than 0.01 and a log(fold-change) higher than 1.0. Low quality mapping hits and poorly expressed genes were filtered out. Genes are listed by locus-tag; when available, gene name and a short functional description are also included. Genes in operons displaying putative NrdR-boxes in their upstream region (see Supplementary Table S2) are indicated. Genes in the *nrd* operons are highlighted in bold.

| Upregulated genes | | | | | |
|---|---|---|---|---|---|
| Gene | | | Δ*nrdR* vs WT | | NrdR-box |
| Locus | Name | Function/Product | LogFC | FDR | Comments |
| PA0201 | | Hypothetical protein | 1.22 | 2.20E-41 | |
| PA0526 | | Hypothetical protein | 1.27 | 8.62E-34 | |
| PA0527 | *dnr* | Transcriptional regulator Dnr | 1.26 | 2.54E-71 | |
| PA0604 | | ABC transporter | 1.20 | 4.56E-85 | |
| PA0605 | | ABC transporter permease | 1.40 | 5.82E-42 | |
| PA0606 | | ABC transporter permease | 1.41 | 5.60E-61 | |
| PA0779 | | ATP-dependent protease | 1.43 | 3.80E-90 | |
| PA0839 | | Transcriptional regulator | 2.60 | 2.79E-223 | |
| PA0996 | *pqsA* | Anthranilate-CoA ligase | 2.54 | 0.00E+00 | |
| PA0997 | *pqsB* | Hypothetical protein | 2.44 | 3.21E-268 | |
| PA0998 | *pqsC* | Hypothetical protein | 2.29 | 2.64E-163 | |
| PA0999 | *pqsD* | 3-oxoacyl-ACP synthase | 1.95 | 1.58E-93 | |
| PA1000 | *pqsE* | Thioesterase PqsE | 1.77 | 8.17E-111 | |
| PA1001 | *phnA* | Anthranilate synthase component I | 1.86 | 1.19E-100 | |
| PA1002 | *phnB* | Anthranilate synthase component II | 1.82 | 2.98E-117 | |
| PA1154 | | Hypothetical protein | 2.31 | 1.40E-53 | |
| **PA1155** | ***nrdB*** | **RNR class Ia, β subunit** | **1.78** | **3.63E-156** | **NrdR-box pair (+2)** |
| **PA1156** | ***nrdA*** | **RNR class Ia, α subunit** | **1.63** | **4.63E-93** | **NrdR-box pair (+2)** |
| PA1331 | | Hypothetical protein | 1.46 | 5.16E-135 | |
| PA1429 | | Cation-transporting P-type ATPase | 1.24 | 2.16E-65 | |
| PA1546 | *hemN* | Oxygen-independent coproporphyrinogen-III oxidase | 1.24 | 1.13E-95 | |
| PA1596 | *htpG* | Chaperone protein HtpG | 1.32 | 3.89E-74 | |
| PA1597 | | Hypothetical protein | 1.97 | 8.81E-184 | |
| PA1600 | | Cytochrome C | 1.92 | 1.01E-108 | |
| PA1601 | | Aldehyde dehydrogenase | 1.83 | 6.93E-202 | |
| PA1602 | | Oxidoreductase | 1.81 | 5.83E-140 | |
| PA1649 | | Short-chain dehydrogenase | 1.54 | 2.87E-67 | |
| PA1690 | *pscU* | Translocation protein in type III secretion | 1.06 | 6.33E-39 | |
| PA1691 | *pscT* | Translocation protein in type III secretion | 1.12 | 1.06E-31 | |
| PA1692 | | Translocation protein in type III secretion | 1.07 | 1.23E-33 | |
| PA1694 | *pscQ* | Type III secretion system protein | 1.27 | 2.11E-71 | |
| PA1695 | *pscP* | Translocation protein in type III secretion | 1.28 | 1.50E-46 | |
| PA1696 | *pscO* | Translocation protein in type III secretion | 1.15 | 1.27E-51 | |
| PA1698 | *popN* | Type III secretion outer membrane protein PopN | 1.03 | 5.26E-50 | |
| PA1700 | | Hypothetical protein | 1.01 | 3.78E-39 | |
| PA1701 | | Hypothetical protein | 1.22 | 2.29E-31 | |
| PA1702 | | Hypothetical protein | 1.26 | 4.50E-28 | |
| PA1703 | *pcrD* | Type III secretory apparatus protein PcrD | 1.08 | 5.05E-79 | |
| PA1707 | *pcrH* | Regulatory protein PcrH | 1.24 | 3.67E-48 | |
| PA1708 | *popB* | Translocator protein PopB | 1.11 | 2.74E-86 | |
| PA1709 | *popD* | Translocator outer membrane protein PopD | 1.03 | 5.57E-70 | |
| PA1715 | *pscB* | Type III export apparatus protein | 1.04 | 6.56E-56 | |
| PA1721 | *pscH* | Type III export protein PscH | 1.06 | 8.49E-45 | |
| PA1722 | *pscI* | Type III export protein PscI | 1.01 | 3.34E-44 | |
| PA1896 | | Hypothetical protein | 1.12 | 1.68E-08 | |
| **PA1919** | ***nrdG*** | **RNR class III activating protein** | **4.89** | **1.05E-155** | **NrdR-box pair** |
| **PA1920** | ***nrdD*** | **RNR class III** | **6.33** | **0.00E+00** | **NrdR-box pair** |

| | | | | | |
|---|---|---|---|---|---|
| PA2018 | | Multidrug efflux protein | 1.37 | 1.01E-126 | |
| PA2019 | | Multidrug efflux lipoprotein | 1.57 | 1.36E-48 | |
| PA2193 | *hcnA* | Hydrogen cyanide synthase subunit HcnA | 1.20 | 1.09E-24 | |
| PA2194 | *hcnB* | Hydrogen cyanide synthase subunit HcnB | 1.11 | 4.54E-33 | |
| PA2322 | | Gluconate permease | 1.42 | 9.49E-60 | |
| PA2403 | | Hypothetical protein | 1.35 | 2.47E-99 | |
| PA2404 | | Hypothetical protein | 1.42 | 1.36E-89 | |
| PA2405 | | Hypothetical protein | 1.41 | 1.09E-80 | |
| PA2406 | | Hypothetical protein | 1.53 | 1.95E-78 | |
| PA2407 | | Adhesion protein | 1.66 | 4.85E-135 | |
| PA2408 | | ABC transporter ATP-binding protein | 1.53 | 6.27E-43 | |
| PA2409 | | ABC transporter permease | 1.41 | 2.08E-84 | |
| PA2410 | | Hypothetical protein | 1.22 | 1.83E-67 | |
| PA2550 | | Acyl-CoA dehydrogenase | 2.77 | 5.73E-261 | |
| PA2593 | *qteE* | Quorum threshold expression protein QteE | 1.23 | 5.65E-18 | |
| PA3126 | *ibpA* | Heat-shock protein IbpA | 1.08 | 9.59E-51 | |
| PA3327 | | Non-ribosomal peptide synthetase | 1.79 | 1.25E-88 | |
| PA3328 | | FAD-dependent monooxygenase | 2.28 | 1.45E-37 | |
| PA3329 | | Hypothetical protein | 1.73 | 1.59E-19 | |
| PA3330 | | Short-chain dehydrogenase | 1.46 | 1.02E-13 | |
| PA3331 | | Cytochrome P450 | 1.66 | 1.67E-26 | |
| PA3332 | | Hypothetical protein | 1.88 | 2.05E-19 | |
| PA3333 | *fabH2* | 3-oxoacyl-ACP synthase III | 1.60 | 1.93E-17 | |
| PA3334 | | Acyl carrier protein | 1.26 | 1.80E-12 | |
| PA3427 | | Short-chain dehydrogenase | 1.17 | 2.47E-58 | |
| PA3449 | | Hypothetical protein | 1.35 | 7.50E-11 | |
| PA3479 | *rhlA* | Rhamnosyltransferase subunit A | 1.03 | 4.83E-14 | |
| PA3661 | | Hypothetical protein | 1.11 | 4.11E-09 | |
| PA3912 | | Hypothetical protein | 1.08 | 9.41E-35 | |
| PA4056 | *ribD* | Riboflavin-specific deaminase/reductase | 1.49 | 4.54E-64 | |
| PA4057 | *nrdR* | Transcriptional regulator NrdR | 1.34 | 6.56E-127 | |
| PA4218 | | Transporter | 2.22 | 2.14E-56 | |
| PA4219 | | Hypothetical protein | 1.63 | 2.85E-60 | |
| PA4220 | | Hypothetical protein | 1.36 | 2.25E-21 | |
| PA4221 | *fptA* | Fe(III)-pyochelin outer membrane receptor | 1.41 | 1.83E-72 | |
| PA4222 | | ABC transporter ATP-binding protein | 1.27 | 1.38E-26 | |
| PA4223 | | ABC transporter ATP-binding protein | 1.05 | 2.03E-32 | |
| PA4224 | *pchG* | Pyochelin biosynthetic protein PchG | 1.74 | 2.11E-47 | |
| PA4225 | *pchF* | Pyochelin synthetase | 1.81 | 2.22E-89 | |
| PA4226 | *pchE* | Dihydroaeruginoic acid synthetase | 1.86 | 1.97E-108 | |
| PA4228 | *pchD* | 2%2C3-dihydroxybenzoate-AMP ligase | 1.61 | 1.60E-39 | |
| PA4229 | *pchC* | Pyochelin biosynthetic protein PchC | 1.63 | 2.94E-22 | |
| PA4230 | *pchB* | Isochorismate-pyruvate lyase | 1.84 | 8.25E-17 | |
| PA4231 | *pchA* | Salicylate biosynthesis isochorismate synthase | 1.58 | 5.14E-40 | |
| PA4542 | *clpB* | Chaperone protein ClpB | 1.12 | 2.39E-89 | |
| PA4759 | *dapB* | 4-hydroxy-tetrahydrodipicolinate reductase | 1.29 | 2.96E-123 | |
| PA4760 | *dnaJ* | Molecular chaperone DnaJ | 1.22 | 1.73E-69 | |
| PA4761 | *dnaK* | Molecular chaperone DnaK | 1.28 | 3.15E-84 | |
| PA4762 | *grpE* | Heat shock protein GrpE | 1.05 | 1.44E-64 | |
| PA5020 | | Acyl-CoA dehydrogenase | 2.01 | 1.51E-169 | |
| PA5054 | *hslU* | ATP-dependent protease ATP-binding subunit HslU | 1.33 | 1.71E-112 | |
| PA5055 | | Hypothetical protein | 1.28 | 3.59E-77 | |
| PA5207 | | Phosphate transporter | 1.12 | 5.07E-52 | |
| **PA5496** | ***nrdJb*** | **RNR class II, polypeptide B** | **3.29** | **0.00E+00** | **NrdR-box pair** |
| **PA5497** | ***nrdJa*** | **RNR class II, polypeptide A** | **3.66** | **0.00E+00** | **NrdR-box pair** |

| Downregulated genes | | | | | |
|---|---|---|---|---|---|
| Gene | | | Δ*nrdR* vs WT | | NrdR-box |
| Locus | Name | Function/Product | LogFC | P-value | Comments |
| PA0132 | | Beta alanine--pyruvate transaminase | -1.28 | 1.46E-54 | |
| PA0887 | *acsA* | Acetyl-CoA synthetase | -1.10 | 5.72E-57 | |
| PA1744 | | Hypothetical protein | -2.19 | 2.44E-12 | |
| PA1942 | | Hypothetical protein | -3.21 | 7.98E-75 | |
| PA1970 | | Hypothetical protein | -1.16 | 2.14E-11 | |
| PA2394 | *pvdN* | Pyoverdine biosynthesis protein PvdN | -1.15 | 9.20E-19 | |
| PA2395 | *pvdO* | Pyoverdine biosynthesis protein PvdO | -1.40 | 7.52E-39 | |
| PA2400 | *pvdJ* | Pyoverdine biosynthesis protein PvdJ | -1.05 | 5.63E-47 | |
| PA2402 | | Peptide synthase | -1.13 | 5.03E-91 | |
| PA2411 | | Thioesterase | -1.10 | 4.98E-38 | |
| PA2412 | | Hypothetical protein | -1.16 | 5.01E-22 | |
| PA2413 | *pvdH* | Diaminobutyrate--2-oxoglutarate aminotransferase | -1.22 | 1.53E-37 | |
| PA2424 | | Peptide synthase | -1.50 | 5.89E-86 | |
| PA2425 | *pvdG* | Pyoverdine biosynthesis protein PvdG | -1.14 | 7.54E-09 | |
| PA2426 | *pvdS* | Extracytoplasmic-function sigma-70 factor | -1.59 | 5.25E-31 | |
| PA2474 | | Hypothetical protein | -7.32 | 3.82E-16 | |
| PA2475 | | Cytochrome P450 | -10.89 | 1.29E-159 | |
| PA2476 | *dsbG* | Thiol:disulfide interchange protein DsbG | -12.30 | 0.00E+00 | |
| PA2477 | | Thiol:disulfide interchange protein | -11.36 | 8.12E-170 | |
| PA2478 | | Thiol:disulfide interchange protein DsbD | -9.25 | 9.89E-282 | |
| PA2479 | | Two-component response regulator | -7.52 | 2.69E-101 | |
| PA2480 | | Two-component sensor | -9.43 | 1.35E-146 | |
| PA2481 | | Hypothetical protein | -11.86 | 0.00E+00 | |
| PA2482 | | Cytochrome C | -11.52 | 0.00E+00 | |
| PA2483 | | Hypothetical protein | -10.81 | 0.00E+00 | |
| PA2484 | | Hypothetical protein | -10.45 | 9.10E-297 | |
| PA2485 | | Hypothetical protein | -10.56 | 1.19E-137 | |
| PA2486 | | Hypothetical protein | -10.18 | 3.95E-102 | |
| PA2487 | | Hypothetical protein | -10.20 | 2.51E-105 | |
| PA2488 | | Transcriptional regulator | -8.54 | 4.36E-116 | |
| PA2489 | | Transcriptional regulator | -10.55 | 5.12E-100 | |
| PA2490 | | Hypothetical protein | -8.40 | 2.87E-29 | |
| PA2491 | *mexS* | Oxidoreductase | -11.09 | 0.00E+00 | |
| PA2492 | *mexT* | Transcriptional regulator MexT | -9.61 | 0.00E+00 | |
| PA2493 | *mexE* | Multidrug efflux membrane fusion protein MexE | -7.95 | 0.00E+00 | |
| PA2494 | *mexF* | Multidrug efflux transporter MexF | -6.51 | 0.00E+00 | |
| PA2495 | *oprN* | Multidrug efflux outer membrane protein OprN | -5.32 | 0.00E+00 | |
| PA2754a | | Hypothetical protein | -1.05 | 4.27E-14 | |
| PA2759 | | Hypothetical protein | -1.23 | 2.71E-20 | |
| PA2811 | | ABC transporter permease | -1.58 | 8.53E-127 | |
| PA2812 | | ABC transporter ATP-binding protein | -1.85 | 5.49E-219 | |
| PA2813 | | Glutathione S-transferase | -3.14 | 0.00E+00 | |
| PA3038 | | Porin | -1.09 | 1.81E-49 | |
| PA3189 | | Sugar ABC transporter permease | -1.06 | 4.68E-53 | |
| PA3229 | | Hypothetical protein | -4.13 | 0.00E+00 | |
| PA3234 | | Acetate permease | -1.25 | 9.17E-77 | |
| PA3235 | | Hypothetical protein | -1.31 | 9.68E-73 | |
| PA3268 | | TonB-dependent receptor | -1.20 | 3.79E-96 | |
| PA3496 | | Hypothetical protein | -1.23 | 2.68E-30 | |
| PA3530 | | Hypothetical protein | -1.30 | 4.77E-66 | |
| PA3568 | | Propionyl-CoA synthetase | -1.12 | 8.31E-37 | |
| PA3901 | *fecA* | Fe(III) dicitrate transporter FecA | -2.51 | 1.53E-299 | |
| PA4354 | | Hypothetical protein | -2.32 | 1.08E-183 | |
| PA4355 | | Major facilitator superfamily transporter | -1.84 | 1.59E-161 | |
| PA4356 | *xenB* | Xenobiotic reductase | -1.72 | 5.29E-134 | |
| PA4514 | | Iron transport outer membrane receptor | -2.10 | 1.15E-214 | |
| PA4623 | | Hypothetical protein | -1.36 | 5.05E-17 | |
| PA4710 | *phuR* | Heme/hemoglobin uptake outer membrane receptor | -1.32 | 1.04E-44 | |
| PA4881 | | Hypothetical protein | -5.03 | 1.00E-161 | |
| PA5171 | *arcA* | Arginine deiminase | -1.71 | 4.53E-140 | |
| PA5172 | *arcB* | Ornithine carbamoyltransferase | -2.50 | 1.29E-231 | |
| PA5173 | *arcC* | Carbamate kinase | -2.46 | 0.00E+00 | |

**Supplementary Table S4. X-ray diffraction data collection parameters; processing, scaling, and refinement statistics.**

Data collection, processing, and scaling parameters (wavelength, resolution range, space group, unit cell parameters, unique reflections, multiplicity, completeness, I/sigma, CCI/2) and refinement statistics (reflections used in refinement and for R-free, R-work and R-free, CC work and CC free, number of non-hydrogen atoms and macromolecules, number of protein residues, RMS bonds and angles, Ramachandran favored, allowed and outliers, clashscore, average B-factor and B-factor macromolecules). Values in brackets are for the last resolution shell.

| | |
|---|---|
| **Wavelength (Å)** | 1.2835 |
| **Resolution range (Å)** | 92.421-2.600 (2.693-2.600) |
| **Space group** | $C222_1$ |
| **Unit cell parameters a, b, c (Å); α, β, γ (°)** | 51.3, 256.3 ,133.4; 90, 90, 90 |
| **Unique reflections** | 27674 (2709) |
| **Multiplicity** | 15.0 (15.2) |
| **Completeness (%)** | 99.91 (99.96) |
| **I/ sigma(I)** | 26.13 (1.37) |
| **CC1/ 2** | 1 (0.614) |
| **Reflections used in refinement** | 27669 (2708) |
| **Reflections used for R-free** | 1372 (112) |
| **R-work** | 0.2102 (0.4087) |
| **R-free** | 0.2505 (0.5051) |
| **CC (work)** | 0.942 (0.758) |
| **CC (free)** | 0.972 (0.631) |
| **Number of non-hydrogen atoms** | 4844 |
| **Macromolecule atoms** | 4543 |
| **Protein residues** | 567 |
| **RMS bonds (Å)** | 0.011 |
| **RMS angles (°)** | 1.33 |
| **Ramachandran favored (%)** | 94.23 |
| **Ramachandran allowed (%)** | 5.77 |
| **Ramachandran outliers (%)** | 0 |
| **Rotamer outliers (%)** | 0.39 |
| **Clashscore** | 21.9 |
| **Average B-factor ($Å^2$)** | 99.08 |
| **B-factor macromolecules ($Å^2$)** | 97.52 |

**Supplementary Table S5. Absolute molecular weight of SUMO-NrdR$_2^{ECO}$ and its mutant derivatives E36A, E42A, Y131A, and Δ132-149, in the absence and presence of nucleotides, determined by SEC-MALS.**

See SEC-MALS experiments on Figure 6. Mn, number-average molar mass; Mw, weight-average molar mass; Est. min/max, estimated minimum or maximum weight-average molar mass; PDI, polydispersity index (Mw/Mn); Wt%, percentage of protein elution mass in the corresponding peak; peaks with Wt% higher than 40% are highlighted in bold, peaks below 3% are unlisted. All molar mass values are listed in kDa. Error is listed a ± standard deviation.

| SEC-MALS peak | | | Quantification | | | | | | |
|---|---|---|---|---|---|---|---|---|---|
| Protein | Co-factor | Peak | Mn | Mw | Est.min | Est.max | PDI | Wt% | Interpretation |
| WT | - | 1 | 135.24 ± 1.29 | 136.31 ± 1.30 | 121.48 | 172.34 | 1.01 ± 0.01 | 6.88 | Tetramer →(+) |
| | | 2 | 72.95 ± 0.69 | 72.99 ± 0.69 | 69.12 | 75.63 | 1.00 ± 0.01 | 81.20 | **Dimer** |
| | | 3 | 42.06 ± 0.45 | 42.16 ± 0.45 | 39.84 | 46.78 | 1.00 ± 0.02 | 11.22 | Monomer →(+) |
| | ADP | 1 | 97.23 ± 0.19 | 97.40 ± 0.19 | 90.50 | 105.09 | 1.00 ± 0.00 | 38.60 | (Trimer) →(+) |
| | | 2 | 69.65 ± 0.32 | 70.65 ± 0.30 | 57.52 | 84.14 | 1.01 ± 0.01 | 61.40 | **Dimer** |
| | dATP | 1 | 175.55 ± 2.38 | 176.80 ± 2.44 | 156.90 | 242.66 | 1.01 ± 0.02 | 50.65 | **Tetramer →Octamer** |
| | | 2 | 139.27 ± 2.79 | 139.54 ± 2.77 | 127.20 | 149.84 | 1.00 ± 0.03 | 47.36 | **Tetramer** |
| | ATP | 1 | 133.05 ± 1.72 | 145.84 ± 1.82 | 70.09 | 213.23 | 1.10 ± 0.02 | 51.52 | **Dimer →Octamer** |
| | | 2 | 273.13 ± 3.17 | 275.41 ± 3.19 | 230.50 | 352.86 | 1.01 ± 0.02 | 44.02 | **Octamer →12mer** |
| E36A | - | 1 | 140.34 ± 2.93 | 141.25 ± 2.95 | 125.43 | 172.34 | 1.01 ± 0.03 | 5.09 | Tetramer →(+) |
| | | 2 | 59.24 ± 0.61 | 59.27 ± 0.61 | 56.47 | 60.94 | 1.00 ± 0.01 | 60.16 | **Dimer** |
| | | 3 | 50.45 ± 0.90 | 50.58 ± 0.88 | 44.77 | 54.05 | 1.00 ± 0.03 | 34.75 | Monomer →(+) |
| | dATP | 1 | 91.91 ± 0.11 | 92.02 ± 0.11 | 86.03 | 101.70 | 1.00 ± 0.00 | 48.14 | **(Trimer) →(+)** |
| | | 2 | 67.22 ± 0.17 | 68.38 ± 0.16 | 51.08 | 81.54 | 1.02 ± 0.00 | 51.49 | **Dimer** |
| | ATP | 1 | 102.70 ± 1.11 | 102.84 ± 1.11 | 95.36 | 108.33 | 1.00 ± 0.02 | 54.91 | **(Trimer) →(+)** |
| | | 2 | 78.58 ± 1.00 | 79.33 ± 1.00 | 64.38 | 90.60 | 1.01 ± 0.02 | 40.70 | **Dimer** |
| E42A | - | 1 | 108.85 ± 2.33 | 108.93 ± 2.33 | 104.77 | 114.80 | 1.00 ± 0.03 | 3.72 | (Trimer) →(+) |
| | | 2 | 67.25 ± 1.28 | 67.29 ± 1.28 | 65.28 | 72.19 | 1.00 ± 0.03 | 11.15 | Dimer |
| | | 3 | 31.39 ± 0.43 | 31.39 ± 0.43 | 31.08 | 32.13 | 1.00 ± 0.02 | 85.13 | **Monomer** |
| | dATP | 1 | 151.27 ± 0.41 | 166.02 ± 0.47 | 104.64 | 147.85 | 1.10 ± 0.00 | 20.27 | (Trimer) →(+) |
| | | 2 | 68.70 ± 0.15 | 68.87 ± 0.15 | 64.45 | 69.18 | 1.00 ± 0.00 | 15.43 | Dimer |
| | | 3 | 33.19 ± 0.07 | 33.21 ± 0.07 | 31.79 | 34.52 | 1.00 ± 0.00 | 63.37 | **Monomer** |
| | ATP | 1 | 72.38 ± 0.99 | 72.61 ± 0.99 | 67.27 | 83.12 | 1.00 ± 0.02 | 16.54 | Dimer →(+) |
| | | 2 | 36.85 ± 0.47 | 36.88 ± 0.47 | 34.85 | 37.83 | 1.00 ± 0.02 | 83.46 | **Monomer** |
| Y131A | - | 1 | 61.12 ± 0.05 | 61.12 ± 0.05 | 60.00 | 62.03 | 1.00 ± 0.00 | 47.86 | **Dimer** |
| | | 2 | 53.99 ± 0.11 | 54.13 ± 0.11 | 47.86 | 58.16 | 1.00 ± 0.00 | 42.17 | **Monomer →Dimer** |
| | | 3 | 39.93 ± 0.12 | 39.97 ± 0.12 | 38.68 | 43.23 | 1.00 ± 0.00 | 7.27 | Monomer |
| | dATP | 1 | 79.57 ± 0.11 | 79.62 ± 0.11 | 75.36 | 84.13 | 1.00 ± 0.00 | 44.14 | **Dimer →(+)** |
| | | 2 | 59.00 ± 0.16 | 59.90 ± 0.15 | 44.37 | 70.74 | 1.02 ± 0.00 | 55.86 | **Monomer →Dimer** |
| | ATP | 1 | 186.66 ± 0.26 | 189.95 ± 0.28 | 161.47 | 263.17 | 1.02 ± 0.00 | 3.01 | Tetramer →Octamer |
| | | 2 | 96.69 ± 0.07 | 96.78 ± 0.07 | 89.87 | 100.28 | 1.00 ± 0.00 | 44.45 | **(Trimer) →(+)** |
| | | 3 | 68.52 ± 0.08 | 69.79 ± 0.08 | 52.18 | 84.35 | 1.02 ± 0.00 | 52.54 | **Dimer →(Trimer)** |
| Δ(132-149) | - | 1 | 55.06 ± 0.61 | 55.08 ± 0.61 | 52.64 | 57.78 | 1.00 ± 0.02 | 72.76 | **Dimer** |
| | | 2 | 32.50 ± 0.41 | 32.52 ± 0.41 | 31.83 | 34.80 | 1.00 ± 0.02 | 27.24 | Monomer |
| | dATP | 1 | 65.84 ± 0.08 | 65.87 ± 0.08 | 62.19 | 67.53 | 1.00 ± 0.00 | 52.24 | **Dimer** |
| | | 2 | 50.40 ± 0.14 | 51.39 ± 0.15 | 34.87 | 59.68 | 1.02 ± 0.00 | 47.07 | **Monomer →Dimer** |
| | ATP | 1 | 59.17 ± 0.70 | 59.24 ± 0.70 | 54.92 | 61.36 | 1.00 ± 0.02 | 78.87 | **Dimer** |
| | | 2 | 31.44 ± 0.69 | 31.44 ± 0.67 | 31.16 | 32.35 | 1.00 ± 0.03 | 18.03 | Monomer |

**Supplementary Table S6. Strains and plasmids in this study.**

Strains and plasmids may also be referred to by simplified names (Alias). Strain genotypes are annotated with the following conventions: changes relative to a parent strain are listed as (Parent / changes); phenotypic notes are indicated between curly brackets (e.g., {restriction, methylation}); square brackets indicate material incorporated in a prophage or natural plasmid (e.g., λ[content]); a capital P indicates a promoter (e.g., P*nrdA*); antibiotic resistance is listed after the genotype with the following abbreviations: Amp (ampicillin), Kn (kanamycin), Tc (tetracycline), Str (streptomycin), and Gm (gentamicin). The origin of *nrdR* genes or RNR promoters is disclosed as *E. coli* K-12 *substr*. MG1655 ($^{ECO}$) or *P. aeruginosa* PAO1 ($^{PAO}$).

| ***List of strains*** | | | | |
|---|---|---|---|---|
| **Strain** | **Alias** | **Genotype, Resistance** | **Description** | **Source** |
| PAO1 | PAO1 | Wild type (ATCC 15692 / CECT4122) | *P. aeruginosa*, wild type lab strain | ATCC 15692 |
| PW7855 | PAO1 Δ*nrdR* | PAO1 / *nrdR*::IS[*lacZ*]/hah, **Tc$^{R}$** | *P. aeruginosa*, PAO1 with *nrdR* interrupt. | Jacobs *et al.* [1] |
| DH5α | DH5α | $F^{-}$ $λ^{-}$ *endA1 recA1 relA1 gyrA96 glnV44 deoR nupG hsdR17* {$r_{K-}m_{K+}$} Δ(*lacZYA-argF*)U169 *purB20 thi*$^{-}$1 φ80d[*lacZ*ΔM15] | *E. coli*, lab strain for cloning procedures | Lucigen |
| K-12 (Migula) | K12 MG1655 | $F^{-}$ $λ^{-}$ ilvG$^{-}$ rfb$^{-}$*50* rph$^{-}$1 | *E. coli*, wild type lab strain | ATCC 700926 |
| ETS106 | K12 Δ*nrdR* | K-12 / Δ*nrdR*::*kan*, **Kn$^{R}$** | *E. coli* K-12 carrying a full *nrdR* deletion | Cendra *et al.* [2] |
| BL21(DE3) | BL21 | B / $F^{-}$ λDE3[*lacI PlacUV5–T7p07 ind1 sam7 nin5*] *ompT dcm lon hsdS$_{B}$* {$r_{B-}m_{B-}$} *malB*$^{+K-12}${$λ_{S}$} | *E. coli* strain for IPTG-induced protein overexpression (NrdR-$H_6$, NrdR$_2$) | Lucigen |
| Biotin Xcell F' | MC1061*bir* | MC1061 / F'[*proAB$^{+}$ lacI* qZΔM15::Tn10] *araD139* Δ(*ara,leu*)7696 Δ*lac* I74 *galU galK mcrB1 rpsL hsdR2* {$r_{K-}m_{K+}$} *ara* P$_{BAD}$-*birA*, **St$^{R}$ Tc$^{R}$** | *E. coli* strain for IPTG-induced protein overexpression (NrdR$_1$) | Lucigen |

| ***List of plasmids*** | | | |
|---|---|---|---|
| **Plasmid** | **Alias** | **Description, Resistance** | **Source** |
| pJET1.2-blunt | pJET1.2 | General carrier vector for cloning procedures, **Amp$^{R}$** | ThermoFisher |
| pET22b$^{+}$ | pET22b+ | Vector for IPTG-induced T7 protein overexpression of His$_6$-fusion proteins, **Amp$^{R}$** | Novagen |
| pET22b$^{+}$::*nrdR*$^{PAO}$ | pET-NrdR(PAO) | pET22b+ derivative producing an NrdR-His6 (NrdR$^{PAO}$-$H_6$) fusion protein, **Amp$^{R}$** | This work |
| pAviTag-NN-His SUMO Kan | pSUMO | Vector for Rhamnose-induced overexpression of SUMO-fusion proteins, **Kn$^{R}$** | Lucigen |
| pAviTag-NN-His SUMO Kan::*nrdR*$^{PAO}$ | pSUMO-NrdR(PAO) | pSUMO derivative producing a His$_6$-AviTag-SUMO-NrdR fusion protein (NrdR$_1$$^{PAO}$), **Kn$^{R}$** | This work |
| pAviTag-NN-His SUMO Kan::*nrdR*$^{ECO}$ | pSUMO-NrdR(ECO) | pSUMO derivative producing a His$_6$-AviTag-SUMO-NrdR (NrdR$_1$$^{ECO}$) fusion protein, **Kn$^{R}$** | This work |
| pCri11a | pCri11a | Vector for IPTG-induced overexpression of His$_6$-SUMO-His$_6$ fusion proteins, **Kn$^{R}$** | Goulas *et al.* [3] |
| pCri11a::*nrdR*$^{PAO}$ | pCri-NrdR(PAO) | pCri derivative producing a His$_6$-SUMO-TEVcs-NrdR (NrdR$_2$$^{PAO}$) fusion protein, **Kn$^{R}$** | This work |
| pCri11a::*nrdR*$^{ECO}$ | pCri-NrdR(ECO) | pCri derivative producing a His$_6$-SUMO-TEVcs-NrdR (NrdR$_2$$^{ECO}$) fusion protein, **Kn$^{R}$** | This work |
| pETS130-GFP | pETS130 | Broad-host range, promoterless GFP, Gm$^{R}$, **Gm$^{R}$** | Sjoberg *et al.* [4] |
| pETS150 | pETS130-PA | pETS130 derivative, GFP controlled by the promoter region P*nrdAB*$^{ECO}$, **Gm$^{R}$** | Cendra *et al.* [2] |
| pReViTA-0 | pReViTA | pETS130 derivative, *in vitro* transcription template plasmid for ReViTA, **Gm$^{R}$** | This work |
| pReViTA::P*nrdD*$^{ECO}$ | pReViTA-PD | pReViTA derivative, carrying the promoter region P*nrdDG*$^{ECO}$, **Gm$^{R}$** | This work |
| pETS176 | pUCP20T::nrdR | pUC2P0T derivative, complementation plasmid for *nrdR*, **Amp$^{R}$** | Crespo *et al.* [5] |
| pCri11a::*nrdR*$^{ECO}$E36A | pCri-NrdR-E36A | pCri-NrdR(ECO) derivative producing a mutant NrdR$_2$$^{ECO}$E36A fusion protein, **Kn$^{R}$** | This work |
| pCri11a::*nrdR*$^{ECO}$E42A | pCri-NrdR-E42A | pCri-NrdR(ECO) derivative producing a mutant NrdR$_2$$^{ECO}$E42A fusion protein, **Kn$^{R}$** | This work |
| pCri11a::*nrdR*$^{ECO}$ Y131A | pCri-NrdR-Y131A | pCri-NrdR(ECO) derivative producing a mutant NrdR$_2$$^{ECO}$Y131A fusion protein, **Kn$^{R}$** | This work |
| pCri11a::*nrdR*$^{ECO}$Δ(132-149) | pCri-NrdRΔ132 | pCri-NrdR(ECO) derivative producing a mutant NrdR$_2$$^{ECO}$Δ(132-149) fusion protein, **Kn$^{R}$** | This work |
| pLG338/30 | pLG338/30 | Low copy number plasmid for complementation assays in *E. coli*, **Amp$^{R}$ Tc$^{R}$** | GenScript [6] |
| pLG338/30::*nrdR*$^{ECO}$ | pLG338-NrdR | pLG338 derivative expressing wild type *nrdR*$^{ECO}$, **Amp$^{R}$ Tc$^{R}$** | GenScript |
| pLG338/30::*nrdR*$^{ECO}$E36A | pLG338-E36A | pLG338 derivative expressing mutant *nrdR*$^{ECO}$E36A, **Amp$^{R}$ Tc$^{R}$** | GenScript |
| pLG338/30::*nrdR*$^{ECO}$ E42A | pLG338-E42A | pLG338 derivative expressing mutant *nrdR*$^{ECO}$E42A, **Amp$^{R}$ Tc$^{R}$** | GenScript |
| pLG338/30::*nrdR*$^{ECO}$Y131A | pLG338-Y131A | pLG338 derivative expressing mutant *nrdR*$^{ECO}$Y131A, **Amp$^{R}$ Tc$^{R}$** | GenScript |

### Supplementary Table S7. Sequence and application of the primers used in this study

Primers are commonly referred to in the text by their numbers as listed here. [D3-PA] in 28 indicates D3-phosphoramidite.

| Number | Name | Sequence | Application |
|---|---|---|---|
| 1 | NrdR_NdeI_fw | CATATGCATTGTCCCTTCTGCGGTG | Cloning, pET-NrdR(PAO) |
| 2 | NrdR_XhoI_rv | CTCGAGTTCCTTGGCCGGCTCGCG | Cloning, pET-NrdR(PAO) |
| 3 | T7-promoter_fw | TAATACGACTCACTATAGGG | PCR test, pET-NrdR / pCri-NrdR |
| 4 | T7-terminator_rv | CTAGTTATTGCTCAGCGGTG | PCR test, pET-NrdR / pCri-NrdR |
| 5 | NrdR-PAO_SUMO_fw | CGCGAACAGATTGGAGGTGGCAGCATGCATTGTCCCTTCTGCGGT | Cloning, pSUMO-NrdR(PAO) |
| 6 | NrdR-PAO_SUMO_rv | GTGGCGGCCGCTCTATTACGTTCATTCCTTGGCCGGCTC | Cloning, pSUMO-NrdR(PAO) |
| 7 | NrdR-ECO_SUMO_fw | CGCGAACAGATTGGAGGTGGATCCATGCATTGCCCATTCTGTTTC[illegible] | Cloning, pSUMO-NrdR(ECO) |
| 8 | NrdR-ECO_SUMO_rv | GTGGCGGCCGCTCTATTAGGCTTAGTCCTCCAGGCGC | Cloning, pSUMO-NRdR(ECO) |
| 9 | SUMO_fw | ATTCAAGCTGATCAGACCCCTGAA | PCR test, pSUMO-NrdR |
| 10 | pETite_rv | CTCAAGACCCGTTTAGAGGC | PCR test, pSUMO-NrdR |
| 11 | NrdR-PAO_TEV_fw | ATTACCATGGGCGAGAACCTTTACTTTCAAGGCAGCGGCAGCGGC AGCATGCATTGTCCCTTCTGC | Cloning, pCri-NrdR(PAO) |
| 12 | NrdR-PAO_TEV_rv | TATATACTCGAGTCATTCCTTGGCCGGCTCGCG | Cloning, pCri-NrdR(PAO) |
| 13 | NrdR-ECO_TEV_fw | TATACCATGGGCGAGAACCTTTACTTTCAAGGATCCGGATCCGGA TCCATGCATTGCCCATTCTGTTTCGC | Cloning, pCri-NrdR(ECO) |
| 14 | NrdR-ECO_TEV_rv | TATATACTCGAGTTAGTCCTCCAGGCGCGCGATCT | Cloning, pCri-NrdR(ECO) |
| 15 | pETS130-backB_fw | AATCTAGATGCCCATGGACGCACAC | Cloning, pReViTA |
| 16 | pETS130-backB_rv | AAGACGTCCGGGGAGGCAGACAAGGTATA | Cloning, pReViTA |
| 17 | PnrdD-ECO_BamHI_fw | AAAGGATCCTTGAGGCTGTCTGGTGGTTAC | Cloning, pReViTA-PD / EMSA probes |
| 18 | PnrdD-ECO_ClaI_rv | AAAATCGATGCACTTTGCAGCCGTCTCG | Cloning, pReViTA-PD / EMSA probes |
| 19 | ReViTA_TEST_fw | AGCACAAGTTTTATCCGGCC | ReViTA, qPCR |
| 20 | ReViTA_TEST_rv | CATATCACCAGCTCACCGTC | ReViTA, qPCR |
| 21 | ReViTA_TEST_rt | TGCTCATGGAAAACGGTGTAAC | ReViTA, rev. transcription |
| 22 | ReViTA_CTRL_fw | TTTCGGTCGTGAGTTCGGAG | ReViTA, qPCR |
| 23 | ReViTA_CTRL_rv | GCAAGCGCGATGAATGTCTT | ReViTA, qPCR |
| 24 | ReViTA_CTRL_rt | CGCCAACAACCGCTTCTTG | ReViTA, rev. transcription |
| 25 | PnrdA-ECO_m13_rv | CTGGGCGTCGTTTTACAAACTGAATGTGGGAGCG | EMSA probes |
| 26 | Anr-ctrlNEG_fw | GAATTCATGGCCGAAACCATCAAG | EMSA probes |
| 27 | Anr-ctrlNEG_m13_rv | CTGGGCGTCGTTTTACCTTCTTCGACAGCAGCAG | EMSA probes |
| 28 | WellRed M13 | [D3-PA]GTCACTGGGCGTCGTTTTAC | EMSA probes, infrared dye |
| 29 | NrdR-ECO-E36A_fw | CTGGTGTGTAATGCACGTTTCACCACC | E36A mutation, NrdR(ECO) |
| 30 | NrdR-ECO-E36A_rv | GGTGGTGAAACGTGCATTACACACCAG | E36A mutation, NrdR(ECO) |
| 31 | NrdR-ECO-E42A_fw | TTCACCACCTTTGCAGTGGCGGAGCTG | E42A mutation, NrdR(ECO) |
| 32 | NrdR-ECO-E42A_rv | CAGCTCCGCCACTGCAAAGGTGGTGAA | E42A mutation, NrdR(ECO) |
| 33 | NrdR-ECO-Y131A_fw | TGCCTCTGTCGCCCGCAGTTTCGAA | Y131A mutation, NrdR(ECO) |
| 34 | NrdR-ECO-Y131A_rv | GAAACTGCGGGCGACAGAGGCAAA | Y131A mutation, NrdR(ECO) |
| 35 | NrdR-ECO-132149_fw | TATACCATGGGCGAGAACCTTTACTTTCAA | 132-149 deletion, NrdR(ECO) |
| 36 | NrdR-ECO-132149_rv | ATACTCGAGTTAGTAGACAGAGGCAAAACGGATATAGGCGAC | 132-149 deletion, NrdR(ECO) |
